# Cas9 Nickase-Mediated Contraction of CAG/CTG Repeats *in Vivo* is Accompanied by Improvements in Huntington’s Disease Pathology

**DOI:** 10.1101/2024.02.19.580669

**Authors:** Alvaro Murillo, Melanie Alpaugh, Ruban Rex Peter Durairaj, Emma L. Randall, Meghan Larin, Laura Heraty, Alys N. Aston, Alysha S. Taylor, Alex Mas Monteys, Nina Stöberl, Aeverie E. R. Heuchan, Pascale Aeschlimann, Kyle Fears, Christian Landles, Georgina F Osborne, Antoine Mangin, Soumyasree Bhattacharyya, Emmanouil Metzakopian, Nicholas D. Allen, Jack Puymirat, Gillian P Bates, Beverly L. Davidson, Francesca Cicchetti, Mariah J. Lelos, Vincent Dion

## Abstract

Expanded CAG/CTG repeats cause over 15 different diseases that all remain without a disease-modifying treatment. Because repeat length accounts for most of the variation in disease severity, contracting them presents an attractive therapeutic avenue. Here, we show that the CRISPR-Cas9 nickase targeted to CAG/CTG repeats leads to efficient contractions in Huntington’s disease patient-derived neurons and astrocytes, and in myotonic dystrophy type 1 patient-derived neurons. The approach is allele-selective and free of detectable off-target mutations. Striatal injection of the Cas9 nickase in a mouse model for Huntington’s disease using adeno-associated viral vectors led to contractions in over half the infected cells. Upon injection, we observed a reduction in the number of inclusion bodies, improved transcriptome, and ameliorated locomotion. The effects were greater than expected from the contractions induced and suggest that non-cell autonomous mechanisms may be involved. Our results provide the proof-of-concept that correction of CAG/CTG repeats can improve Huntington’s disease phenotypes *in vivo*.

**One sentence summary:** The Cas9 nickase contracts CAG/CTG repeats at multiple disease loci in patient-derived cells and improves molecular and behavioral phenotypes in HD.

## Introduction

The heterozygous expansion of CAG/CTG repeats at 15 different loci in the genome causes clinically distinct intractable neurodegenerative and neuromuscular disorders^1^, including Huntington’s disease (HD) and myotonic dystrophy type 1 (DM1)^2,3^. They are all currently without a disease modifying treatment^4,5^. The size of the expanded repeat tract explains up to 60% of the variation in the age at disease onset in HD^6^, with longer repeats leading to more severe phenotypes. This is compounded by repeat expansions accumulating over time in somatic tissues, especially in those cell types that are most affected by the disease^7–9^. Moreover, DNA repair factors involved in somatic expansion have been identified as disease modifiers^10–14^. These observations suggest that somatic expansion is a major driver of disease pathogenesis and that preventing expansions in relevant cell types, or, better, inducing contractions, may provide a much-needed therapeutic avenue.

Several studies have used gene editing *in vivo* at the *DMPK* and *HTT* loci, where repeat expansions cause DM1 and HD, respectively^15–25^. All these approaches mutate the repeat locus, either by excising and producing uncontrollable insertions, deletions, and rearrangements or by mutating the repeat tract using base editors. A small molecule has also been developed that induces contractions of the repeat^26^. Editing efficiencies ranged from 2% to 54% of the targeted cells. Molecular and behavioural rescues have not been consistently tested, but some studies reported partial rescues^15,22^.

Importantly, despite patients having a heterozygous expansion, typical gene editing approaches mutate both alleles, which may have undesirable consequences^4,27,28^. Three studies have circumvented this problem by designing sgRNAs that target a specific single-nucleotide polymorphism distinguishing the two *HTT* alleles^16,21,29^. However, this approach reduces the total number of individuals who stand to benefit since the most common variant is found in only 30% of HD individuals of European ancestry^29^.

Altogether, these results highlight the need to correct - rather than mutate - the expanded allele. Gene editing approaches need to be efficient, allele-selective, and applicable to as many individuals as possible, regardless of disease. Double-strand breaks, which are highly mutagenic, should be avoided to reduce the risk of off-target mutations. Importantly, behavioural and molecular pathology must be improved by the treatment.

We showed previously that the Cas9D10A nickase could induce contractions of ectopic CAG repeats when targeted directly to the repeat tract in immortalised cells using one single-guide RNA (sgRNA) in a transcription-dependent and replication-independent manner^30–32^. Here, we harnessed the Cas9D10A nickase to correct expanded CAG/CTG repeats and provide the *in vivo* proof-of-concept for the safe correction of expanded repeats as a therapeutic avenue.

## Results

### Nickase-induced contractions in patient-derived cells

We first investigated the possibility of contracting CAG/CTG repeats in disease-relevant cells. Astrocytes contribute considerably to HD pathogenesis^33–37^. We differentiated HD human induced pluripotent stem cells (iPSCs) (CS09iHD109-n1), harbouring 130 to 140 CAG/CTG repeats at the *HTT* locus^38^, into astrocyte progenitors and then to mature S100β^+^ astrocytes (Fig. 1a and Supplementary Fig. 1a). We transduced them with a lentivirus expressing Cas9D10A (Supplementary Table 1). A second lentivirus expressed the sgRNA against the repeat tract (sgCTG) and contained a GFP cassette to monitor silencing of the vector. We found that GFP and Cas9D10A expression was sustained over 42 days (Supplementary Fig. 1bc). We assessed repeat size distribution using small-pool PCR (Fig. 1b)^39^ and observed a 6-fold increase in the proportion of contracted alleles in the dual treated cultures. We also saw a >2-fold decrease in the number of large alleles (>134 CAGs) compared to populations transduced with Cas9D10A only (P≤0.0001, Fig. 1c). On average, the cells treated with both Cas9D10A and sgCTG had 20 fewer repeats than the cells treated with Cas9D10A only. These results suggest that HD iPSC S100β^+^ astrocytes undergo robust contractions when exposed to both the Cas9D10A and sgCTG.

**Figure 1.**
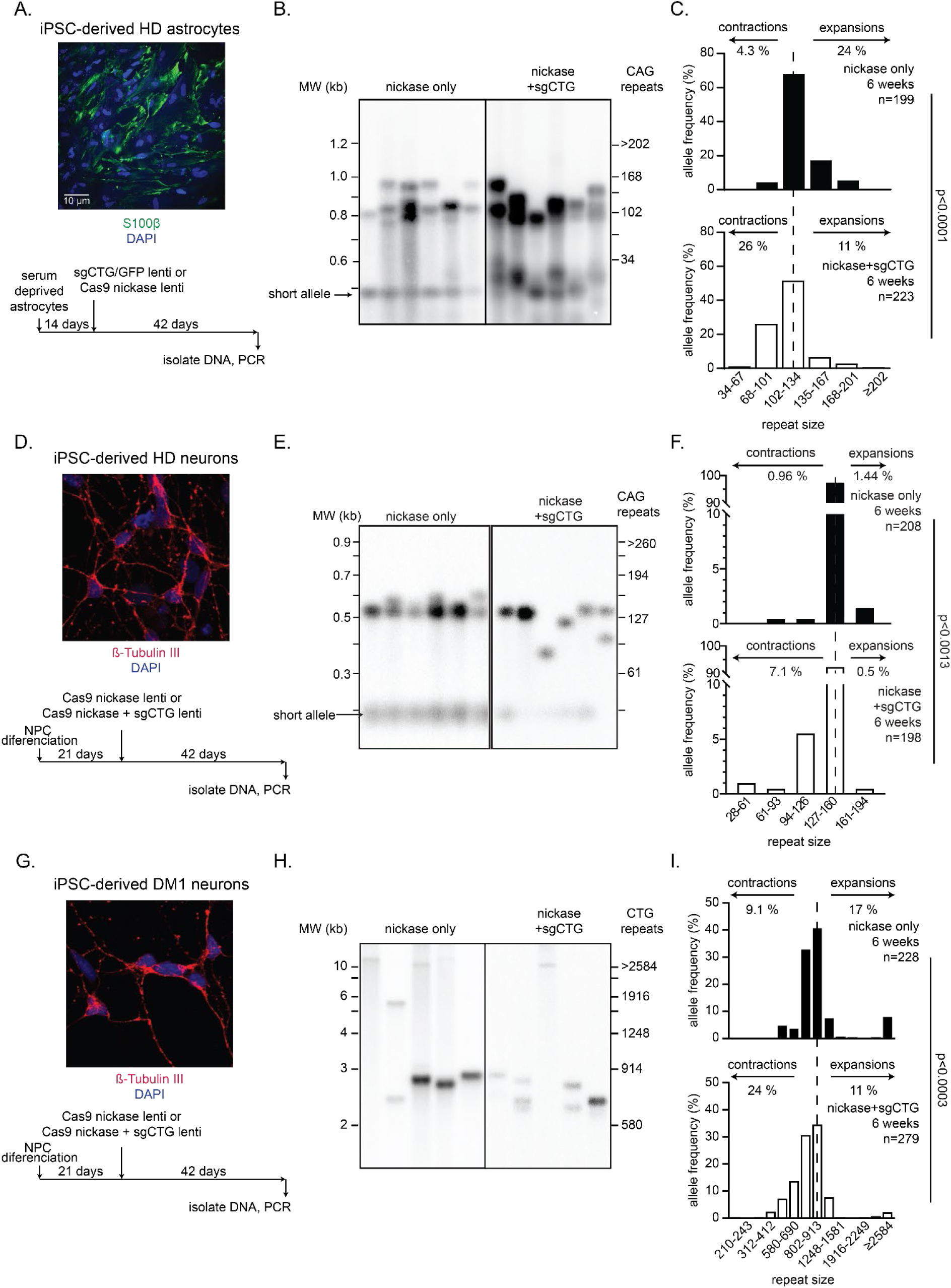
Cas9D10A-induced contractions in HD and DM1 iPSC-derived cells. A) Top: Representative confocal image of HD iPSC-derived astrocytes stained with S100β and DAPI. Bottom: timeline of the experiments. B) Representative small-pool PCR blot showing contractions in HD iPSC-derived astrocytes that were either transduced with the Cas9D10A only or both the Cas9D10A and the sgCTG for 42 days. C) Quantification of the small-pool PCR blots for HD iPSC-derived astrocytes. D) Top: Representative confocal image of HD iPSC-derived cortical neurons stained with β-Tubulin III and DAPI. Bottom: timeline of the experiments. E) Representative small-pool PCR blot showing contractions in HD iPSC-derived cortical neurons that were transduced with Cas9D10A only or both Cas9D10A and the sgCTG for 42 days. F) Quantification of the small-pool PCR blots for HD iPSC-derived cortical neurons. G) Top: Representative confocal image of DM1 iPSC-derived cortical neurons stained with β-Tubulin III and DAPI. Bottom: timeline of the experiments. H) Representative small-pool PCR blot showing contractions in DM1 iPSC-derived cortical neurons that were transduced with Cas9D10A only or both Cas9D10A and the sgCTG for 42 days. I) Quantification of the small-pool PCR blots for DM1 iPSC-derived cortical neurons. ‘n’ is the number of alleles counted in the small-pool PCR experiments. Scale bar = 10 μm. Dashed lines in panels C, F, and I indicate modal repeat size.

In HD, the driving factor of pathogenesis is neuronal death, which occurs first in the striatum with medium spiny neurons, followed by neurons in the cortex^40^. We differentiated the same HD iPSCs into neurons (Fig. 1d)^41^. Our cultures were 99% pure as judged by ꞵ-tubulin staining (Supplementary Fig. 1d). We exposed the neurons to the Cas9D10A and sgCTG for 21 or 42 days and performed small-pool PCRs (Supplementary Fig. 1ef, Fig. 1e). We found an increase in the frequency of alleles shorter than 127 CAGs over time, reaching a 7-fold increase after 42 days and a 3-fold decrease in the frequency of alleles with over 160 repeats (Fig. 1e-f, P≤0.0013). We confirmed the results using single-molecule real-time (SMRT) long read sequencing (Supplementary Fig. 2). These results suggest that the Cas9D10A can contract expanded CAG/CTG repeats in HD iPSC-derived neurons.

The advantage of our design is that the repeat tract itself is targeted and, therefore, it should induce contractions at multiple disease loci. Although DM1 patients display debilitating myotonia and muscle wasting, the brain is extensively affected, including the cortex^42^. We generated cortical neurons from an iPSC line that harbours an average of 996 CTGs at the *DMPK* locus, with some alleles exceeding 2500 CTGs (Supplementary Fig. 1g, Fig. 1gh). Although it was not sufficient to lead to repeat sizes in the non-pathogenic range, the cells treated with both Cas9D10A and sgCTG had 166 fewer repeats on average than the cells expressing only Cas9D10A (Fig. 1i). We conclude that Cas9D10A induces contractions in at least two distinct disease loci without changing the Cas9D10A construct or the sgRNA.

One important question is: what is the minimum expansion length that remains a substrate for Cas9D10A? We have shown previously that repeats of at least 101 CAG/CTGs contract readily to non-pathogenic sizes (i.e., 35 CAGs or lower for HD and below 50 for DM1) but that 42 repeats remain stable^30^. To determine the threshold with more precision, we used a HD patient-derived lymphoblastoid cell line that harbours 60 CAG repeats at the expanded *HTT* locus and delivered both Cas9D10A and sgCTG via lentiviruses (Supplementary Table 1). Forty-two to 56 days after transduction of the sgCTG virus, we isolated the cells and performed SMRT sequencing. We found that cells transduced with both viruses contracted readily (Supplementary Fig. 3), suggesting that the threshold for contraction with Cas9D10A is between 42 and 60 repeats.

### Off-target mutations remain at background levels in the presence of both Cas9D10A and sgCTG

An important safety concern in the context of gene editing is the generation of off-target mutations^43^. Most approaches mitigate this issue by carefully designing the sgRNA and adjusting the target sequence accordingly. Here, this is not possible as we target the repeat tract itself. To address whether our approach led to unwanted off-target mutations, we first used PCR-free whole genome sequencing using Illumina short-read sequencing. We used the same HD iPSC-derived neurons cultured without transduction for 42 days and compared them with cultures that were transduced with either Cas9D10A only or Cas9D10A and the sgCTG. We obtained at least 5.5×10^9^ reads per treatment, including >3.9×10^8^ reads per condition spanning 1830 genes that contain a targetable CAG/CTG repeat (see methods, Supplementary Table 2). The frequency of mutations was not different between the transduced and the non-transduced cells (P>0.31, Table 1, Supplementary Fig. 4). Similar results were found using the HD iPSC-derived astrocytes (P>0.41, Supplementary Figure 4, Table 1). In total, we found 8 mutations overlapping with CAG/CTG repeats, but they were not enriched in the samples treated with Cas9D10A and sgCTG (Fisher’s exact test P>0.26, Supplementary Table 2). These results suggest that the expression of Cas9D10A together with sgCTG did not induce mutations above the background seen in untreated cells.

**Table 1.** Analysis of mutations in HD iPSC-derived cortical neurons and astrocytes.

|  |  | CAG/CTG repeat genes |  |  | Whole Genome |  |  |
| --- | --- | --- | --- | --- | --- | --- | --- |
| Cell type | Treatment | # of mutations* | # of reads x10<sup>8</sup> | P-value<sup>§</sup> | # of mutations* | # of reads x10<sup>9</sup> | P-value<sup>§</sup> |
| Neurons | - | 51±15 | 4.06 | - | 1558±260 | 5.6 | - |
|  | Cas9D10A | 60±18 | 4.17 | 0.52 | 1806±396 | 5.7 | 0.23 |
|  | Cas9D10A + sgCTG | 39±15 | 3.98 | 0.31 | 1512±261 | 5.5 | 0.99 |
| Astrocytes | Cas9D10A | 26±22 | 1.53 | 0.41 | 1819±793 | 2.1 | 0.52 |
|  | Cas9D10A + sgCTG | 13±4 | 1.33 |  | 1361±308 | 1.9 |  |

\*: The number of mutations obtained per sample (8 samples/treatment for the neuronal cultures and 3 samples/treatment for the astrocytic cultures). No difference was seen between 21 and 42 days in astrocytes and therefore the data were combined for this analysis.
§: The number of mutations found at the selected 1830 genes and genome-wide were normalised to the total reads per sample and indicated as the number of mutations normalised to the number of reads (Supplementary Figure 4). Differences in normalised mutations between neuronal cultures conditions were calculated by one-way ANOVA with a post-hoc Tukey's multiple comparison test, except for the whole genome data from astrocytes where we used a Student's t-test.

Second, we looked for rearrangements at the non-expanded *HTT* allele 42 days after treatment in HD iPSC-derived neuronal cultures using the targeted long-read sequencing data presented above (Supplementary Fig. 2). We used Repeat Detector^44^ to determine repeat size and Sniffles2^45^ to detect rearrangements in this dataset. These analyses suggested that one sample transduced with Cas9D10A and sgCTG had an insertion downstream of the repeat tract at a frequency of 0.107. However, we could not identify this rearrangement with the Integrative Genomics Viewer^46^ or by manually looking for the rearranged alleles in the circular consensus sequences. Moreover, we found no rearrangements in the other 3 samples treated with both Cas9D10A and sgCTG, which came from the same neuronal epithelium, but were differentiated separately from neuronal progenitor cells. Thus, rearranged alleles were not detectable in this sample. We conclude that this was a false negative and that mutations are rare, if present at all, in the non-expanded alleles upon a 42-day exposure to Cas9D10A and sgCTG.

### Expression of AAV-delivered GFP and Cas9D10A in vivo

Next, we established whether we could deliver Cas9D10A and sgCTG directly to the striatum of mice using recombinant adeno-associated viral vectors (AAV) serotype 9. The whole CRISPR system is too large to be packaged into a single AAV and therefore we used a two-vector system with one AAV harbouring Cas9D10A and the other expressing both sgCTG and GFP (Fig. 2ab, Supplementary Table 3).

When injected in R6/1 mice two days after birth (P2), GFP was detectable throughout the striatum along with some expression in the cortex 6 weeks after injection (Supplementary Fig. 5a). Expression was still present 3 months post-injection without a noticeable drop in expression over time (Supplementary Fig. 5b). AAV9 exhibited strong tropism to NeuN^+^ neurons and expressed GFP well (Supplementary Fig. 5cd), consistent with previous work^47^. Furthermore, using qPCR, we found that there were 26 ± 6 copies of the sgCTG/GFP AAV genome per cell in striata, on average, one month after injection (Supplementary Fig. 5e). These data demonstrate that AAV9 can infect striatal neurons in R6/1 mice and that GFP is robustly expressed over several months.

Cas9D10A expression, however, proved more challenging. We tested four different AAV constructs for Cas9 expression (Supplementary Table 3) that were previously shown to induce edits *in vivo*^21,48,49^ and introduced the D10A mutation. Of the four constructs injected in wild type and R6/1 mice, one (version 1 - v1) did not express any detectable Cas9D10A, v2 and v3 were expressed 2 weeks post-injection but silenced later (Supplementary Fig. 6a-f). The last construct, v4, containing a miniCMV promoter, was still expressed 2 months after injection in adults, albeit at lower levels (Supplementary Fig. 6g). When we injected P2 mice, we could detect Cas9D10A for up to 5 months, but expression decreased over time (Supplementary Fig. 6gh). These data suggest that Cas9D10A expression wanes over time at a rate heavily dependent on the construct used. We continued with Cas9D10A v4 to test whether the nickase can induce contractions *in vivo*.

### Large contractions after delivery of Cas9D10A and sgCTG to neonatal mice

Next, we aimed to determine whether Cas9D10A could induce contractions *in vivo*. Contractions in the mouse brain are measured against an ongoing rate of somatic expansion. SMRT sequencing of the expanded repeats showed, as expected, that there was a small but progressive increase in the modal repeat size over time, accompanied by a bimodality beyond 10 weeks of age (Supplementary Fig. 2e). This is due to different cell types having different rates of expansion, with the medium spiny neurons having the largest repeat sizes^50^. The average copy number of Cas9D10A 1 month after injection of R6/1 mice at P2 was 0.3 ± 0.15 copies per cell on average (Supplementary Fig. 6i). Therefore, the efficiency of contractions is capped at 26% on average in these experiments, as predicted by a Poisson distribution.

Repeat size in P2-injected mice showed fewer expansions in the expanding medium spiny neurons as well as an accumulation of contractions over time (Fig. 2d-h). Specifically, short alleles of 12 to 17 CAGs, which are well within the non-pathogenic range, appeared 5 months post-injection (Supplementary Fig. 7a-f; Fig. 2d-h). These large contractions were seen in all 3 mice sequenced at this late time point. The instability index^51^ showed a slower increase over time in the hemisphere injected with both Cas9D10A and sgCTG compared to Cas9D10A alone (Supplementary Fig. 8a, P=0.0004), which was accompanied by a more negative contraction index (Supplementary Fig. 8b, P=0.0002), and a lower expansion index (Supplementary Fig. 8c, P=0.005). We further found that the average repeat size was reduced compared to mice injected only with Cas9D10A, especially after 4 and 5 months (Supplementary Fig. 8d, P<0.001). We observed a rise in the percentage of short alleles in the samples treated with both Cas9D10A and sgCTG from 5.1 ± 1.5% 1-month post-injection to 15.7 ± 7.6% 5 months after the injections (Supplementary Fig. 8d, Fig. 2i, P=0.035). Given that we have not sorted the infected cells, the expected maximum was 26%. We conclude that most cells that express Cas9D10A together with sgCTG in the striatum of R6/1 mice harbor contractions after 5 months.

**Figure 2.**
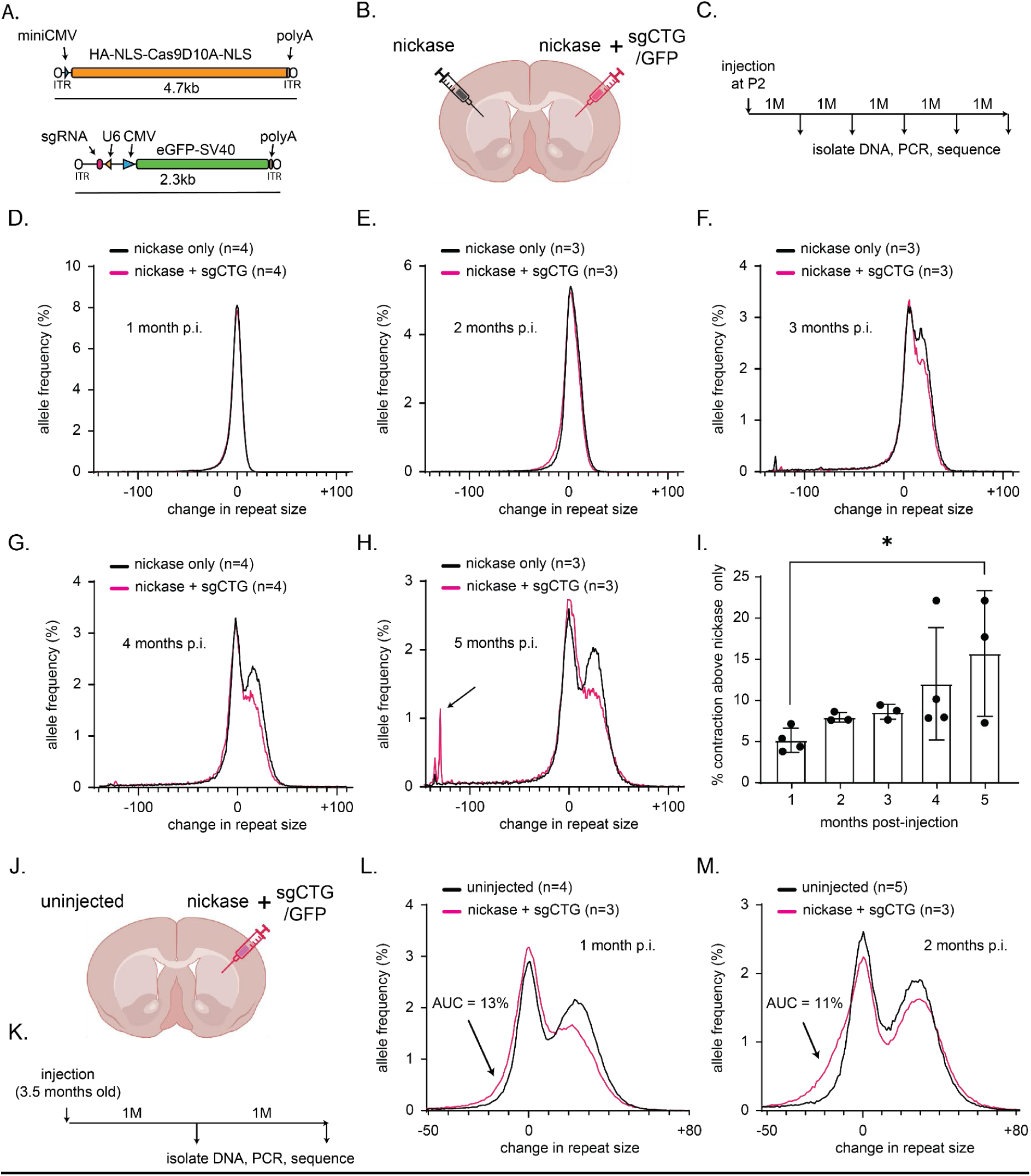
CAG repeat contraction in P2- and adult injected R6/1 animals. A) Schematic of the AAV constructs used. B) Schematic of the experimental approach. C) Timeline of the experiment. P2 = 2 days after birth. D) Aggregated graphs of repeat size distribution in hemispheres of R6/1 mice injected with Cas9D10A only (black) versus Cas9D10A + sgCTG (magenta) 1 month, E) 2 months, F) 3 months, G) 4 months and H) 5 months post-injection. Arrow points to large contractions with 17 remaining CAGs. Note that the graphs are normalised to the modal peak size found in each hemisphere (see methods). The mice used in these experiments had a modal repeat size ranging from 138 to 151 CAGs. A nonparametric Wilcoxon matched-pairs signed rank test showed no significant difference in repeat size in animals sacrificed 1 to 3 months post-injection. After 4 and 5-months, the difference was significant (P value <0.0001 (****)). I) Difference in the area under the curve between the Cas9D10A + sgCTG treatment and Cas9D10A alone (*P<0.05 using a one-way ANOVA). Error bars represent ± standard deviation. Number of animals: Cas9D10A only; 1 month n=4, 2 months n=3; 3 months n=3; 4 months n=4; 5 months n=3; Cas9D10A + sgCTG; 1 month n=4, 2 months n=3; 3 months n=3; 4 months n=4; 5 months n=3). J) Schematic of the experimental approach for injections in adult mice. K) Timeline of the experiments. L) Aggregated graphs of repeat size distribution in the striatum of uninjected adult R6/1 mice (black) compared to striata injected with Cas9D10A + sgCTG (magenta) 1 month post-injection. Unpaired, nonparametric Kolmogorov-Smirnov test returned a significant difference in cumulative distributions. (P <0.023 (*)). M) Same as L, but 2 months after injection ( P<0.0001 (****)). The mice used in these experiments had a modal repeat size ranging from 138 to 145 CAGs. Number of animals: uninjected 1-month post-injection n=4, 2 months post-injection n=5; Cas9D10A + sgCTG; 1 month post-injection n=3, 2 months post-injection n=3.

### Contractions in the R6/1 adult striatum upon Cas9D10A and sgCTG delivery

Next, we asked whether contractions also occurred in adult mice upon treatment with the Cas9D10A. We repeated the experiments, injecting 3.5 months old adult R6/1 mice with both Cas9D10A and sgCTG and comparing them with age-matched un-injected controls. We tracked the instability at 1 and 2 months post-injection (Fig. 2j-m). We found robust infection efficiencies with an average of 464 ± 217 and 663 ± 92 AAV DNA copies per cell when injecting AAV-sgCTG-GFP and AAV-Cas9D10A, respectively, 1 month post-injection (Supplementary Fig. 6jk). Consequently, we saw contractions *in vivo* 1 and 2 months post-injection (Supplementary Fig. 7g-i; Fig. 2lm) with the instability index being lower and the contraction index decreasing over time, indicative of an increase in the number and/or a decrease in the size of the short alleles (Supplementary Fig. 8e-g). We conclude that our gene editing regimen induces contractions in adult R6/1 mice after symptom onset.

### Optimisation of the AAV constructs for robust expression in vivo

We next optimised the Cas9D10A construct further by using a cell-type specific codon optimization as described (patent WO2023105212A1). We found that the best construct for expression in the striatum amongst the ones tested was a skeletal muscle codon optimised Cas9D10A (v5 - Supplementary Table 3) used together with an optimised sgRNA scaffold containing an extra loop and a point mutation that improves editing efficiency^52^ injected at a ratio of 1:2 (Supplementary Fig. 10&11). Under these conditions, injecting 2-month-old animals yielded an average of 330 ± 92 and 1020 ± 173 AAV genomes per cell for the Cas9D10A v5 and the sgCTG-Mut+5 AAVs (Supplementary Fig. 12), respectively, 3 months post-injection. However, presumably because of Cas9D10A silencing, immunofluorescence data suggest that 19% of neurons expressing GFP also expressed Cas9D10A (Supplementary Fig. 13). We used these conditions to assess the impact of contractions *in vivo*.

### CAG repeat contractions translate to shorter polyglutamine stretches

We assessed whether contractions in the CAG/CTG repeat of the m*HTT* gene could be observed in the protein. To this end, we used Homogeneous Time Resolved Fluorescence (HTRF, Figure 3a)^53^. This assay uses striatal extracts together with antibody combinations to assess how close two epitopes are from each other within the mHTT protein. To establish the method in R6/1 mice, we first determined levels of soluble and aggregated mHTT. As determined for other HD mouse models^53^, we found that soluble mHTT decreases over the life span of the R6/1 mice and aggregated mHTT increases with age (Supplementary Fig. 14a-c). We then compared the levels of the various forms of mHTT in the mice injected with either Cas9D10A v5 alone or together with the sgCTG-Mut+5. We found no change in the amounts of aggregating (4C9 and MW8 antibodies) and soluble (2B7 and MW8 antibodies) mHTT (Fig 3bc). However, when we used the 2B7 antibody together with MW1, which targets the polyglutamine tract, we observed a reduction in signal (Fig. 3d, Supplementary Table 4), suggesting that fewer polyglutamines were found after the treatment with Cas9D10A *in vivo*.

**Figure 3.**
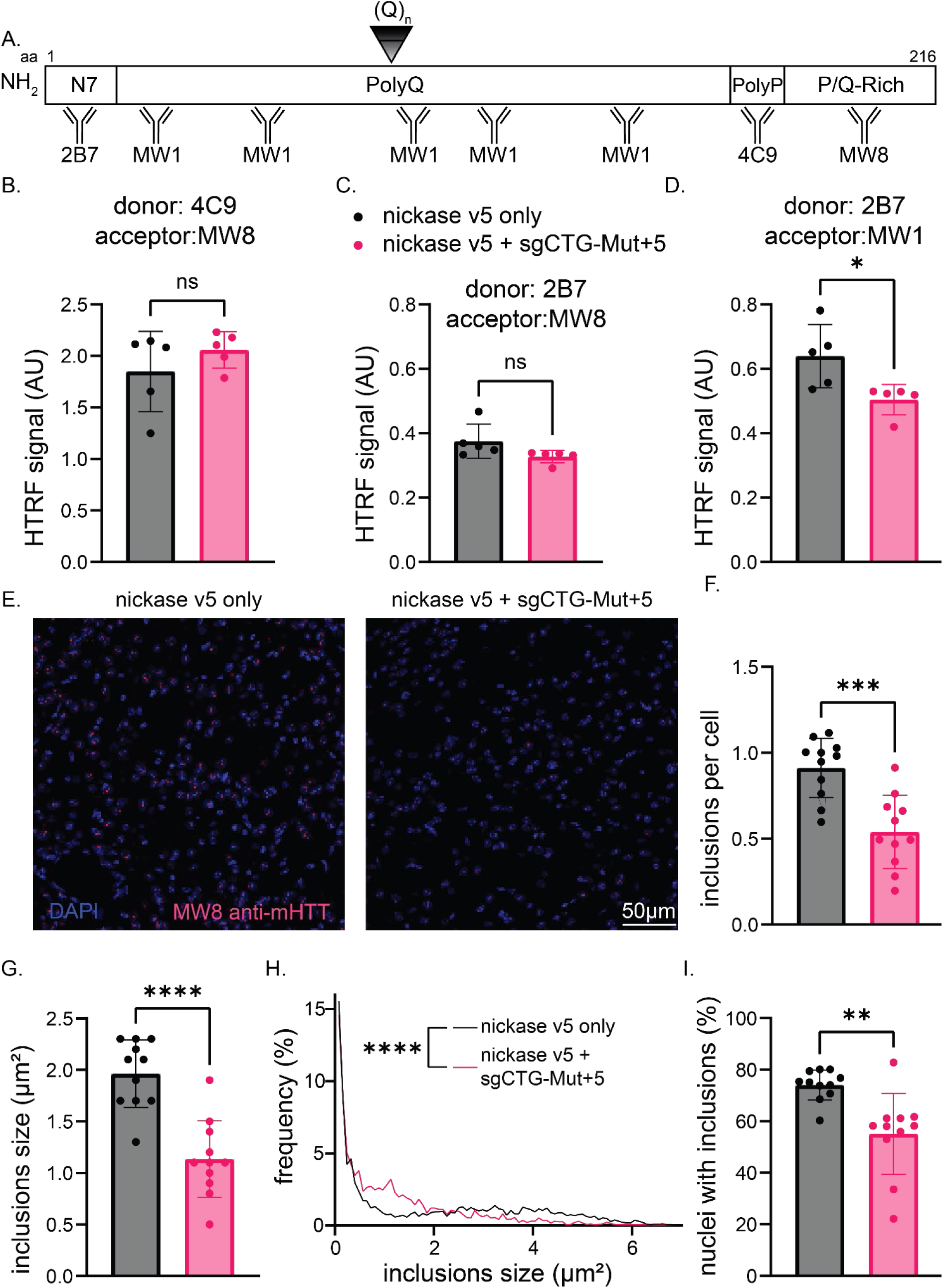
CRISPR-Cas9D10A injection reduces polyQ length and aggregation of mHTT *in vivo*. A) Schematic scaled diagram of a mHTT with 150 glutamines showing the relative location of HTT antibodies. Four antibodies detect mHTT exon 1: 2B7 maps to within the first 17 amino acids, MW1 detects expanded polyQ tracts, 4C9 detects the proline-rich region that lies between the two proline repeats in human HTT and MW8 acts as a neo-epitope antibody to the C-terminus of HTT. B) 4C9-Tb with MW8-d2 were used to track changes in aggregated mHTT, C) 2B7-Tb with MW8-d2 were used to track soluble mHTT and D) 2B7-Tb with MW1d2 pair for polyglutamine size assay by HTRF in striatal lysates from R6/1 mice at 5 months of age (3 months post-injection). (*P<0.05 using a t-test). Error bars represent ± standard deviation. Number of animals: 5 per group). E) representative z-projection images of striatal areas from 5-months-old R6/1 animal brains sacrificed 3 months post-injection of Cas9D10A only or Cas9D10A and sgCTG-Mut+5 and stained with DAPI (blue) and mutant polyglutamine inclusions (red). F) Analysis of total number of inclusions normalised to the total number of nuclei per image in striatal regions injected with Cas9D10A and the sgCTG-Mut+5 in comparison with their littermates that received Cas9D10A only. G) Two-dimensional analysis of the average aggregate size in treated animals. H) Distribution frequency of all analysed inclusions. Kolmogorov-Smirnov test to compare inclusion size distributions was used. P value <0.0001 (****)(n = 4009 and 2303 inclusions in Cas9D10A v5-injected only and Cas9D10A v5 + sgCTG-Mut+5 injected animals, respectively). I) Quantification of the number of mHTT positive nuclei in mice injected with Cas9D10A only or together with the sgCTG-Mut+5. Data are mean ± SD. fgi Student t-test were used (**P<0.01; ***P<0.001; ****P<0.0001 ). (n= 3-4 images per mouse, between 1000 and 1900 cells were assessed per mouse, with 3 mice per condition).

### Cas9 nickase and sgRNA delivery in vivo reduces the levels of mHTT inclusions

A molecular hallmark of HD in R6/1 mice is the presence of mHTT aggregates in the striatum^54^. To determine whether the injection of both the Cas9D10A and sgCTG-Mut+5 AAVs could improve this phenotype, we performed immunofluorescence using the specific MW8 antibody^55–57^. We found that the number of inclusions per cell as well as the size of the remaining inclusions decreased by over 40% in the striatum of mice injected with both AAVs compared to the ones injected with only Cas9D10A (Fig. 3e-h). Importantly, fewer cells in the striatum had inclusions (Fig. 3i). These results were specific for the injected region since a similar analysis in cortical areas of the same mice did not show any differences between groups (Supplementary Fig. 15d-f). We conclude that the injection of both the Cas9D10A and the sgCTG-Mut+5 AAVs decreases the burden of mHTT inclusions in the striatum. This is consistent with Cas9D10A slowing the transition from diffuse aggregates to inclusions suggested by the HTRF analysis.

### Nickase treatment partially mitigated gene expression alteration in the striatal medium spiny neurons

Dysregulation of transcription is a hallmark of HD pathology^58,59^. Thus, we aimed to determine whether the nickase could improve transcriptome dysregulation. To this end, we performed single-nuclei RNA sequencing of striatal samples from both wild type and R6/1 littermates 3 months after injecting them with either the Cas9D10A v5 alone or together with sgCTG-Mut+5 AAV. We first identified the medium spiny neuron population (Supplementary Fig. 16a-c, Fig. 4a) and determined the effect of the treatment by comparing wild type mice injected with only the Cas9D10A v5 AAV to those injected with both AAVs. We found that the treatment dysregulated only 100 genes in wild type mice (adjusted p-value = 0.01, fold change = 1.5; Supplementary Table 5, Supplementary Fig. 16d; Fig. 4b). Similar results were found using a pseudobulk approach (Supplementary Table 6, Supplementary Fig. 16e).

**Figure 4.**
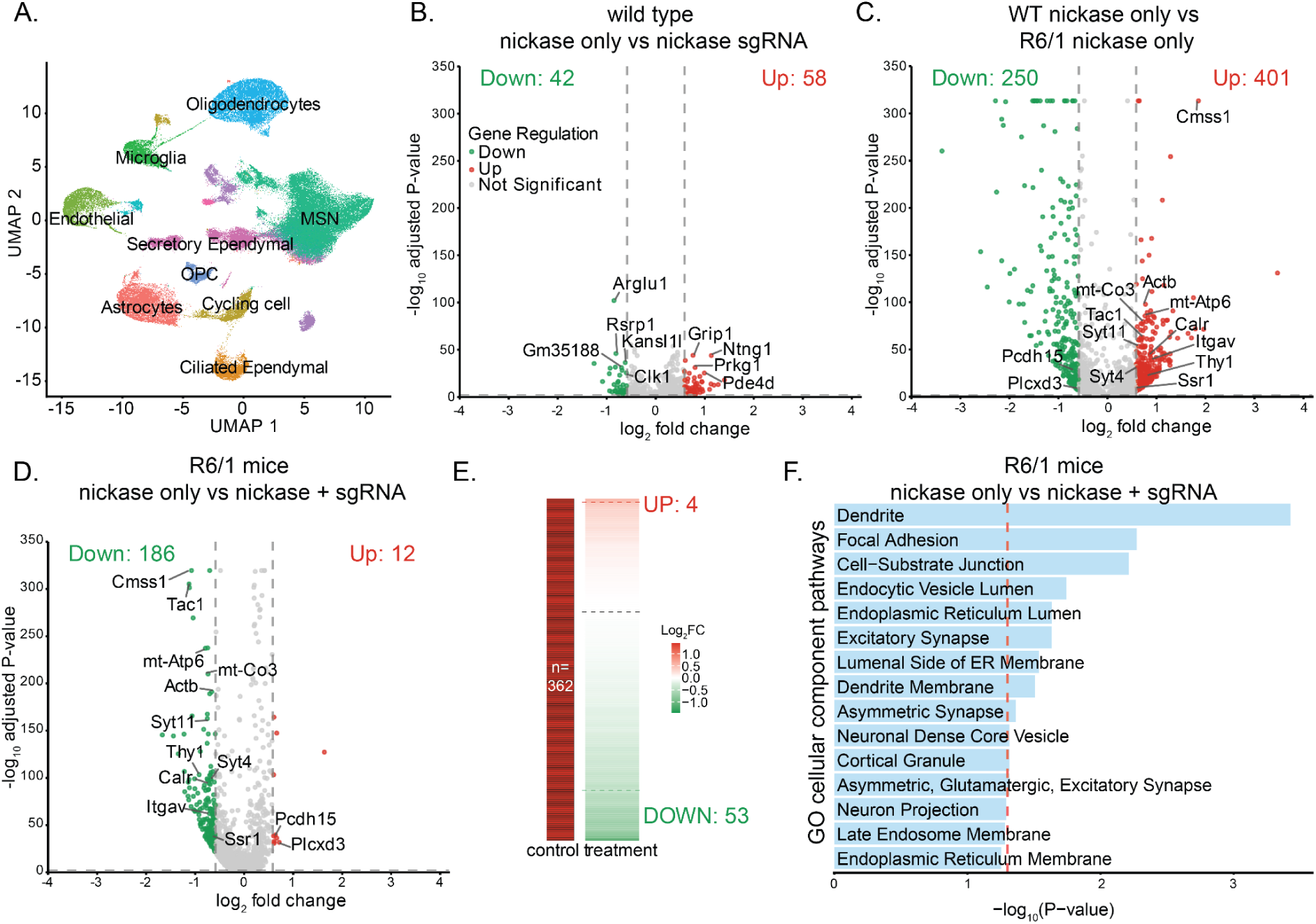
Cas9D10A co-injection with a sgCTG-Mut+5 against the repeat tract mitigates genes upregulated in HD medium spiny neurons 3-months post-injection. A) U-MAP plot of all nuclei across striatal regions from disease and control mice showing all major cell types in the brain (MSNs: medium spiny neurons). B) Volcano plot of differentially expressed genes (DEGs) in medium spiny neurons between wild type (WT) animals injected with Cas9D10A v5 only versus Cas9D10A v5 + sgCTG-Mut+5. C) Volcano plot for DEGs in medium spiny neurons between WT versus R6/1 animals injected with Cas9D10A v5 only. D) Volcano plot comparing DEGs in CRISPR-treated R6/1 medium spiny neurons relative to Cas9D10A v5-injected R6/1 medium spiny neurons. Number of animals: WT Cas9D10A v5 only n=3, WT Cas9D10A v5 + sgCTG-Mut+5 n=3, R6/1 Cas9D10A v5 only n=3, R6/1 Cas9D10A v5 + sgCTG-Mut+5 n=3. E) Heatmap of the changes in expression due to the co-injection of the Cas9 Cas9D10A v5 and sgCTG-Mut+5 specifically in the genes that are upregulated in the R6/1 medium spiny neurons compared to WT medium spiny neurons shown in C. The threshold for significance for differentially expressed genes was an adjusted P < 0.01 and an absolute log2 fold change > 0.585. F) Gene ontology (GO) cellular component pathways terms for DEGs which are altered by CRISPR injection in R6/1 medium spiny neurons comparing the treated mice with and without both Cas9D10A and sgCTG-Mut+5.

The effect of expressing mHTT led to more pronounced changes, R6/1 injected with Cas9D10A alone showed 651 dysregulated genes in medium spiny neurons compared to wild type littermates with the same treatment (Fig 4c, Supplementary Table 5). The pathways affected included dopaminergic synapse, calcium signaling, ribosomes, and multiple pathways of neurodegeneration, as expected from other HD models^58,59^. Similar results were obtained when we used a pseudo-bulk approach in medium spiny neurons with ion channels and neurotransmitter receptors being most affected (Supplementary Fig. 16f). These changes were comparable to those seen in two other HD mouse models^58,60^ (Supplementary Fig. 17).

Next, we compared transcriptome alterations caused by the expression of both the Cas9D10A and sgRNA-Mut+5 in medium spiny neurons in R6/1 mice (Fig. 4d). We found that the treatment led to 198 dysregulated genes, with 94% (186) of these being down-regulated. We assessed whether these changes rescued dysregulated genes caused by the presence of the HD transgene. To do so, we looked at the subset of genes that were significantly upregulated due to the R6/1 genotype (401 genes) and that were also found expressed in R6/1 medium spiny neurons from mice injected with both Cas9D10A and sgRNA-Mut+5 AAVs (362 genes). We found that in this subset, 15% (53 genes) showed a significantly increased expression (P<0.01, ❘log_2_FC<0.585❘), in the animals treated with both AAVs. (Fig. 4e). GO cellular component analysis showed that neuron to neuron synapse, dense core granule and synaptic vesicles genes were specifically rescued (Fig. 4f). These changes in gene expression were not due to a change in the number of medium spiny neurons or glia present in the dual injected R6/1 animals (Supplementary Fig. 16b) or to off-target binding of the Cas9 nickase as the misregulated genes were not enriched for genes containing potential off-target sites (Supplementary Table 5). We conclude that our gene editing treatment can mitigate some of the transcriptome alterations in the striatum of R6/1 mice.

### Intrastriatal Cas9 nickase and sgRNA injections improve motor performance

We investigated whether the Cas9D10A treatment could improve motor performance. Thus, we injected the striata bilaterally of 2-month-old wild type and R6/1 animals with the Cas9D10A v5 alone or together with sgCTG-Mut+5 (Fig. 5a) and tested them one month after injection. We found significant behavioural impairments in R6/1 mice expressing Cas9D10A only, versus wild type mice with the same treatment, including the balance beam and accelerating as well as fixed rotarod (Supplementary Fig. 18a-d, Supplementary Tables 7 to 10), as expected from previous work^61,62^. The wild type mice injected with both AAVs were not significantly different from those injected with only one AAV in these tests. When we performed an open field test (Fig. 5b), we found that wild type mice injected with both AAVs displayed less rearing than those injected with only the Cas9D10A AAV, but they remained unaffected in all other measures (Supplementary Fig. 19a-g). These results suggest that the treatment with both AAVs has little effect in wild type mice.

**Figure 5.**
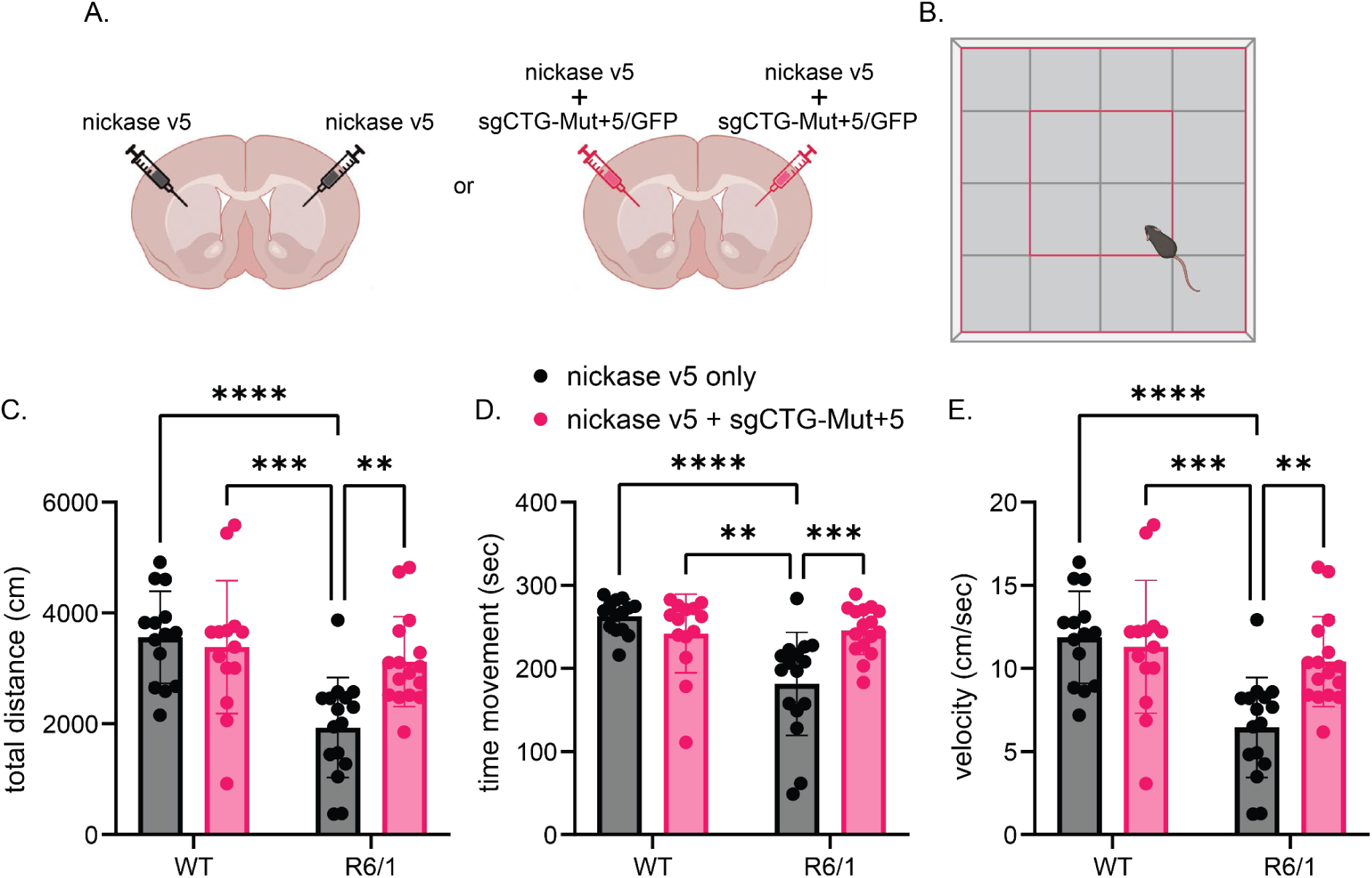
Injection of the Cas9D10A targeting the CAG repeats improves motor phenotypes in R6/1 HD mice in open field test. A) Schematic of the experimental approach with wild type (WT) and R6/1 animal bilaterally injected with the Cas9D10A v5 only or the Cas9D10A v5 and the sgCTG-Mut+5 AAVs. B) Open field schematic showing criteria adopted for defining arena areas; total area 80×80; external area of the arena 40×40 total; internal arena of the arena 40x40 in the centre of the arena (red square). C) Total distance travelled (WT Cas9D10A v5 only vs R6/1 Cas9D10A v5 only: P≤0.0001; R6/1 Cas9D10A v5 only vs R6/1 Cas9D10A v5 together with sgCTG-Mut+5: P≤0.004). D) Total time in movement (WT Cas9D10A v5 only vs R6/1 Cas9D10A v5 only: P≤0.0001; R6/1 Cas9D10A v5 only vs R6/1 Cas9D10A v5 together with sgCTG-Mut+5: P≤0.0005), and E) velocity (WT Cas9D10A v5 only vs R6/1 Cas9D10A v5 only: P≤0.0001; R6/1 Cas9D10A v5 only vs R6/1 Cas9D10A v5 together with sgCTG-Mut+5: P≤0.004) in WT and R6/1 mice injected with the Cas9D10A only or the Cas9D10A with sgCTG-Mut+5. Data presented is the average of two trials per mouse with the standard deviation between groups (for individual trials, see Supplementary Fig. 20). N of animals =14-16 per group. (**P<0.01; ***P<0.001; ****P<0.0001). Statistical analyses are in Supplementary Table 7.

When comparing the R6/1 mice expressing Cas9D10A alone in an open field test, we found significant behavioural impairments compared to wild type littermates injected with the same AAV. The R6/1 mice showed a 46% reduction in the distance travelled and velocity as well as a 31% reduction in the total time in movement (P≤0.004, Fig. 5c-e). When comparing R6/1 mice expressing both Cas9D10A and sgRNA to those that received only the Cas9D10A AAV, we found an improvement of 61% in distance travelled and velocity as well as a 35% increase in the total time in movement in R6/1 mice injected with both AAVs compared with those injected with only one AAV (P≤0.004, Fig. 5c-e). The R6/1 mice injected with both AAVs were not significantly different from the WT animals. Amelioration of the locomotor phenotype in dual-injected R6/1 mice was not due to changes in motor coordination or strength because parameters tested on the rotarod and balance beam were not affected by the injection of the CRISPR system in R6/1 mice (Supplementary Fig. 18; Supplementary Fig. 19fg). We conclude that the striatal injection of the Cas9D10A v5 together with sgCTG-Mut+5 improved gross locomotor phenotypes as measured by an open field test in R6/1 mice.

## Discussion

Here we provide evidence that Cas9D10A can induce contractions when targeted to expanded CAG/CTG repeats at different disease loci, cell types, and *in vivo*. Cas9D10A did not increase off-target mutation frequencies above background and contractions only occurred on the expanded allele. To our knowledge, the data presented here is the first example of a gene editing approach using a single Cas9 and sgRNA pair capable of correcting the mutation that causes clinically distinct disorders in a safe and efficient manner. *In vivo*, nickase-mediated contractions translated to fewer glutamines in the human mHTT fragment and were accompanied by the improvement of molecular and locomotor phenotypes of HD mice.

Importantly, contraction efficiency improved over time *in vivo*, suggesting that contracted alleles will accumulate for as long as expression of both Cas9D10A and sgCTG is maintained. Thus, the duration of the expression is expected to be a rate limiting factor for the editing efficiency and it is unknown how long expression would be needed for clinically relevant improvements in patients. On the other hand, the propensity of Cas9D10A to be silenced *in vivo* means that self-inactivating approaches, which involve the generation of double-stranded breaks^16^, may not be needed, potentially improving safety.

Although we speculate that contractions drive much of the improvements that we observed, other mechanisms are likely to be also at play. For instance, it may be that some of the changes in gene expression induced by the injection of the AAVs may be beneficial to HD pathology. Another possibility is that the reduction in somatic expansion that accompanies contractions plays a role. This is attractive because several drivers of somatic expansion impact the age at disease onset in HD^11,12^. However, reducing expansions when the repeat tract is already large, as in this mouse model, may have only limited effect on disease onset and progression^63,64^. In these cell types, improvements may require contractions beyond simply prevention of somatic expansion. We also considered that the presence of Cas9D10A bound to DNA would change expression of CAG/CTG-containing genes, but found no evidence to support this hypothesis from the snRNA-seq data. Intriguingly, the size of the mHTT inclusions were reduced uniformly across striatal sections and not only in cells that were infected by both AAVs, suggesting a potential non-cell autonomous effect of contractions on the surrounding cells. Such an effect may potentiate the improvements induced by the treatment. This is in line with previous observations that *HTT* silencing in neurons improved astrocyte phenotypes^65^. Moreover, Oura *et al*^66^ used a chimeric mouse approach to show that as few as 20% of edited cells, all of which had the same repeat excision, led to an improvement in HTT inclusion pathology, weight, and clasping phenotypes compared to R6/2 HD mice without any edited cells. Regardless of the exact mechanism, our data suggest that expressing Cas9D10A and a sgRNA against the repeat tract itself leads to significant improvements in the molecular and the locomotor phenotypes.

One limitation of our work is the use of two different AAVs to deliver Cas9D10A and sgCTG. This is common to other gene editing approaches as well^23,24^. One way around this is to use smaller Cas9 orthologs, for example the Cas9 from *S. aureus*^67^, as has been done *in vivo* for the *DMPK* locus^19,20^. Unexpectedly, this did not lead to higher levels of editing compared to a two-vector system^19,20^. Therefore, the size of the Cas9 enzyme and its delivery in the brain remain important challenges.

It is noteworthy that upon expression of the Cas9D10A we induced exclusively repeat contractions, rather than a collection of uncontrolled insertions and deletions^15–25,66,68–70^. Our approach is as efficient as excising the repeat tract, while remaining more precise and avoiding unwanted on-target and off-target mutations. This is of note because other approaches that involve base editing of the repeat tract targeted all CAG/CTG repeats, including non-expanded repeats^23,24^. The Cas9 nickase approach is therefore more precise and produces fewer unnecessary mutations.

A gene editing approach would have several advantages over more traditional approaches, including antisense oligonucleotides and microRNAs, which reduce the amount of *HTT* or *DMPK* mRNA^5,71,72^. Specifically, our gene editing approach would permanently improve the most proximal cause of the disease by reducing the size of the expanded CAG/CTG repeat. Recently, a small molecule that targets DNA secondary structures formed by expanded CAG/CTG repeats was shown to induce contractions *in vivo*^26,73^. However multiple stereotactic injections or continuous supply of the drug over 16 weeks were required for the effect to be significant. A gene editing approach would potentially require only a single dose. Crucially, from a patient benefit standpoint, it is imperative that we develop multiple approaches to maximise the chances of delivering successful drugs into the clinic.

## Methods

### iPSC cultures and differentiation conditions

For the HD iPSC-derived astrocytes, we obtained CS09iHD109-n1^38^ iPSCs from the Cedars-Sinai Medical Center’s David and Janet Polak Foundation Stem Cell Core Laboratory. We found the expanded *HTT* allele to be around 135 CAG repeats and the short one having 19 units. Cultures of iPSCs were tested regularly and found to be negative for the presence of mycoplasma using a service provided by Eurofins. IPSCs were grown in plates coated with Matrigel in E8 Flex Medium at 37°C, 5% CO_2_. For astrocyte differentiation iPSCs grown in E8 medium were supplemented with Rho-associated protein kinase inhibitor (RI) at approximately 60% confluency for 24 h. Neural progenitors (NPCs) were first derived by culture in SLI medium (advanced DMEM F-12 supplemented with 1x PenStrep, 1x Glutamax, 1% Neurobrew (w/o) vitamin A, 10 μM SB-431542, 200 nM LDN-193189, 1.5 μM IWR-1) for 8 days. On day 8, cells were treated with RI for 1 h and split 1:4 into NB medium (advanced DMEM F-12 1x PenStrep, 1x Glutamax, 2% Neurobrew (w/o) vitamin A) supplemented with RI for 24 h. From day 14 to 16, 20 μM PluriSIn was added to the medium to eliminate any undifferentiated iPSCs. On day 16, differentiated NPCs were frozen. Astrocyte progenitor cells (APCs) were differentiated from these NPCs as previously described^74^. Six million NPCs were thawed into NB medium containing 20 μM PluriSIn. After 2 days, the medium was changed to NF (advanced DMEM F-12 containing 1x PenStrep, 1x Glutamax, 2% Neurobrew with vitamin A, 20 ng ml^-1^ fibroblast growth factor 2 "Improved Sequence" (FGF-2 IS) and cells were allowed to reach confluency. They were then passaged three times in NEL medium (advanced DMEM F-12 with 1x PenStrep, 1x Glutamax, 2% Neurobrew with vitamin A, 20 ng ml^-1^ epidermal growth factor 20 ng ml^-1^ leukaemia inhibitory factor). CD44 positive cells were collected using the DB FACSAria Fusion and maintained into NEF medium (advanced DMEM F-12 supplemented with 1x PenStrep, 1x Glutamax, 2% Neurobrew with vitamin A, 20 ng ml^-1^ epidermal growth factor, 20 ng ml^-1^ FGF-2 IS). To mature astrocytes, APCs were seeded onto Matrigel coated 6 well plates at 200,000 cells per well with STEMdiff^TM^ Astrocyte Maturation medium (STEMCELL Technologies). After 2 weeks, the media was changed to advanced DMEM F-12 with 2% FBS for another 2 weeks. Once mature, astrocytes were plated in advanced DMEM F-12 medium without FBS for treatment with Cas9D10A.

CS09iHD109-n1 and DM1 iPSCs (GM24559, obtained from the Coriell Institute) were differentiated to cortical neurons^74^ by first bringing the cells to confluency and maintaining them in neuronal induction medium (0.5x advanced DMEM F-12 and 0.5x Neurobasal medium supplemented with 25 U ml^-1^ PenStrep, 2.5 µg ml^-1^ insulin solution human, 50 µM 2-Mercaptoethanol, 1x MEM non-essential amino acids solution, 500 µM sodium pyruvate solution, 1x N2 supplement, 1x B27 supplement, 1mM L-Glutamine, 10 μM SB-431542, 1 μM dorsomorphin dihydrochloride) for 12 days. The resulting neuroepithelium was disaggregated and plated in a new precoated 2x Matrigel well with neuronal maintenance media (NMM; 50x advanced DMEM F-12 and 50x Neurobasal medium supplemented with 25 U ml^-1^ PenStrep, 2.5µg ml^-1^ insulin, 50 µM 2-Mercaptoethanol, 1x MEM non-essential amino acids, 500 µM sodium pyruvate solution, 1x N2 supplement, 1x B27 supplement, and 1 mM L-Glutamine) also containing 20 ng ml^-1^ human FGF-2 IS. Once neuronal rosettes were visible, Neural Rosette Selection Reagent (Stem Cell Tech) was added to the culture for 1 h at 37°C for NPC selection. NPCs in suspension were collected and plated. NPC were maintained in NMM+FGF-2 IS. 50,000 live cells cm^-2^ and 30,000 live cells cm^-2^ of HD and DM1 NPC, respectively, were plated for neuronal differentiation in wells pre-coated with Laminin from Engelbreth-Holm-Swarm murine sarcoma basement membrane (Merck). Cells were plated in NMM and half the media was replaced twice a week. Neuronal cultures were kept for 3 weeks for maturation before being used for lentiviral transduction.

Mature neuronal and astrocytic cultures were transduced with lentiviral vectors at a multiplicity of infection of 5. The lentiviruses drive the expression of the Cas9D10A together with a blasticidin resistance gene, BSD, alone or together with the sgCTG cassette (Supplementary Table 1). Transduced cells were selected using blasticidin (5 µg ml^-1^) for 7 days. Cells were collected with scrapers 3 and 6 weeks after the infection to isolate DNA and measure repeat size.

### Lymphoblastoid cultures

Lymphoblastoid cell line from a Huntington’s disease patient (GM03620, obtained from the Coriell Institute) was cultured in RPMI with 2mM L-Glutamine + 15%FBS media. Cell cultures were transduced with lentiviral vectors expressing Cas9D10A (pLenti-EF1alpha-Cas9D10A nickase-Blast; Supplementary Table 1) at a multiplicity of infection of 10 along with 5 µg ml^-1^ polybrene. Transduced cells were selected using blasticidin (10 µg ml^-1^) for a minimum of 14 days. Then, cells were verified to be expressing Cas9D10A at multiple timepoints throughout to measure nickase protein expression via western blot (Supplementary Fig. 3a) and further transduced with a lentiviral vector expressing the sgCTG (pLV(gRNA)-CMV-eGFP:T2A:Hygro-U6(sgCTG); Supplementary Table 1) at a multiplicity of infection 20 + 5 µg ml^-1^ polybrene on day 0 of the experiment. Transduced cells were selected using hygromycin (50 µg ml^-1^) for 28 days. Cells were collected 42 and 56 days after the infection with the sgCTG to isolate DNA and measure repeat size using long-read sequencing.

### Immunostaining

Mature astrocytes were plated onto CELLSTAR 96-well tissue culture plates (Greiner Bio-One Ltd.), fixed with 4% paraformaldehyde (PFA), washed with PBS, permeabilized with 0.4% Triton-X in PBS, washed with PBS, and then blocked for 1 h at RT with 0.2% Triton-X in PBS + 1% normal goat serum (NGS, Merck). Antibodies used and their concentration is found in Supplementary Table 11. Cells were counterstained with DAPI (1 μg ml^-1^) before imaging with the Opera Phenix High Content Screening System (Perkin Elmer).

NPCs were plated on 13 mm coverslips for neuronal differentiation. Neurons were fixed in 4% PFA. Cells were washed in PBS, blocked, and permeabilized for 1 h in 1% BSA, 0.1% Triton X-100, and 4% NGS in PBS, and incubated with primary antibody in 1% BSA, 0.1% Triton X-100, and 1% NGS in PBS overnight at 4°C. After washing with PBS, cells were incubated with appropriate Alexa-conjugated secondary antibody in 1% BSA, 0.1% Triton X-100 in PBS for 1 h at room temperature. Cells were counterstained with DAPI Staining Reagent (1 μg ml^-1^; Chemometec) and after several washes in PBS, coverslips were mounted onto SuperFrost Plus slides (Thermo Fisher Scientific) with Fluoromount aqueous mounting medium (Merck) and imaged with the Leica SP8 confocal microscope. Images were captured using LAS X software and were analysed with Fiji software.

### Flow cytometry

Mature cultures were resuspended by Accutase (Fisher Scientific) and fixed with PFA 2%, Saponin 0,1%, in PBS. Fixation step was inactivated using 1.25 M glycine. Cells were washed/blocked with 500 µl of blocking solution (0.5% Saponin; 0.5% BSA; in PBS) and incubated with primary antibody in 300 µl 0.5% Saponin; 0.5% BSA; 1% NGS in PBS for 30 min at 4°C. After washing with a cleaning buffer ( 0.1% Saponin; 0.2% BSA; in PBS), cells were incubated with appropriate Alexa-conjugated secondary antibody in the cleaning buffer for 30 min at 4°C. Cells were counterstained with DAPI Staining Reagent (1 ng ml^-1^; Fisher Scientific). Finally, cells were resuspended in 0.5% BSA and EDTA 5 mM in PBS. Samples were run on the BD FACSAria Fusion and analysed using FlowJo v10 software.

### Small-pool PCR

Small-pool PCR was done as previously described^30,31^, but using primers listed in Supplementary Table 12. Genomic DNA was extracted with the NucleoSpin Tissue DNA extraction kit (Macherey-Nagel). PCRs were run using between 10-500 pg of DNA measured by Quant-iT Qubit dsDNA HS Assay Kit (ThermoFisher) to find the amount of amplifiable alleles for each sample. The products were run on 1.5% agarose gels in 1X TAE buffer and transferred onto a GeneScreen Hybridization Transfer Membrane. The hybridization was done at 48°C (for *HTT* blots) or 51°C (for *DMPK* blots) in ULTRAhyb Ultrasensitive Hybridization buffer (Fisher Scientific) with a ^32^P-labelled oligo containing 10 CAG repeats (oVIN-100, Supplementary Table 12). The membranes were exposed to a phosphor screen and scanned using Bio-Rad PMI Personal Molecular Imager. Images were captured using Quantity One 1-D Analysis Software (Basic V4.6.6) and analysed with Fiji version 1.53f51. Blinded allele counts were completed for each membrane based on allele frequency calculated by Poisson distribution as described^31^. Each band was sized using the ladder run on either side of the gel. For the *HTT* locus in astrocytes, we excluded repeats below 34 repeats so as to not conflate contractions with the non-expanded allele. For the *DMPK* locus, small-pool PCRs, sizing was further divided between fragment sizes of 2 kb and 3 kb by fitting an exponential decay of the distance travelled by the ladder bands. This allowed us to subdivide this region of over 580 repeats into three bins, where most of the contractions were observed. Differences between conditions were determined using a Mann-Whitney U Test calculated by GraphPad Prism software (v.10.0.0).

### Potential off-target identification

To assess potential off-target effects of our CRISPR-Cas9 system targeting CAG/CTG repeats in the human genome (GRCh38/hg38), we used the CasOFFinder web tool^75^. We used two different queries CTGCTGCTGCTGCTGCTG and CTGCTGCTGCTGCTGCTGC to capture potential binding. All analyses were conducted using the 5’-NRG-3’ PAM sequence, allowing up to 2 mismatches. This resulted in 1846 potential off-target genes genome-wide. Coverage analysis of those genes revealed 16 genes with very low coverage, which were filtered out, leading to a total of 1830 potential off-target loci (Supplementary Table 2). The same queries were used on the mouse genome (mm10). We found 3132 potential off-targets (Supplementary Table 5).

### Whole genome sequencing

Genomic DNA was extracted from cells using the Macherey-Nagel NucleoSpin Tissue DNA mini kit and made into dual-indexed single stranded DNA libraries using ‘Illumina DNA PCR-free Prep kit, Tagmentation’ (Illumina). Quantification of the libraries was done using the KAPA Illumina Universal Library Quant kit (Roche). The sequencing was performed on an Illumina NovaSeq 6000 using the v1.5 S4 reagent kit, 300 cycles using XP 4-lane splitter kit. Together the 33 samples generated 1600Gb of raw data.

Demultiplexing was done using bcl2fastq conversion software (Illumina) and the paired-end sequences were aligned to GRCh38/hg38 with the Burrows-Wheeler Aligner (BWA)^76^ version bwa/0.7.17 and the mem algorithm. Subsequently, the Sequence Alignment/Map (SAM)^77^ files produced were sorted via the Picard (picard/2.27.5 - https://broadinstitute.github.io/picard/). These files were then converted to binary alignment/map (BAM) format and indexed using SAMtools (samtools/1.21)^77^.

To assess sequence coverage, genome-wide and potential off-target candidate regions, Mosdepth^78^ (0.3.11) was applied to all BAM files. The resulting coverage bed files were combined to create an average coverage using an in-house python script (available on https://github.com/DionLab/Murillo-et-al). Read groups were attributed to the BAM files with the Picard tool, which also facilitated the subsequent analysis for single nucleotide polymorphisms using bcftools^79^ version 1.21 with htslib version 1.22.

To capture rare mosaic variants, we used an approach similar to one used by Dong et al^80^. Variant calling was performed on the whole genome as well as candidate off-target regions specified in the BED file using Bcftools mpileup^77^. Variants with an allele frequency of at least 0.05 and DP2+DP3 ≥ 2. The output was compressed to BCF format following the standard bcftools workflow for each sample. Variants were further filtered for quality using the bcftools filter, excluding those with the following criteria: “QUAL < 30 or DP < 10 or MQ<40 or VAF ≥ 0.2, calculated as (DP4[2] +DP4[3]) / (DP4[0]+DP4[1]+DP4[2]+DP4[3])”. Filtered results from all VCF files for 33 samples with 4 lanes were subsequently compiled into a single CSV (Supplementary Table 2) format using a custom Python script for further analysis. Only variants that were found in more than one lane were considered and we removed the ones found at day 0 as they were already in the population before the treatments. We ran this analysis on both the 1830 potential off-targets and the whole genome separately.

### Long read sequencing of repeats

Expanded repeats were sequenced as before using SMRT HiFi amplicon sequencing^44^ (see Supplementary Table 12 for primers). Mouse samples were mechanically disaggregated in Nucleospin Tissue DNA extraction T1 buffer using a mini hand-held homogenizer (ThermoFisher). Genomic DNA was extracted with the NucleoSpin Tissue DNA extraction kit (Macherey-Nagel). Three PCRs (3 x 50 µl volume) per sample were combined and cleaned with AMPure PB beads according to PacBio’s barcoded overhang adapter protocol (Pacific Biosciences #101-791-700). Barcoded sample pool purity was analysed on the 5200 Fragment Analyser (Agilent) and final library concentration determined using Invitrogen Qubit 1X HS dsDNA kit. Sample pools were sequenced using the Pacific Biosciences Sequel IIe.

Precise CAG/CTG repeat sizes were determined using Repeat Detector^44^ (version 1.0.15eb445). Unaligned reads were assessed using the restrictive profile with a repeat size range of [0–250]. For each analysis, the -–with-revcomp option was enabled and data was output to a histogram (-o histogram option). Obtained histograms were plotted using GraphPad Prism software (v.10.0.0). As the sequences are unaligned when we determine repeat size, the CCSs containing between 0 and 5 CAG/CTGs are removed from analysis as they include a large proportion of truncated PCR products and other sequences that do not align with the *HTT* locus.

### Alignment of long reads

The repeat regions were aligned with the NGMLR aligner using the *HTT* region from GRCh38 containing 19 CAG repeats as reference sequence. The resulting SAM files were converted into BAM and .bai index files using Picard and Samtools, respectively. The bam files were checked for percentage of mapped reads using Samtools flagstat command. The indexed BAM files were then visualised using the Integrative Genomics Viewer (IGV 2.16.2)^46^ to assess the alignment quality and to confirm the presence of the expected CAG repeat count. Sniffles2^45^ was run on the aligned BAM files for identifying rearrangements.

### Area under the curve, repeat size indices, and comparisons

To calculate the difference in the area under the curve (Supplementary Fig. 9), we used the histograms generated by Repeat Detector. The total read counts were normalised for each sample and adjusted such that the mode is set to 0. Then we subtracted the frequency of reads of the treated samples from those in the control conditions for each repeat size. The positive values were then added together to produce the difference in the area under the curve. It represents the frequency of alleles that has changed compared to the control samples. We also generated delta plots to compare the differences in repeat size between two treatments. To do so, allele frequencies were normalized to the number of total reads in the samples and grouped into 5-repeat bins. Then the relative frequencies of the experimental samples were deducted from that of the control sample. Statistical distribution analysis using a matched-pair Wilcoxon or unpaired Kolmogorov-Smirnov tests was performed without binning. The instability index was calculated as previously described for images^51^, but using normalised read frequencies and a 5% threshold. The expansion and contraction indices were calculated such that expansion index + contraction index = instability index.

### Mouse housing

C57BL/6J R6/1 mice^81^ (B6.Cg-Tg(HDexon1)61Gpb/J) were maintained as hemizygotes with *ad libitum* access to food and water and maintained in a temperature and humidity-controlled environment on a 12 h dark/light cycle. Health and wellbeing of the animals were monitored daily and their weight were checked every week in a T1 biosafety level 2 animal facility. All experimental procedures done in Cardiff followed protocols in accordance with the United Kingdom Animals (Scientific Procedures) Act of 1986. All experimental procedures performed on R6/1 mice were approved by Cardiff University Animal Welfare and Ethical Review Board and carried out under Home Office Licenses P49E8C976 and PP7595333. We did not go beyond 2 months in these experiments because the R6/1 mice become markedly unwell beyond this age and to continue would have risked unnecessary suffering. Stereotactic injections done in wild type mice presented in Supplementary Fig. 6a-d were done at the Center de Recherche du CHU de Québec (Québec, QC, Canada). All procedures on these animals were completed in accordance with the guidelines of the Canadian Council on Animal Care and were approved by the Comité de Protection des Animaux du CRCHUQ-UL under protocol number 2019-49, CHU-17-106.

### Stereotactic injections

The R6/1 mice used here were of both sexes and had between 138 and 151 CAG/CTG repeats. Mice were anaesthetised with isoflurane and placed in a stereotactic frame. In P2 animals, a syringe was used to penetrate the skin and skull. In adult (3.5 months old) mice, the scalp was shaved, a longitudinal incision was made to expose the skull surface, and 2 burr holes were drilled above the infusion sites. A 4 μl viral suspension (4.8×10^10^ vg of Cas9D10A v4 and sgCTG/GFP AAVs in 1:1 ratio) was stereotactically injected into the striatum of adult mice according to the Paxinos and Franklin mouse brain atlas (AP +0.8; ML +-1.8; DV -2.2 mm from bregma). 2-months old animals were injected in the same adult coordinates with a 4 μl viral suspension (7.98×10^10^ vg of Cas9D10A v5 and sgCTG-Mut+5/GFP AAVs in 1:2 ratio). P2 mice were stereotactically injected with 1 µl (1.44×10^10^vg of Cas9D10A v4 and sgCTG/GFP AAVs in 1:1 ratio) virus suspension into the striatum (AP +2.3; ML +-1.4; DL 1.8 mm from lambda). Hamilton syringes 701 N 10 µl and 5 µl FIX NDL (26S/51/3) were used for the infusion in adult and post-natal mice respectively. The infusion rate was 200 nL min^-1^, and the needle remained in place for 5 min after infusion for vector absorption before removal. In adult mice, the injection site was closed with stitches, and mice recovered in incubators. For the wild type animals injected with the different constructs found in Supplementary Fig. 6a-d, each construct was injected together with the sgCTG AAV9 into the cortex and the striatum (AP: +/- 1.6, MD: +/- 1.4, DV -0.75) using a total of 6×10^10^ vg, at a ratio of Cas9D10A to sgCTG of 1:1, divided equally between the two hemispheres. Animals were between 8 and 9 months of age.

### Behavioural tests

Male and female mice were injected at 2 months of age and tested 1 month post-injection. Mice were handled for 1 minute per day for 5 days each before the first behavioural test. For motor coordination assessment, mice were placed on a rotarod with a fixed speed of 12 rpm and 24 rpm (two total trials, 3 hours apart) and the time of the first fall and the total number of falls were recorded until the sum of latencies to fall reached a total of 60 s per trial. For accelerated rotarod, animals were placed on the rod with a constant increase from 5 to 40 rpm over the 5-minute trial. A balance beam test was used to evaluate fine motor coordination and balance. Mice were placed at one end of the beam, and the time to reach an escape box containing nesting material and located in the opposite end was recorded. The beam dimensions were: L80 cm, W0.5–1.5 cm, H34–54 cm with an incline of 17°. The house dimensions were: L11 cm, W11 cm, H10 cm. Mice were allowed to rest for 3 hours before the next trial with a total of two trials. Muscular strength was measured by placing the mice on top of a wire rectangle of approximately 20 cm × 21 cm, surrounded by tape to prevent mice walking off the edge, and after a light shaking so mice gripped the wires, the lid was turned upside down. Latency to the first fall was recorded within the total 60 s of the test. Open field test was conducted by positioning the animal at the centre of the white open field arena (80 cm x 80 cm x 30 cm). The locomotive behaviour was monitored for a duration of 5 minutes with the EthoVision tracking software. All experiments were conducted during the light phase and were performed and analysed in a blinded manner.

### Perfusion and immunostaining of mouse tissues

R6/1 animals were deeply anaesthetised with Dolethal (Vetoquino) and transcardially perfused with 4% PFA in PBS. Brains were removed, 2 h post-fixed in 4% PFA, and stored in 30% sucrose in PBS at 4°C. Wild type animals presented in Supplementary Figure 6a-d were anaesthetised the same way, but the perfusion was done with PBS. 30 µm coronal sections were cut with a freezing sliding microtome and stored at −20°C in cryoprotectant solution (30% ethylene glycol, 30% glycerol, 20 mM PBS) until processing. Free-floating brain sections were washed in PBS and an antigen retrieval step was performed. For IBA1, GFAP and HA antibodies immunostaining, tissue sections were incubated for 30 minutes at 75°C in sodium citrate buffer pH 6. For GFPs, NeuN and DARPP-32 immunostaining, tissue sections were boiled in citrate buffer pH 6 for 2 minutes. Brain sections were then washed in PBS and blocked and permeabilized for 1 h in 1% BSA, 0.2% Triton X-100, and 4% NGS in PBS for IBA1 and GFAP antibodies; 1 h in 1% BSA, 0.3% Triton X-100, and 4% NGS in PBS for NeuN and DARPP-32 antibodies; 1 h in 1% BSA, 0.5% Triton X-100, and 4% NGS in PBS for HA antibody. Then, floating sections were incubated with primary antibodies in 1% BSA, 0.1% Triton X-100, and 1% NGS in PBS overnight at 4°C. After washing with PBS, sections were incubated with appropriate Alexa-conjugated secondary antibody in 1% BSA, 0.1% Triton X-100 in PBS for 1 h at room temperature. Tissues were stained with DAPI and after several washes in PBS, sections were mounted onto SuperFrost Plus slides (Thermo Fisher Scientific), air-dried, and coverslipped with Fluoromount aqueous mounting medium (Merck). Leica SP8 confocal (LAS X software) and Evos FL Auto 2 (Invitrogen EVOS FL Auto 2.0 Imaging System) microscopes were used to capture images.

Quantification of the inclusion bodies was done using the total number of cells or nuclei, as measured by DAPI staining, taking 3 to 4 images from 3 different mice at several different antero-posterior and dorso-ventral coordinates in the striatum. The reported decreases were calculated as 100% minus the percentage of the Cas9D10A treatment alone. We reported the results the same way for the size of the inclusion bodies and their intensities. All image capture and analysis were done where we are blind to the treatment done. Total cells and inclusions quantification were performed with ImageJ (v2.3.051).

### Western blotting

To assess Cas9 levels in astrocytic and lymphoblastoid cultures, proteins were extracted using RIPA buffer supplemented with cOmplete^TM^ EDTA-free protease inhibitors. Mouse brain tissues were mechanically disaggregated in RIPA buffer supplemented with cOmplete^TM^ EDTA-free protease inhibitors using a mini hand-held homogenizer (ThermoFisher). Samples were incubated for 30 min in a rotator at 4°C and soluble fractions were collected by centrifugation (30 min; 13000g at 4°C). Protein concentrations were determined using Pierce BCA protein assay (ThermoFisher). 10 to 30 μg protein was loaded on 4-12% Bis-Tris Plus precast gels (ThermoFisher) and run in 1X MES buffer (ThermoFisher) alongside Bio-Rad Kaleidoscope molecular weight marker. Proteins were transferred onto a nitrocellulose membrane using the Bolt system (ThermoFisher). The results were imaged on the Odyssey Imager (LI-COR Biosciences).

### HTRF assay

HTRF assay was performed as previously described^53,82^. In summary, a 5% (w/v) total protein homogenate was prepared in ice-cold bioassay buffer (PBS, 1% Triton-X-100) with complete protease inhibitor cocktail tablets (Roche), by homogenizing three times for 30 s in lysing matrix D tubes at 6.5 m s^-1^ (MP Biomedicals) in a Fast-Prep-24TM instrument (MP Biomedicals). Lysates were snap frozen and used for assays the following day. Tissue homogenates to a final volume of 10 μL were pipetted in triplicate into a 384-well proxiplate (Greiner Bio-One). Antibody concentrations used were 1 ng of donor per well and 40 ng of acceptor per well. For HTRF assays, antibodies were added per well in 5 μL HTRF detection buffer [50 mM NaH2PO4, 0.2 M KF, 0.1% bovine serum albumin, 0.05% Tween-20] with complete protease inhibitor cocktail tablets (Roche). Plates were incubated for 3 h on an orbital shaker (250 rpm) at room temperature, before reading on an EnVision (Revvity) plate reader using optimised HTRF detection parameters as described previously^82^.

### Single-nuclei RNA sequencing and analysis

#### Library preparation and sequencing

Nuclear isolation was performed in a Genomics Chromium platform from 12 striatal samples. Droplet-based snRNA sequencing libraries were prepared at the UK DRI Single Cell and Spatial Omics Facility using the Chromium Next GEM Single Cell 3’ Kit v3.1 (10x Genomics, Pleasanton CA) according to the manufacturer’s protocol. The library was sequenced at UCL genomic facility using NovaSeq 6000 S4 v1.5 (200 Cycles) with configuration 28-10-10-90.

#### Processing, quality control, and filtering

Raw FASTQ files were processed using Cell Ranger v8.0.0 ^83^(10x Genomics) on a high-performance computing cluster. A custom mouse reference genome was created by adding GFP, Cas9D10A, and human huntingtin exon1 (*huHTT*) transgene sequences to the standard mouse genome (GRCm39). The data were processed using Seurat v5.0 in R^84–87^. Quality control metrics were calculated including mitochondrial gene percentage using the PercentageFeatureSet() function with the pattern "^mt-". Cells were filtered to retain those with 200-5000 detected genes and less than 5% mitochondrial reads. Sex information was mapped to each sample and converted to numeric format for downstream regression.

#### Normalization and batch effect correction

Unique Molecular Identifier (UMI) counts from each cell were normalized using SCTransform with the mitochondrial gene percentage, total UMI count (nCount_RNA), and sex regressed out using vars.to.regress^88^. Principal component analysis was performed on the normalized data using RunPCA() with default parameters, and the first 20 principal components were selected for downstream analysis. Batch effect correction was performed using Harmony integration implemented through the RunHarmony() function^89^. Integration was performed on the SCTransform-normalized data using sample identity (orig.ident) as the batch variable with theta = 1. The integrated representation was saved as "pca_integrated" for subsequent clustering and visualization.

#### Cell clustering, visualization, and identification

Cell clustering was performed using the integrated principal components with the FindNeighbors() and FindClusters() functions in Seurat^84–87,90^. A resolution of 0.5 was used for Louvain clustering. UMAP (Uniform Manifold Approximation and Projection) dimensionality reduction was performed using RunUMAP() on both unintegrated and integrated PCA embeddings with default parameters.

Cell type annotation was performed using a curated list of marker genes for 13 cell types^58^. Direct and indirect medium spiny neurons were pooled for the subsequent analyses. Module scores were calculated for each cell type using AddModuleScore() with the respective marker gene sets. Automatic cell type assignment was performed by calculating average module scores for each cluster and assigning the cell type with the highest average score using Seurat v5.0 ^84–88,90^.

#### Single-nuclei differential expression analysis

Cell type-specific differential expression analysis was performed using FindMarkers() implemented in Seurat v5.0 ^84–88,90^. Prior to analysis, SCTransform models were prepared using PrepSCTFindMarkers() to ensure compatibility across sample subsets.

Differential expression analysis was conducted using Seurat’s FindMarkers function with the MAST framework. Analysis was restricted to genes expressed in at least 10% of cells in either comparison group, with logfc.threshold = 0.

#### Pseudo-bulk differential expression analysis

We performed pseudobulk differential expression (DE) analysis separately on each annotated cell type using DESeq2 (v1.48.2)^91^. For each annotated cell type, raw UMI counts from the Seurat object were aggregated by its *orig.ident* to generate sample-level pseudobulk count matrices, treating each replicate as a separate unit for statistical testing. Genes with less than 10 UMI counts in all samples or expressed in less than two replicates per group were excluded. Differential expression testing was performed using a design formula of ∼ sample_group. For each comparison, DESeq2 was utilized to estimate dispersion and fit negative binomial generalized linear models. This resulted in csv files with normalized counts and differential gene expression statistics for all genes (Supplementary Tables 5 and 6).

#### Filtering and pathway enrichment analysis

The threshold for significance for differentially expressed genes were adjusted p-value < 0.01 and absolute log_2_ fold change > 0.585. Pathway enrichment analysis was performed on these filtered gene sets using the enrichR package. The Gene Ontology Cellular Component (2023 version) was used and the enrichment results were ranked by p-value and combined score, with specific tracking of Huntington’s disease-related pathways across all comparisons. For KEGG pathway analysis, Tubb3 emerged as the most stable housekeeping gene calculated by the coefficient of variation and used as housekeeping gene for DGE normalization. We used a targeted gene panel from the KEGG Huntington’s Disease pathway (<u>hsa05016</u>, <u>MSigDB M13486</u>), which is available through the Molecular Signatures Database.

#### Comparison with previously published datasets

To compare gene expression at the single-nucleus level against published datasets, we first subdivided our MSN population into the direct (dMSN) and indirect (iMSN) medium spiny neurons. For each subtype, we extracted lists of significantly differentially expressed genes comparing R6/1 mice and wildtype littermates treated with the Cas9D10A AAV only. These were compared to differentially expressed genes reported by Lee et al^58^ and Lim et al^92^ for R6/2 mice as well as compared to zQ175 mice^58^. We determined the direction of gene expression changes by using the sign of the log₂ fold-change in each dataset and assessed whether they were concordant with our observations. A Fisher’s exact test was used to determine whether the number of common / divergent genes were different between datasets (Supplementary Fig. 17).

#### Off-target enrichment analysis

We extracted the set of genes overlapping Cas9D10A potential off-target loci in the mouse genome. We used a Fisher’s exact test (Supplementary Table 5) to determine whether off-target containing genes were more likely to be differentially regulated compared to the total number of dysregulated genes. We used the pseudobulk medium spiny neurons for this analysis (Supplementary Fig. 16e-g).

### Q-PCR

Quantitative real-time PCR (qPCR) was performed using the Applied Biosystems QuantStudio 7 Flex Real-Time PCR system and analysed with the QuantStudio Real-Time PCR Software (v1.7 Thermo Fisher). Briefly, qPCR was assayed in a total volume of 10 μl reaction mixture containing the ready-to-use FastStart Universal SYBR Green Master (ROX) (Thermo Fisher) and 0.5 µM of forward and reverse mix appropriate primers (Supplementary Table 12). All qPCR reactions were run in triplicates. Mean cycle threshold (Ct) values for each reaction were recorded and the relative DNA copies were calculated and normalised to Actin (2^(ActinCt-geneCt)).

### Statistics

Total number of alleles amplified by Small-pool PCR was calculated using Poisson distribution as described^30^. We determined P-values for small-pool PCR using a Mann-Whitney nonparametric U-test comparing control and Cas9D10A + sgCTG conditions. We determined statistical significance between the Cas9D10A only and Cas9D10A + sgCTG HD iPSC-derived neuronal populations shown in Supplementary Fig. 2cd using a Student’s t-test. A nonparametric paired Wilcoxon test was used to compare two hemispheres from the same animal and a nonparametric unpaired Kolmogorov-Smirnov test when comparing samples from different animals. Delta plots used aggregated data visualization purposes only, not for statistical testing. A Student’s t-test was also used HTRF and aggregate measures. Furthermore, differences in instability indices over time were calculated using a one-way ANOVA with a post-hoc Tukey’s multiple comparison test. We used two-way ANOVA to estimate differences on treatment and time, followed by a post-hoc Tukey’s multiple comparison test. The same test was used for behavioral analysis. For repeat size comparison, in allele frequency or delta plot analysis, after long-read sequencing two tests were used. For comparison of repeat instability between striatal samples within the same animal a paired nonparametric Wilcoxon signed rank test was used. For comparison of repeat sizes between different animals and in the lymphoblastoid experiments, an unpaired nonparametric Kolmogorov-Smirnov test was used. Statistical analyses were done using GraphPad Prism software version 10.0.0 and were always two-tailed tests using a significance cut off at P=0.05.

## Data availability statement

The CCSs and snRNA-seq sequences are available from SRA (https://www.ncbi.nlm.nih.gov/sra) PRJNA1077893.

## Code availability statement

Custom scripts are available at https://github.com/DionLab/Murillo-et-al.

## Acknowledgements

Mouse image (Fig. 5b) and brain slice (Fig. 2bj&5a) were created with <u>BioRender.com</u> (agreement numbers are MK28EFZJ1S and JM28EFZ3G9, respectively). We would like to thank Andrew Jefferson for training and for keeping the fluorescence microscopes in excellent working order and Joanne Morgan and the core team at the Centre for Neuropsychiatric Genetics and Genomics at Cardiff University for support with sequencing. Also, we would also like to thank Daria Gavriouchkina and Modesta Blunskyte from the UK DRI Single Cell and Spatial Omics Facility for their help in single nuclei RNA-seq library preparation. We thank the UCL genomic facility staff for the snRNA-seq sequencing. We thank Vanessa Wheeler and Geneviève Gourdon for critical reading of the manuscript. We also thank Sarah Langley for advice with the snRNA-seq data analysis and Sandeep Sundara Rajan for designing the Cas9D10A v5. This research was undertaken using the supercomputing facilities at Cardiff University operated by Advanced Research Computing at Cardiff (ARCCA) on behalf of the Cardiff Supercomputing Facility and the HPC Wales and Supercomputing Wales (SCW) projects. We acknowledge the support of the latter, which is part-funded by the European Regional Development Fund (ERDF) via the Welsh Government. We also thank the University of Lausanne for providing some plasmids used in this study.

## Author contributions

AMurillo: Conceptualization, Formal Analysis, Investigation, Methodology, Supervision, Visualization, Writing – original draft, Writing – review & editing

MA: Conceptualization, Formal Analysis, Investigation, Methodology, Visualization, Writing – review & editing

RRPD: Formal Analysis, Visualization, Writing – review & editing

ELR: Methodology, Writing – review & editing

MLarin: Conceptualization, Formal Analysis, Investigation, Methodology, Visualization, Writing – review & editing

LH: Methodology, Writing – review & editing

ANA: Methodology, Writing – review & editing

AST: Formal Analysis, Visualization, Writing – review & editing

AMM: Resources, Writing – review & editing

NS: Investigation, Resources, Writing – review & editing

AERH: Formal Analysis, Writing – review & editing

PA: Methodology, Writing – review & editing

KF: Methodology, Writing – review & editing

CL: Methodology, Writing – review & editing

GFO: Methodology, Writing – review & editing

AMangin: Methodology, Writing – review & editing

SB: Methodology, Writing – review & editing

EM: Resources, Supervision, Writing – review & editing

NDA: Resources, Funding acquisition, Supervision, Writing – review & editing

JP: Conceptualization, Supervision, Funding Acquisition, Writing – review & editing

GPB: Resources, Supervision, Writing – review & editing

BLD: Resources, Supervision, Writing – review & editing

FC: Conceptualization, Funding acquisition, Supervision, Writing – review & editing

MLelos: Conceptualization, Funding acquisition, Supervision, Writing – review & editing

VD: Conceptualization, Funding acquisition, Project administration, Supervision, Visualization, Writing – original draft, Writing – review & editing

## Supplementary Material

**Supplementary Figure 1:**
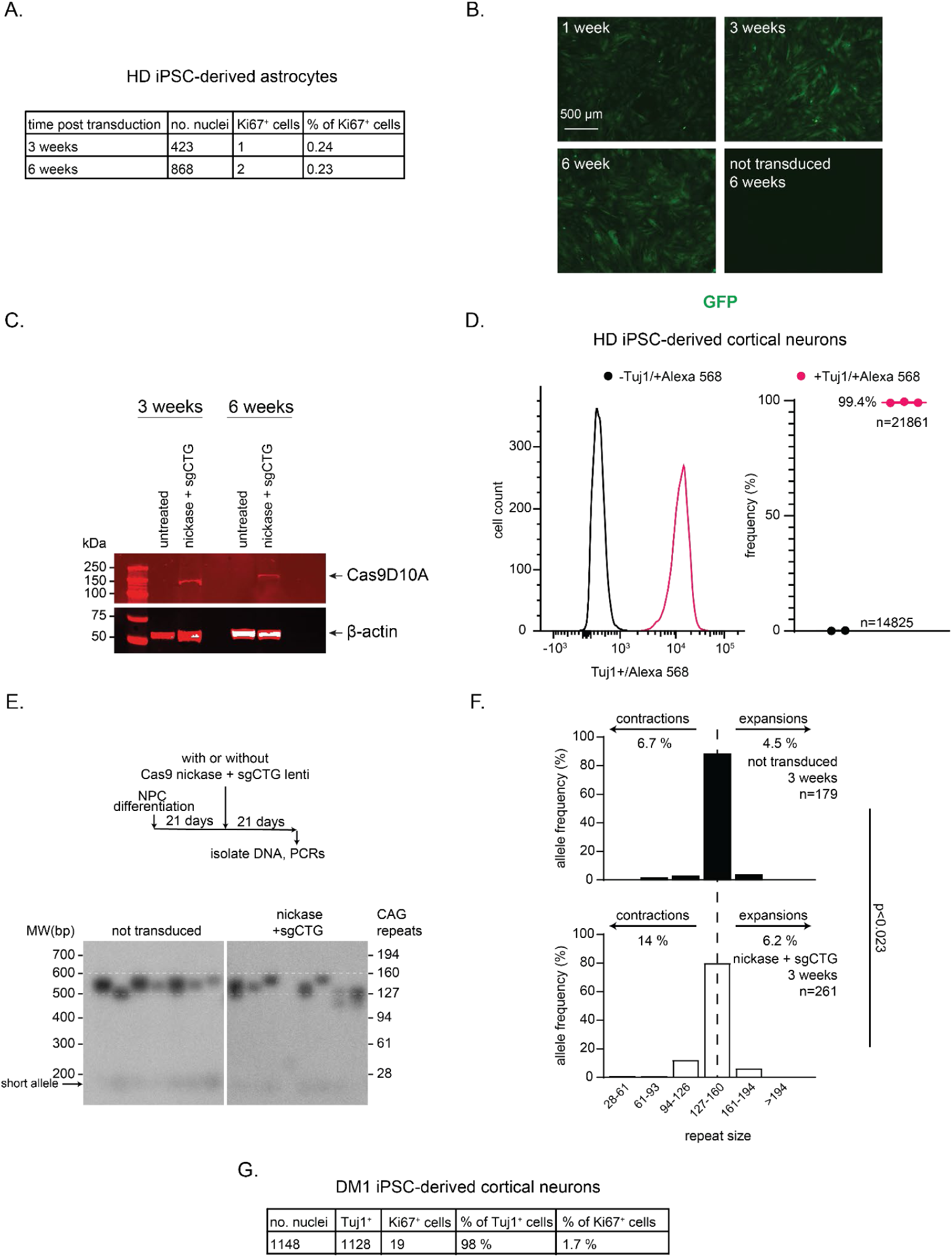
CAG/CTG repeat contractions in HD iPSC-derived astrocytes and neurons. A) HD iPSC-derived astrocyte cultures were sorted using CD44, ensuring that they were all of an astrocytic lineage. We found little replication in these cultures over 21 and 42 days after transduction with Cas9D10A and sgCTG as measured by Ki67 staining. B) Lentivirus-encoded GFP expression on the sgCTG vector over 42 days42 days time course in HD iPCS-derived S100ꞵ^+^ astrocytes (pLV[gRNA]-CMV-EGFP-U6>{sgCTG}). C) Cas9D10A expression in HD iPSC-derived S100ꞵ^+^ astrocytes over 21 and 42 days from (pLenti-EF1alpha-Cas9D10A nickase-Blast). D) Characterisation of the HD iPSC-derived cortical neurons by the expression of Tuj1 as a marker for neurons. Left: Representative flow cytometry histogram of cultures stained with ꞵ-tubulin III (Tuj1; magenta) and only with the secondary antibody (black), suggesting low variability in Tuj1 expression and high quality neuronal cultures Right: quantification of 2 to 3 independent experiments. N = number of cells. E) Timeline and representative small pool PCR blot of HD iPSC-derived neuronal cultures grown in the presence of both Cas9D10A and sgCTG or untreated for 21 days. F) Quantification of the small pool PCRs. P-value determined using a Mann-Whitney U test comparing non-transduced and Cas9D10A + sgCTG. G) DM1 iPSC-derived neuronal cultures stained for Ki67, as a marker of proliferation, and Tuj1, as a marker of neuronal cell type.

**Supplementary Figure 2:**
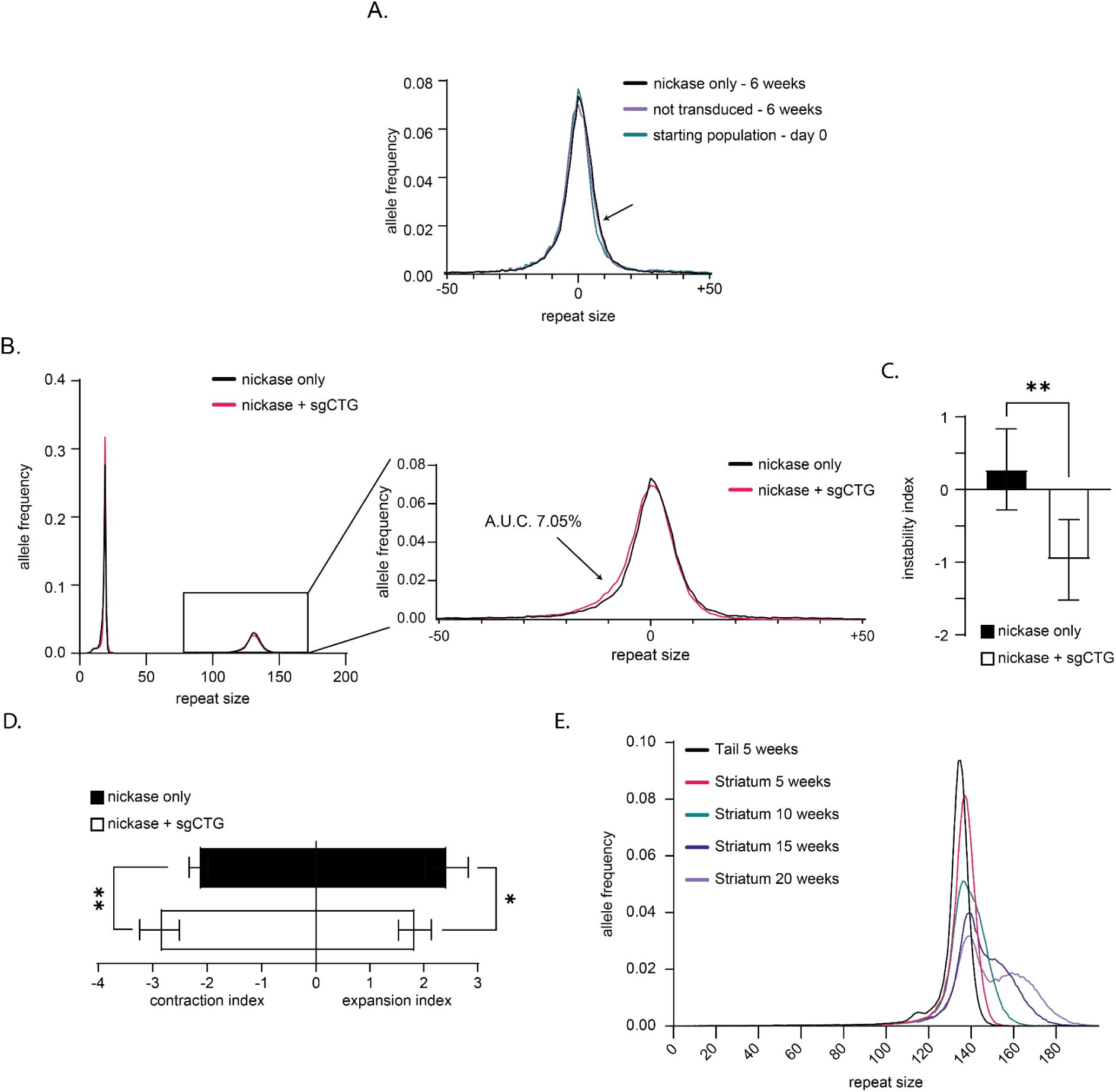
SMRT sequencing reproduces expected patterns of repeat size distribution. A) SMRT sequencing of DNA from HD iPSC-derived cortical neurons showing changes in repeat size between cultures of HD-derived cortical neurons that have not been transduced (n=4), transduced with Cas9D10A only (n=4), compared to the starting repeat size distribution (n=2). Arrow indicates the accumulation of expansions over the 42 days period. The calculated AUC difference for expansions between day 0 and non-transduced cells at 42 days was 5.6 % whereas between D0 Cas9D10A only the AUC was 7.8%. B) SMRT sequencing of DNA from HD iPSC-derived cortical neurons showing both the non-pathogenic and the expanded allele (left) and the change in repeat size between the cells treated with Cas9D10A only versus those treated with both Cas9D10A and sgCTG for 42 days (n=4 per condition). Analysis of the difference in the area under the curve (AUC) shows that 7% of the cells transduced with both Cas9D10A and sgCTG had shorter alleles above those treated with Cas9D10A only. This is comparable to small-pool PCR data. C) Instability indices of the HD iPSC-derived cortical neurons treated with Cas9D10A only (black boxes) and Cas9D10A + sgCTG (white boxes) (n=4 per condition). **:P<0.01 calculated using a Student’s t-test test. D) Same as in (C) but plotting both the expansion and contraction indices. *: P<0.05. E) Representative example time course analysis using SMRT sequencing showing the expected increase in the frequency of longer alleles over several weeks in the R6/1 striatum. These data show that the approach can measure somatic expansions readily.

**Supplementary Figure 3.**
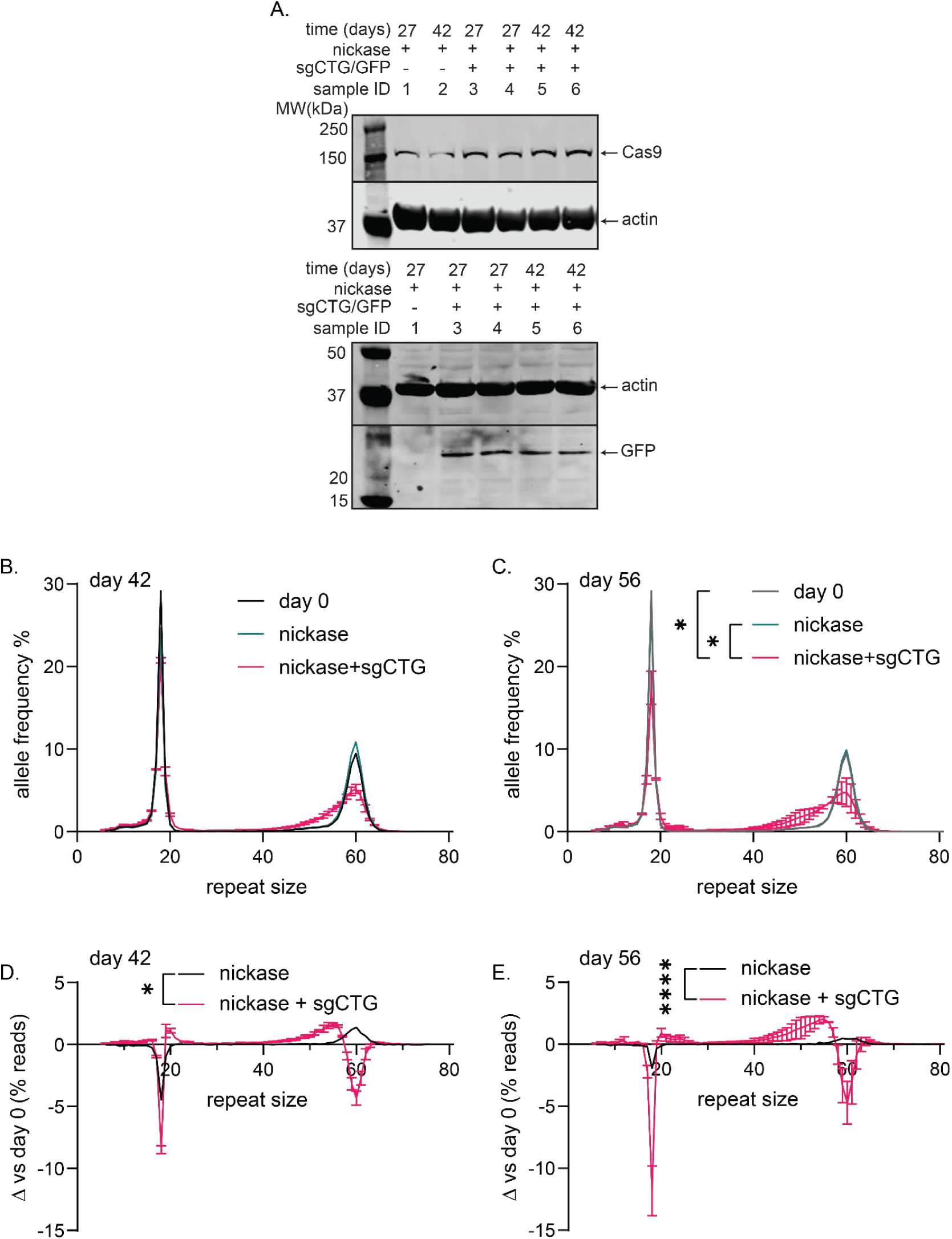
As few as 60 CAG/CTG repeats are a target for contraction by the Cas9D10A. A) Western blot of Cas9D10A and GFP expression from lymphoblastoid cultures transduced with pLenti-EF1alpha-Cas9D10A nickase-Blast and pLV(gRNA)-CMV-eGFP:T2A:Hygro-U6(sgCTG) at the indicated post-infection days. Full blots are found in Supplementary Fig. 21. B) Aggregated graphs of repeat size distribution in lymphoblastoid cultures with Cas9D10A only (magenta) and Cas9D10A + sgCTG (blue) at 42 days of treatment versus day 0 (black). C) Same as B at 56 days after the treatment time point. D-E) Delta plots (see methods) of the data presented in B and C showing the difference at 42 days (D) and 56 days (E) after continuous treatment. This is compared to the starting repeat size distribution at D0. Kolmogorov-Smirnov tests to compare cumulative distributions were used in all graphs. P value <0.05 (*); <0.0001 (****). Data are mean ± SD (D0 n = 1; D42 and D56 Cas9D10A only n=1; D42 and D56 Cas9D10A + sgCTG = 2).

**Supplementary Figure 4.**
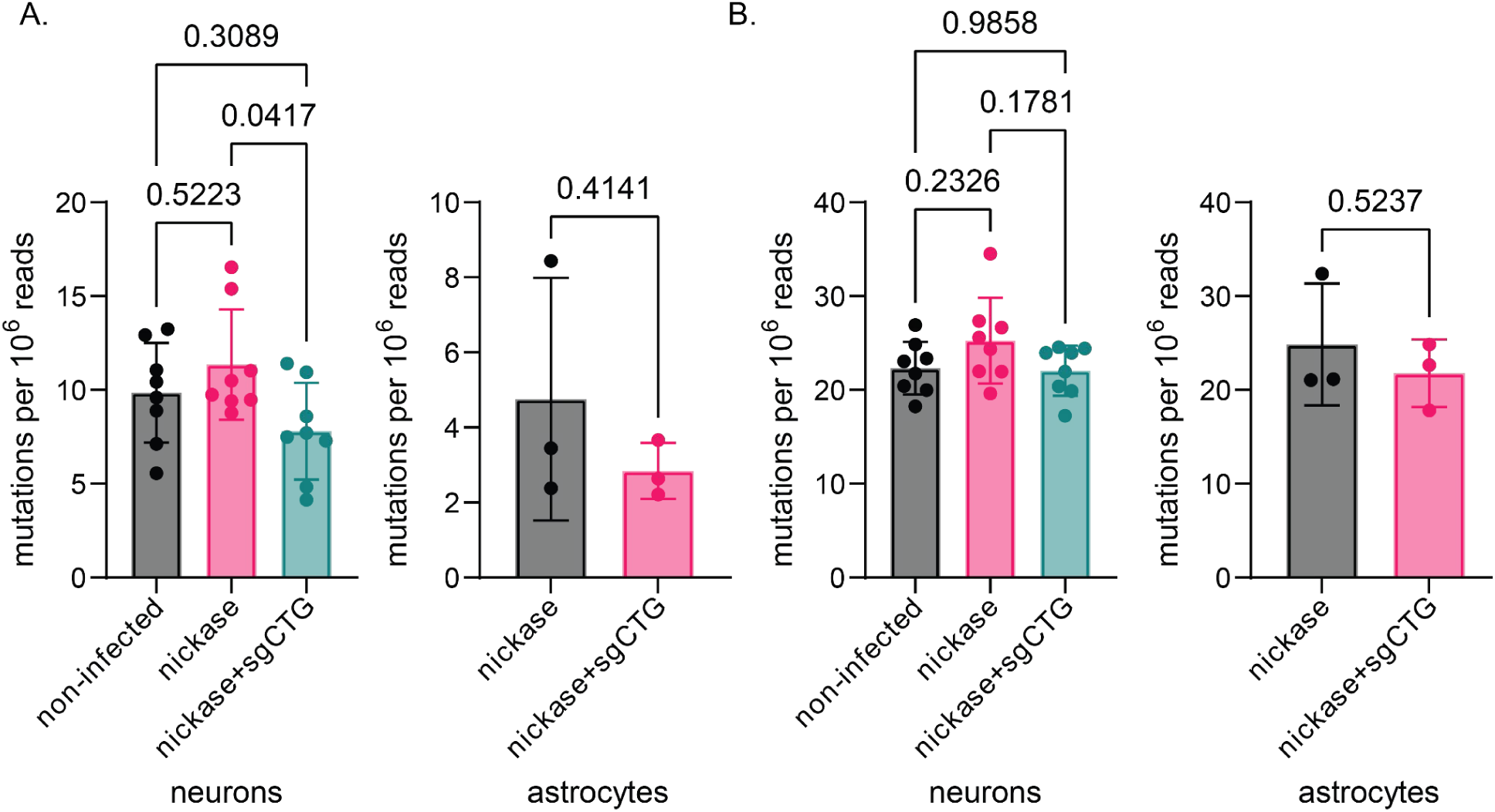
Analysis of mutations in HD iPSC-derived cells. This figure provides a graphical representation of the data listed in Table 1. A) The number of mutations found at the selected 1830 genes were normalised to the total reads per sample and indicated as the number of mutations normalised to 10^6^ reads. B) The number of mutations found genome-wide were normalised to the total reads per sample and indicated as the number of mutations normalised to 10^6^ reads. Differences in normalised mutations between neuronal cultures conditions were calculated by one-way ANOVA with a post-hoc Tukey’s multiple comparison test, except for the whole genome data from astrocytes where we used a Student’s t-test.

**Supplementary Figure 5:**
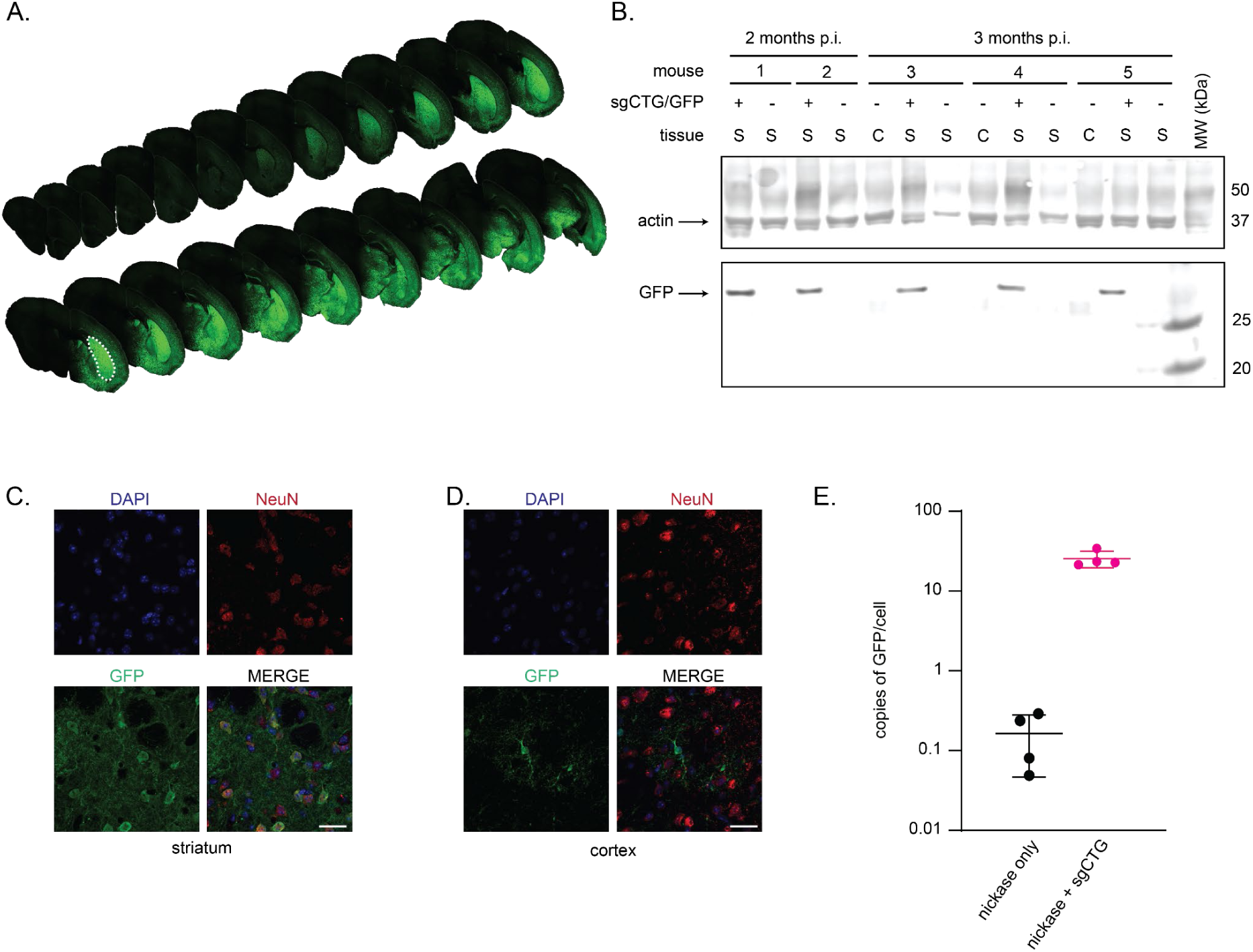
GFP expression in P2-injected mice. A) Coronal sections through the mouse brain showing GFP expression in the injected hemisphere 6 weeks post-injection (p.i.) throughout the striatum (dashed lines) and into much of the cortex. B) GFP and actin expression 2 and 3 months p.i. in striatum (S) and cerebellum (C). C) Representative example of GFP colocalization with NeuN+ neurons in the striatum. D) GFP expression in the cortex did not colocalize with NeuN, but rather the GFP positive cells had an astrocytic morphology. Scale bar = 25 µm. E) qPCR results for the number of GFP DNA found inCas9D10A-only or Cas9D10A + sgCTG-injected striatum, showing that there were, on average, 26 ± 6 copies of the AAV per cell 1 month after injection n=4 per condition.

**Supplementary Figure 6:**
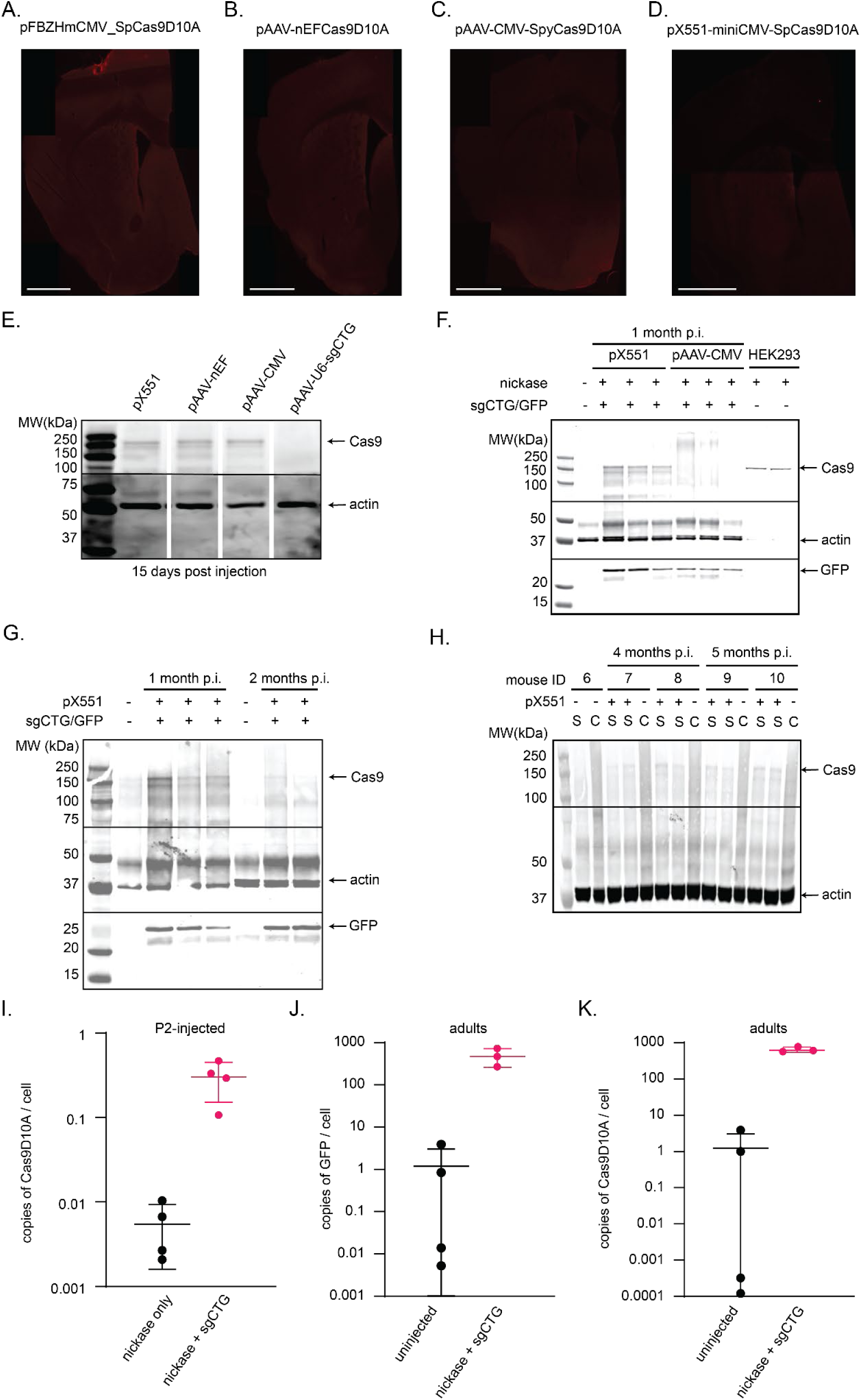
Cas9D10A expression *in vivo*. A) Immunofluorescence 1 month post-injection of Cas9D10A expressed from pFBZHmCMV_SpCas9D10A AAV9 injected into wild type striatum. B) Same as A, but using pAAV-nEFCas9D10A. C) Same as A but using pAAV-CMV-SpyCas9D10A. D) Same as A but using pX551-miniCMV-SpCas9D10A. Scale bars = 1000 µm E) Western blot of Cas9D10A expression from the indicated AAVs in the striatum of adult wild type mice 15 days post-injection (p.i.). This gel was spliced for ease of comparison. Full blots are found in Supplementary Fig. 21. F) Western blot of Cas9D10A expression in the striatum of adult R6/1 mice injected with the indicated AAVs 1 month post-injection. G) Western blot of Cas9D10A expression in the striatum of adult R6/1 mice from an AAV9 packaged using pX551-miniCMV-SpCas9D10A, 1 and 2 months after injection. H) Cas9D10A expression in the striatum of P2-injected R6/1 mice using an AAV9 packaged from pX551-miniCMV-SpCas9D10A, 4 and 5 months post-injection. I) qPCR data quantifying the number of Cas9D10A AAV genomes present compared to the amount of actin. This was done using striatum DNA of P2-injected R6/1 mice 1 month after injection in mice injected with Cas9D10A AAV only (black) or both AAVs (pink). Each dot is a different mouse, n=4 per condition. J) Same as (I), but mice were injected at 3.5 months of age and analysed one month later for the GFP AAV. K) Same as J, but the data shown is for the number of Cas9D10A AAV DNA copies (Number of animals: uninjected n=4; Cas9D10A + sgCTG n=3).

**Supplementary Figure 7.**
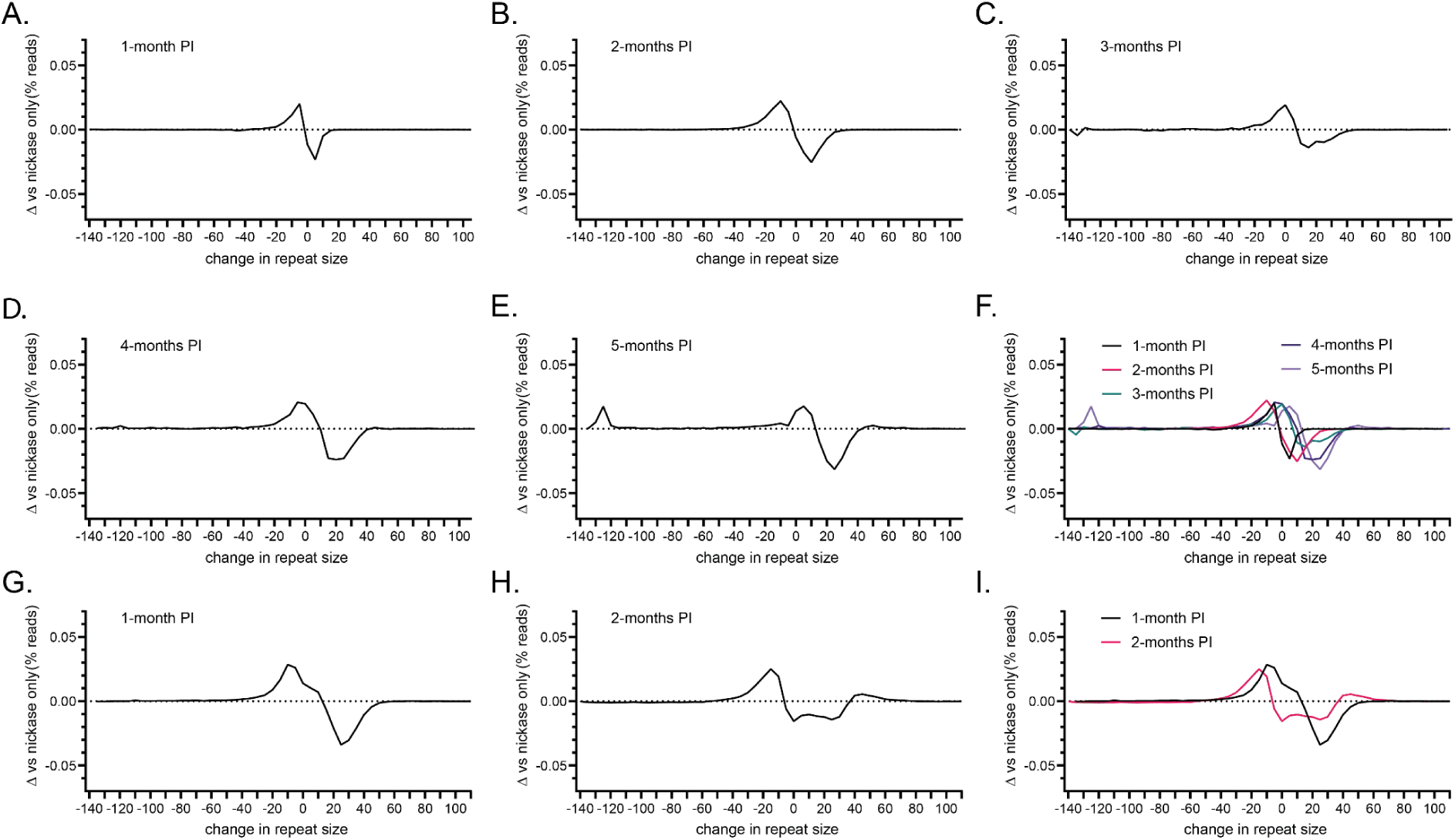
Nickase treatment induces contractions that accumulate over time. A) Aggregated delta plot of repeat size distribution in hemispheres of P2 R6/1 mice injected with Cas9D10A + sgCTG subtracting Cas9D10A only repeat size distribution (see methods) at 1 month, B) 2 months, C) 3 months, D) 4 months and E) 5 months. F) Combined time points to show changes over time. Delta-Plot comparing 1M vs 4M and 1vs 5M PI were compared using an unpaired, nonparametric Kolmogorov-Smirnov test. P value <0.0001 (****). G) Aggregated delta plot of repeat size distribution in hemispheres of adult R6/1 mice injected with Cas9D10A + sgCTG subtracting noninjected repeat size distribution (see methods) at 1 month and H) 2 months post-injection. I) Combined time points to show changes over time. Delta plots of 1 month and 2 months post-injection were compared using an unpaired, nonparametric Kolmogorov-Smirnov test. P value <0.002 (**).

**Supplementary Figure 8.**
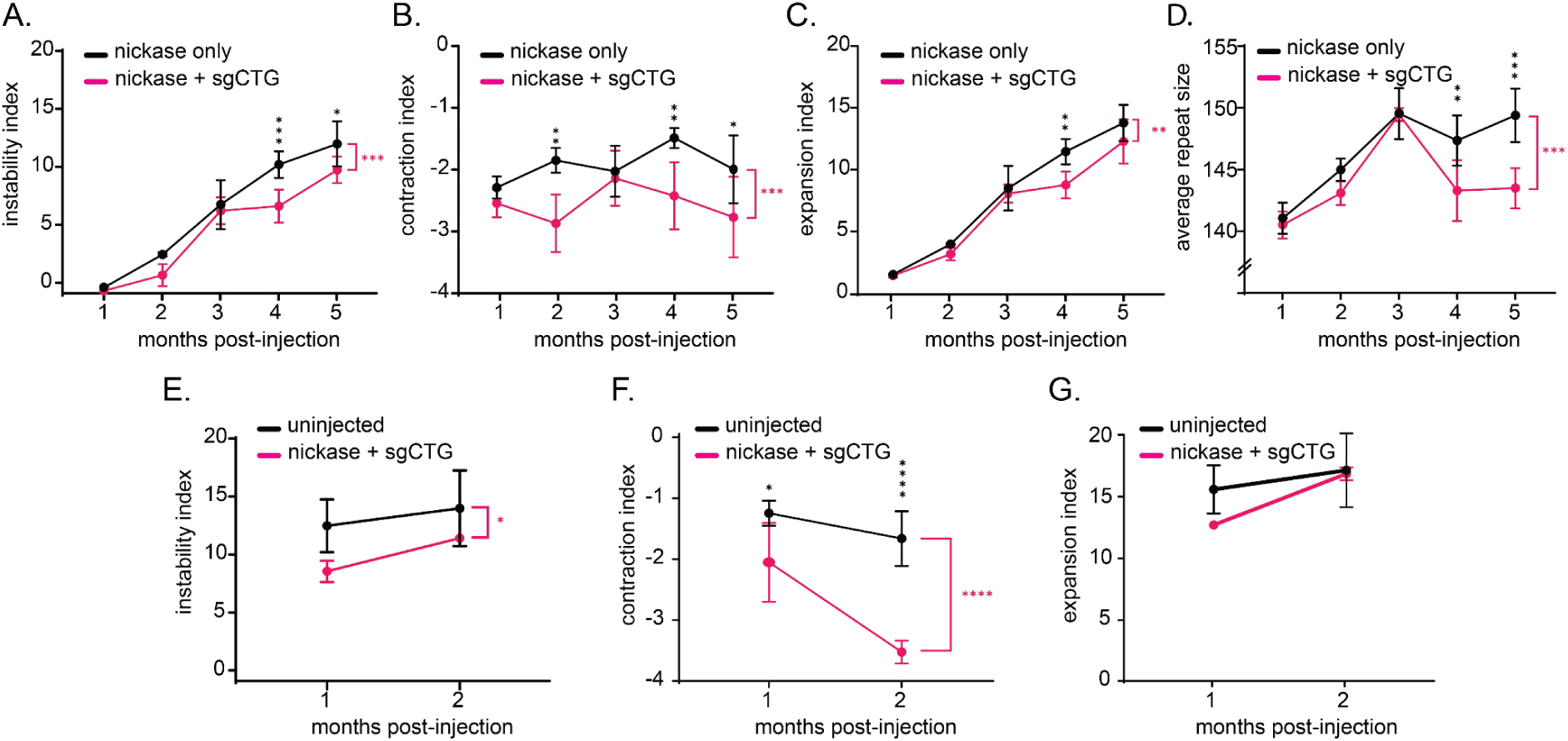
CRISPR-CasD10A reduces instability and increases contraction index. A) Instability index of P2 injected mice treated with both Cas9D10A and sgCTG (magenta) versus Cas9D10A only (black). A two-way ANOVA showed an effect of the treatment (P<0.001, magenta asterisks), with a post-hoc Tukey’s multiple comparison test showing significant differences 4 and 5 months post-injection (*P<0.05, ***P<0.001, black asterisks). B) The same data as in A, but showing only the contraction index (more negative values indicating more contractions). *P<0.05, **P<0.01, ***P<0.001. C) Same as in F but only showing the expansion index.**P<0.01. D) The average repeat size over time. **P<0.01, ***P<0.001 using a two-way ANOVA. Error bars represent standard deviation. Number of animals: Cas9D10A only; 1 month n=4, 2 months n=3; 3 months n=3; 4 months n=4; 5 months n=3; Cas9D10A + sgCTG; 1 month n=4, 2 months n=3; 3 months n=3; 4 months n=4; 5 months n=3). E) Instability index of the adult mice treated with both Cas9D10A and sgCTG (magenta) versus uninjected (black). A two-way ANOVA showed an effect of the treatment *P<0.05. F) The same data as E, but showing only the contraction index (more negative values indicating more/larger contractions). Magenta asterisks: Two way ANOVA, ****P<0.0001, post-hoc Tukey’s multiple comparison test at individual time points (black asterisks) *P<0.05, ****P<0.0001. G) Same as in E but only showing the expansion index. Error bars represent ± standard deviation. Number of animals: uninjected 1 month post-injection n=4, 2 months post-injection n=5; Cas9D10A + sgCTG; 1 month post-injection n=3, 2 months post-injection n=3.

**Supplementary Figure 9:**
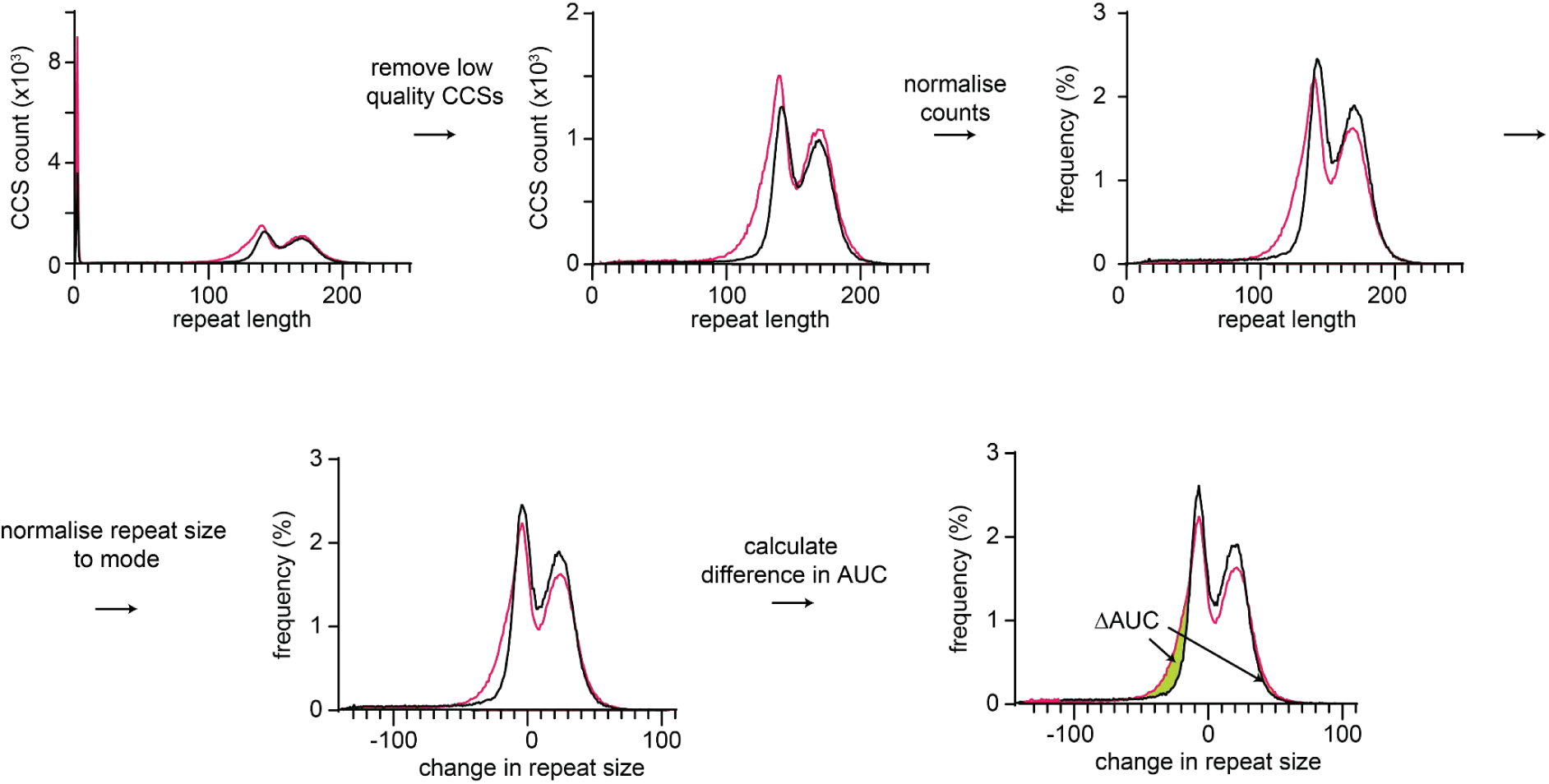
Explanation of how the difference in the area under the curve was calculated. This is an example dataset. Repeat Detector is used to determine repeat sizes. Then repeat size between 0 and 5 CAGs are removed as they are low quality reads that do not align to the *HTT* locus and truncated PCR products. Then the graphs are normalised from CCS counts and for the modal repeat size of each sample. This prevents taking small expansions in the untreated samples to inflate the number of contractions. The difference in the area under the curve is calculated for both shorter and longer alleles.

**Supplementary Figure 10.**
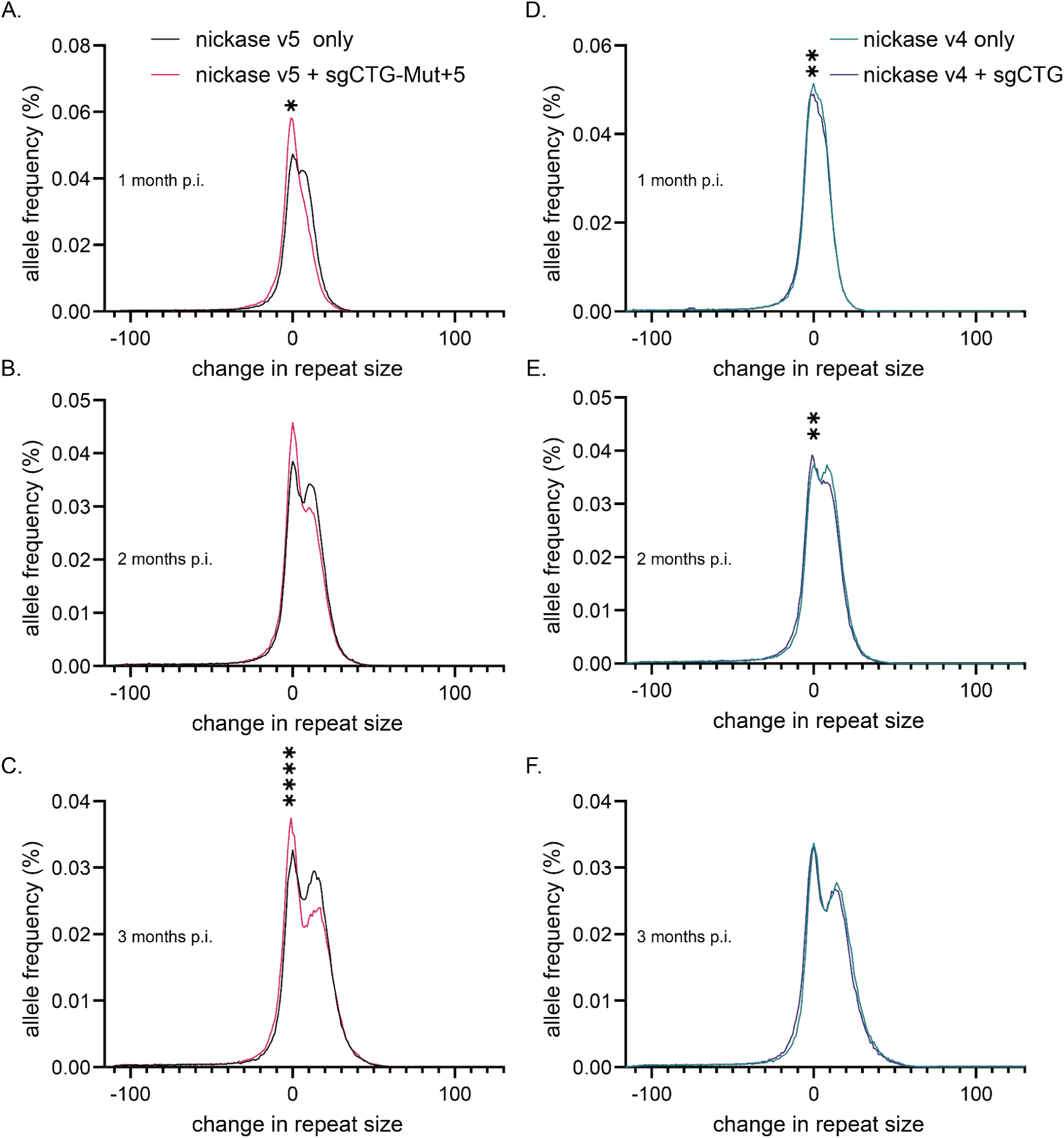
Instability of CAG repeats in adult-injected mice over time.A) Aggregated graphs of repeat size distribution in the striatum of adult R6/1 mice injected with the Cas9D10A v5 only (black) compared to striata injected with Cas9D10A v5 + sgCTG-Mut+5 (magenta) 1 month post-injection. B) Same as A but 2 and C) 3 months-post-injection. D) Aggregated graphs of repeat size distribution in the striatum of adult R6/1 mice injected with the Cas9D10A v4 only (green) compared to striata injected with Cas9D10A v4 + sgCTG (blue) 1 month post-injection.E) Same as D but 2 and F) 3 months-post-injection. Note that the graphs are normalized to the modal peak size found in each mouse (see methods). T test, paired nonparametric test Wilcoxon matched-pairs signed rank test to compare cumulative distributions (*P value <0.05; **P value <0.01; ****P value <0.0001). Number of animals: Cas9D10A v5 only; 1 month n=3, 2 months n=3; 3 months n=3; Cas9D10A v5 + sgCTG-Mut+5; 1 month n=3, 2 months n=3; 3 months n=3); Cas9D10A v4 only; 1 month n=2, 2 months n=3; 3 months n=3; Cas9D10Av4 + sgCTG-Mut+5; 1 month n=2, 2 months n=3; 3 months n=3).

**Supplementary Figure 11.**
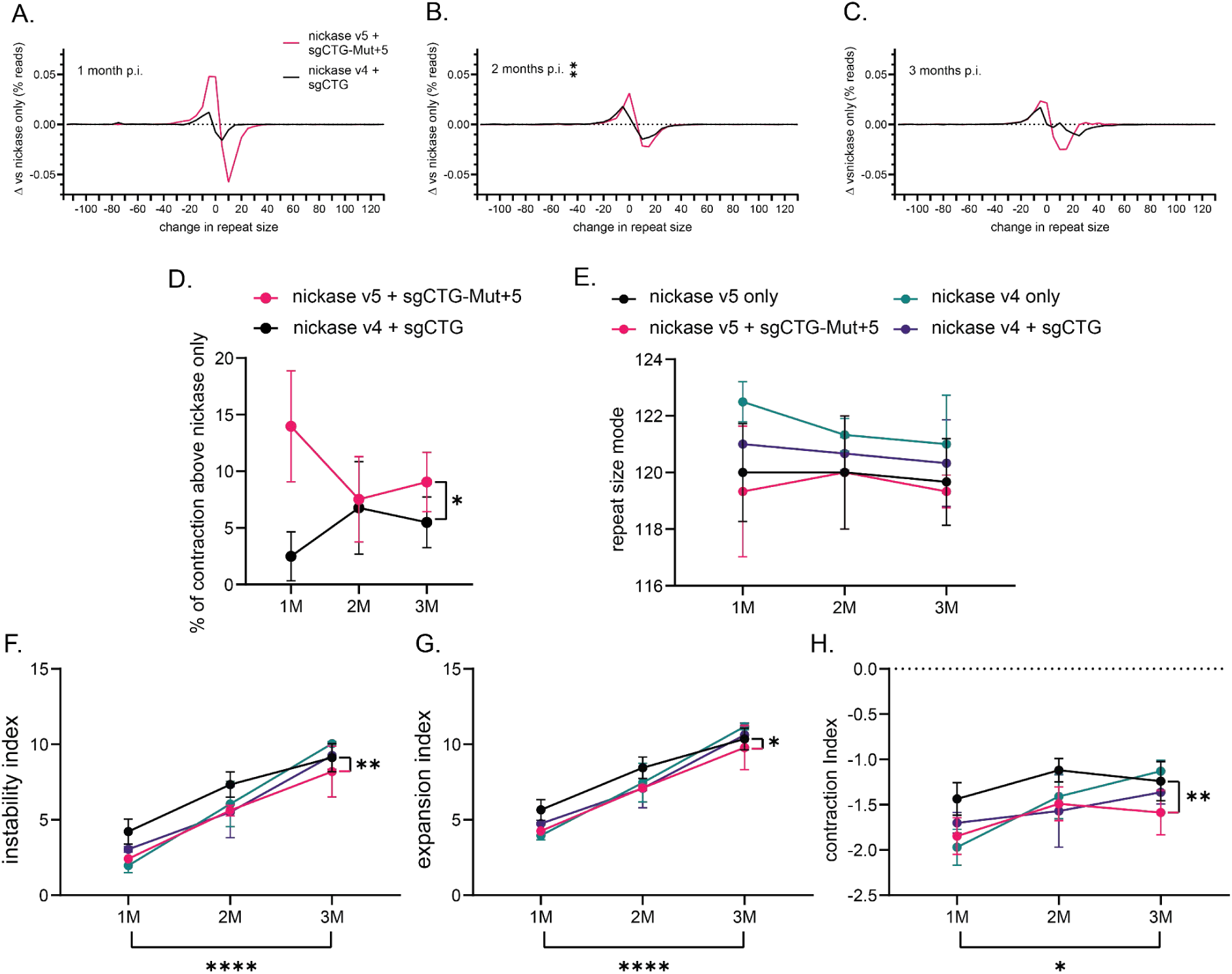
Optimizing the AAV cargos. 2-month-old R6/1 mice were injected in the striatum with the Cas9D10A v5 + sgCTG-Mut+5 ratio 1:2 or Cas9 v4 + sgCTG ratio 1:1. The other striatum received the respective Cas9D10A AAV only. A) Delta plot analysis revealed significant differences in the contractions induced between treatments at 1, 2 (B) or 3 (C) months post-injection. Unpaired nonparametric Kolmogorov-Smirnov test was used to compare cumulative distributions (**P<0.01) D) Area under the curve comparing the Cas9D10A v5 + sgCTG-Mut+5 treatment with Cas9D10A v5 alone (*P<0.05 using a one-way ANOVA). E) Analysis of the modal CAG repeat size showed no difference between groups by 2-way ANOVA test. F) Instability index of the mice treated versus Cas9 only. A two-way ANOVA showed an effect of the treatment reducing repeats only in the Cas9v5 and sgCTG-Mut+5 treated mice. G) Same as in F, but showing only the expansion index and H) contraction index. D-H) Cohorts show an effect of the timing as expected by the two-way ANOVA analysis (P<0.0001). Two way ANOVA, *P<0.05, **P<0.0, ****P<0.0001, post-hoc Tukey’s multiple comparison test. Error bars represent ± standard deviation. Number of animals: indicated in Supplementary Fig. 10.

**Supplementary Figure 12.**
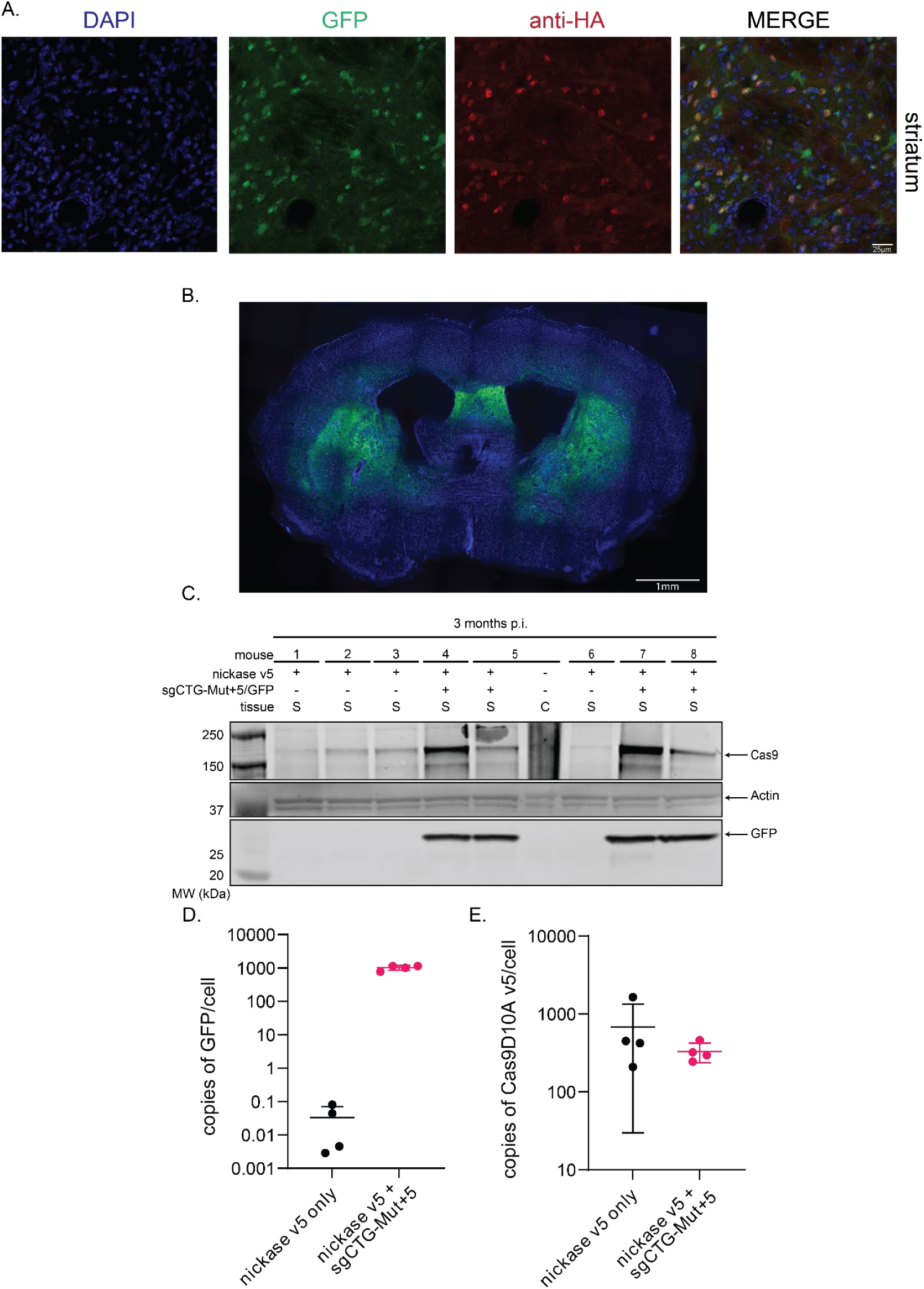
Cas9 nickase optimized and scAAV-sgCTG-Mut+5 express in mouse brain. A) Representative image of 2-months-old wild type injected mouse sacrificed 15 days PI. Immunofluorescence against Cas9OPTI with the anti-HA antibody reveal co-expression with GFP express by the AAV genome. B) 3-months post-injected brain analysis showed that a single injection of AAV9 resulted in widespread and efficient transduction of GFP detected by IF with anti-GFP antibody C) Western blot shows high levels of Cas9D10A expression as well as GFP 3-month PI. Full blots are found in Supplementary Fig. 21. D) DNA extracted from striatal samples were used to determine the number of Cas9OPTi (D) and GFP (E) AAV genome copies by qPCR in control and treated mouse samples. Data are mean ± SD (n = 4 animals per condition).

**Supplementary Figure 13.**
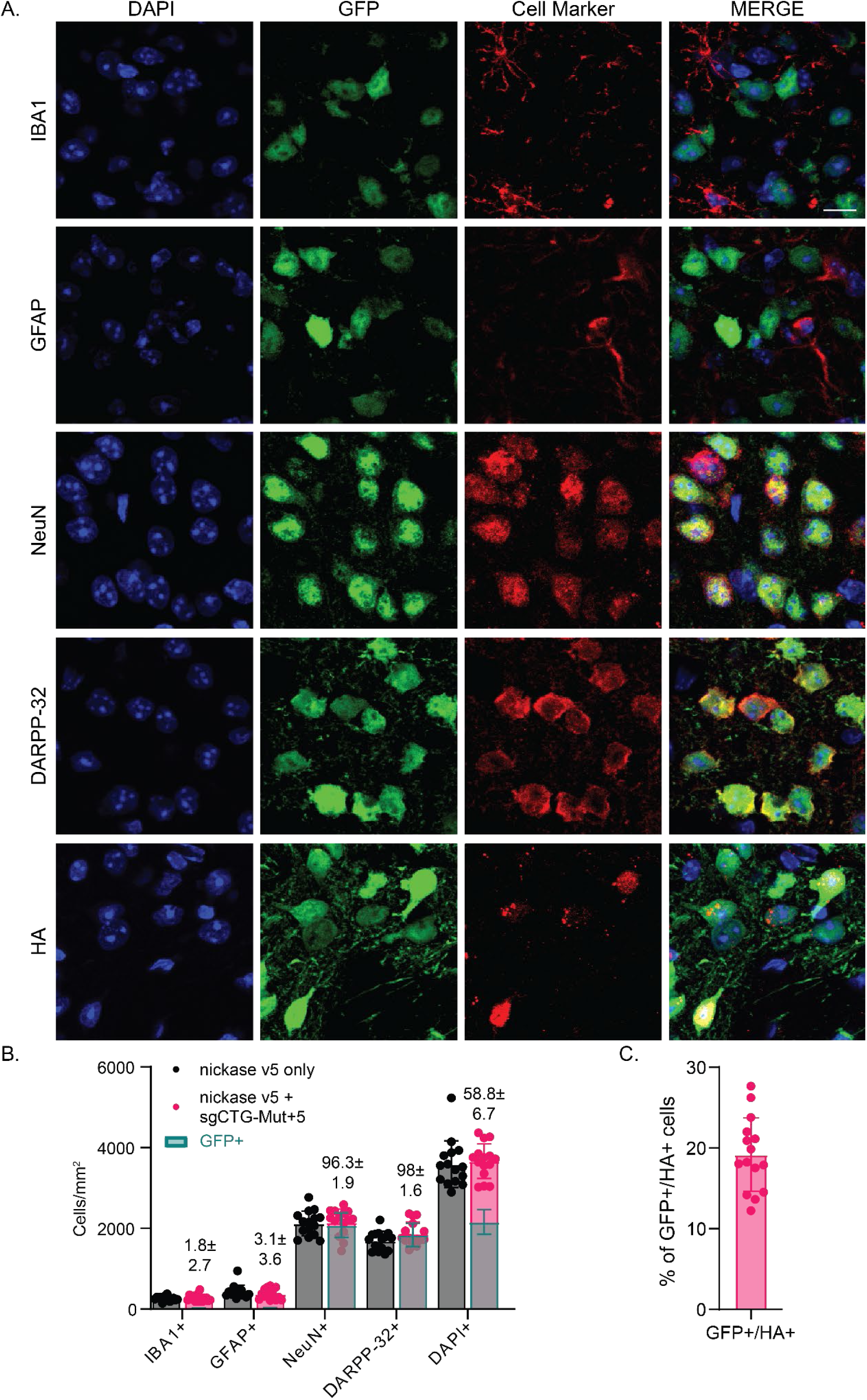
Infection rates of sgCTG-Mut+5/GFP and Cas9D10A v5 AAVs injected in the mouse striatum. A) Representative images of 5-months-old R6/1+ injected with the nickase v5 and sgCTG-Mut+5/GFP mouse sacrificed 3 months post-injection. Immunofluorescence with the anti-GFP antibody reveals co-expression with other cell type markers: Iba1 (microglia), GFAP (astrocytes), NeuN (neurons), DARPP-32 (medium spiny neurons). Cas9D10A was detected using an anti-HA antibody. Scale bar: 10µm. B) Quantification of striatal cells positive for a given marker in R6/1 mice injected with the Cas9D10A only compared with those injected with the Cas9D10A and sgCTG-Mut+5 show no significant differences in the total cell types or in total cell per mm^2^ (DAPI+cells). Quantification of transduced cells with the sgCTG-Mut+5/GFP (green bars) shows a tropism for neurons of the AAV9. Percentages indicate the proportion of GFP+ for each cell marker. C) Quantification of striatal cells positive for GFP that co-express the nickase detected with the anti-HA antibody (19.2±4.6). Data are mean ± SD (n = 3 animals per group, average of 298 cells per image from 5 striatal fields at different antero-posterior and dorso-ventral levels were assessed per mouse).

**Supplementary Figure 14.**
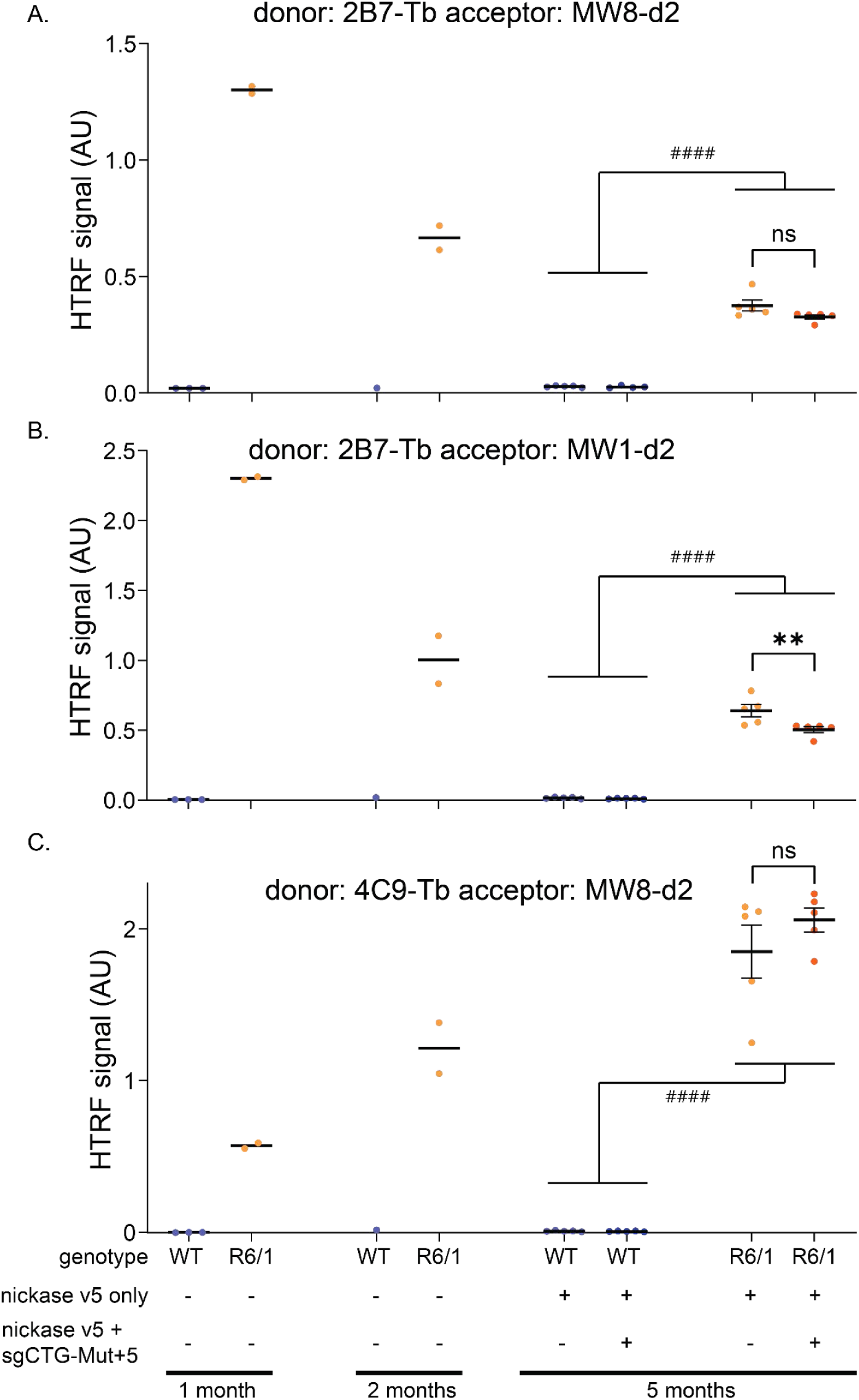
mHTT expression analysis by HTRF assay over the time. Cortical tissues from 1 and 2-months-old control wild type (WT) and R6/1 were used to validate the assay in the R6/1 mouse HD model. The expression of mHTT was compared with striatal tissue samples from experimental animals in Fig. 3b-d. A) 4C9-Tb with MW8-d2 was used to track aggregated mutant HTT, B) 2B7-Tb with MW8d2 were used to track changes in soluble mutant HTT and C) 2B7-Tb with MW1-d2 were used for polyglutamine size assay by HTRF in striatal lysates from R6/1 mice at 1, 2 and 5 months of age. Data are mean ± SD (n 1–5 animals per group). 2-way ANOVA test was run in 5-months old animals (see Supplementary Table 4) with a post-hoc Tukey’s multiple comparison test showing significant differences between WT and R6/1 animals (^####^P<0.0001, black hash) and between R6/1 injected with Cas9D10A only versus dual AAVs injection (**P<0.01, black asterisks).

**Supplementary Figure 15.**
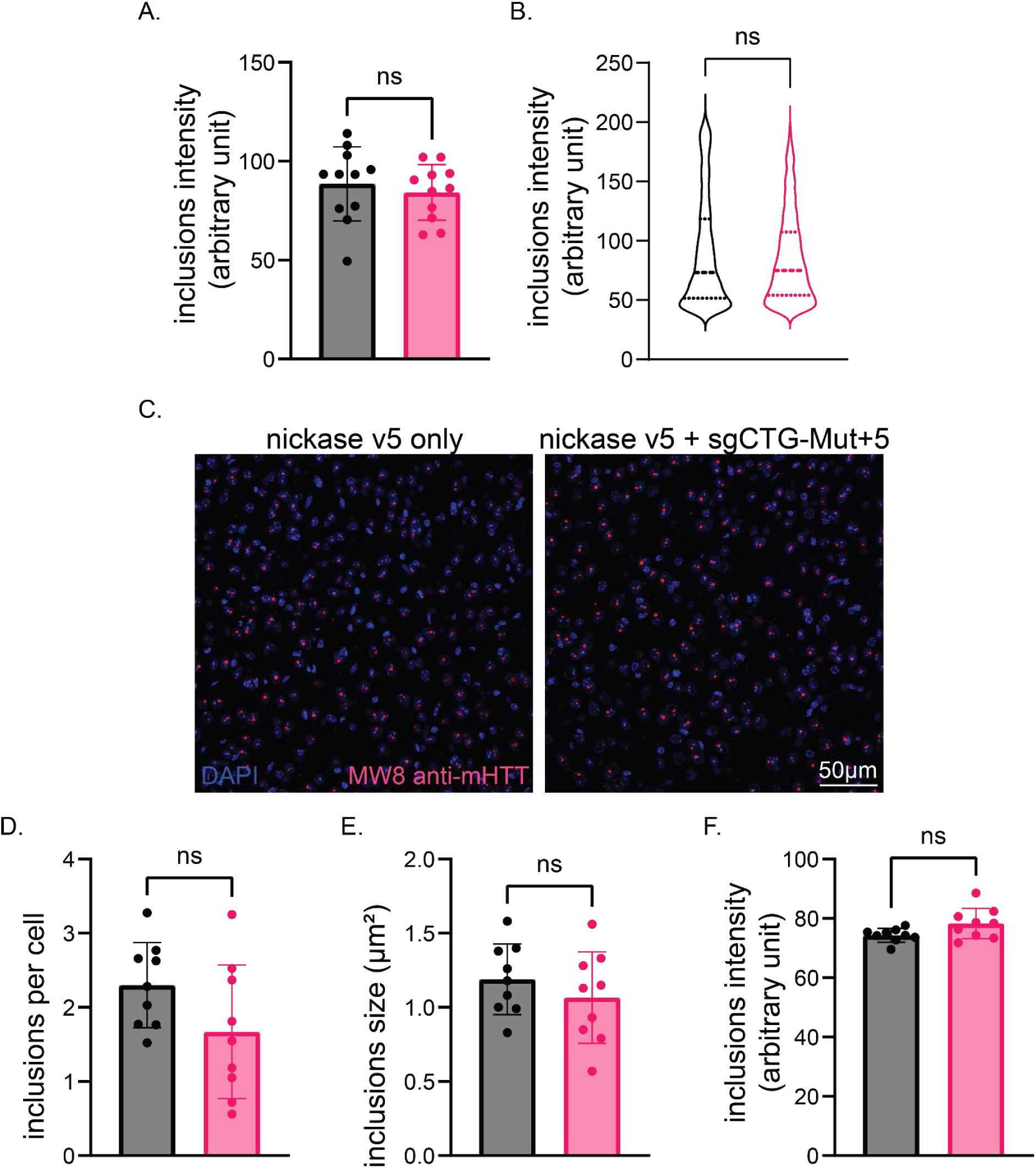
Cas9 nickase brain injection improves mHTT aggregation pathology in striatal areas but not in uninjected cortical areas. A) Average intensity of inclusions between groups and B) intensity distribution shows no differences between animal groups, suggesting that the remaining foci, although smaller, remain inclusions (4009 and 2303 inclusions assessed in total in Cas9 only and Cas9 + sgCTG injected animals respectively). C) Representative z-projection images of layer 2-3 cortical areas from the same preparations as in Fig. 3e of 5-months-old R6/1 animal brains sacrificed 3 months post-injection of the Cas9D10A only or Cas9D10A and sgCTG and stained with DAPI (blue) and mutant polyglutamine inclusions (red). mHTT analysis in cortical areas revealed no significant differences between R6/1 animal groups in number of inclusions per cell (D), inclusions sizes (E) or intensity (F). Data are mean ± SD from a Student t-test. (n = 3-4 images per mouse, ∼900-1600 cells assessed total per mouse, with n = 3 mice per condition).

**Supplementary Figure 16.**
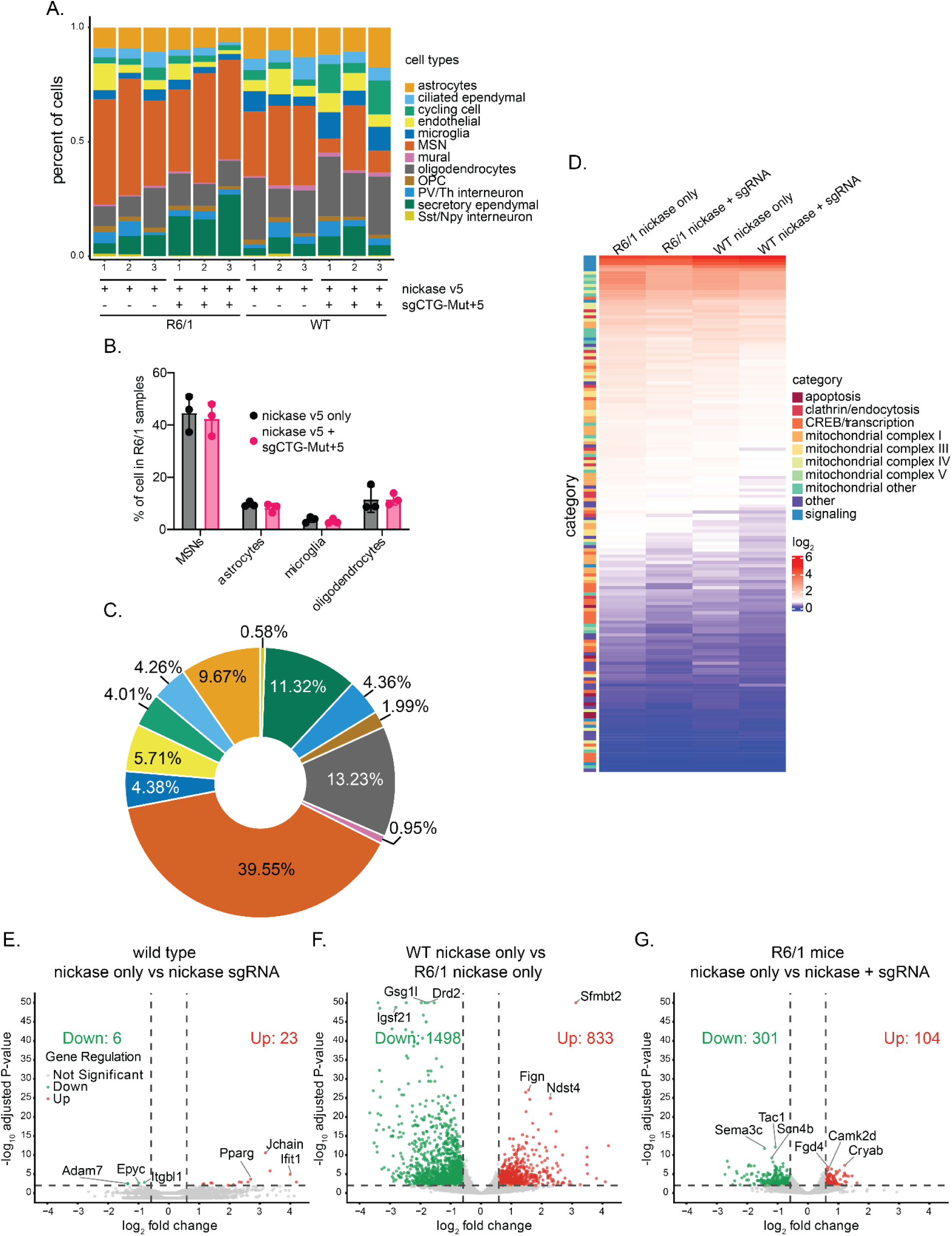
CRISPR injection mitigates disease-associated transcription pathology in HD medium spiny neurons and pseudo-bulk differential expression analysis. A) Cell type frequency distribution found in striatal tissue per sample condition. B) Cell types distribution in R6/1 mice injected with the Cas9D10A only compared with those injected with the Cas9D10A and sgCTG-Mut+5. C) Cell type proportion distribution for all samples combined. D) Heatmap of normalized gene expression showing a select subset of DEGs in HD-relevant pathways. The DEGs (rows) are colour-coded on the right by the direction of gene expression in the specified region (right columns). Horizontal bars in the left column are colour-coded for pathway gene related categories in HD. E) Volcano plot for differentially expressed genes (DEGs) in medium spiny neurons (MSNs) with a pseudo-bulk differential expression analysis between wild type animals injected with Cas9D10A v5 only versus Cas9D10A v5 + sgCTG-Mut+5, F) WT versus R6/1 animals injected with Cas9D10A v5 only and G) CRISPR-treated R6/1 MSNs relative to Cas9D10A v5-injected R6/1. Number of animals: WT Cas9D10A v5 only n=3, WT Cas9D10A v5 + sgCTG-Mut+5 n=3, R6/1 Cas9D10A v5 only n=3, R6/1 Cas9D10A v5 + sgCTG-Mut+5 n=3.

**Supplementary Figure 17.**
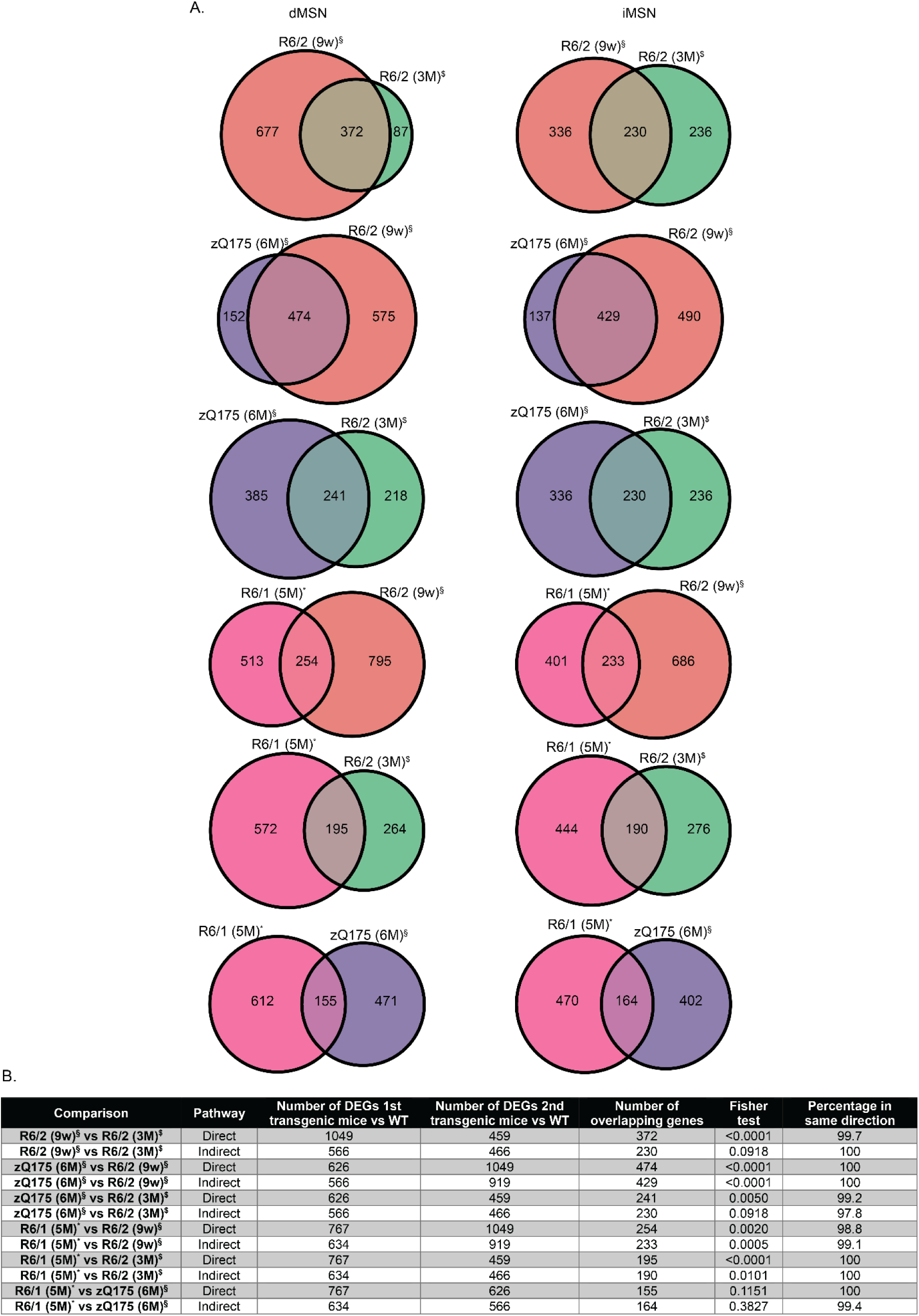
Differential expression analysis in medium spiny neurons (MSNs) from R6/1 mice injected with the Cas9D10A only show similar alterations as other HD animal models. A) Venn diagrams with the number of genes altered compared to wild type animals. Diagrams show the comparison of D1+ MSNs of the direct pathway (dMSNs) and D2+ MSNs of the indirect pathway (iMSNs) of the indicated mouse models and ages from: *: animals from this study; §: animals from Lee et al^58^. $: animals from Lim et al^92^. B) Table summarizing differentially expressed genes (DEGs) in MSNs from previous Venn diagrams. Fisher’s exact test was run using total DEGs and overlapping genes to assess similarities between animal models.

**Supplementary Figure 18.**
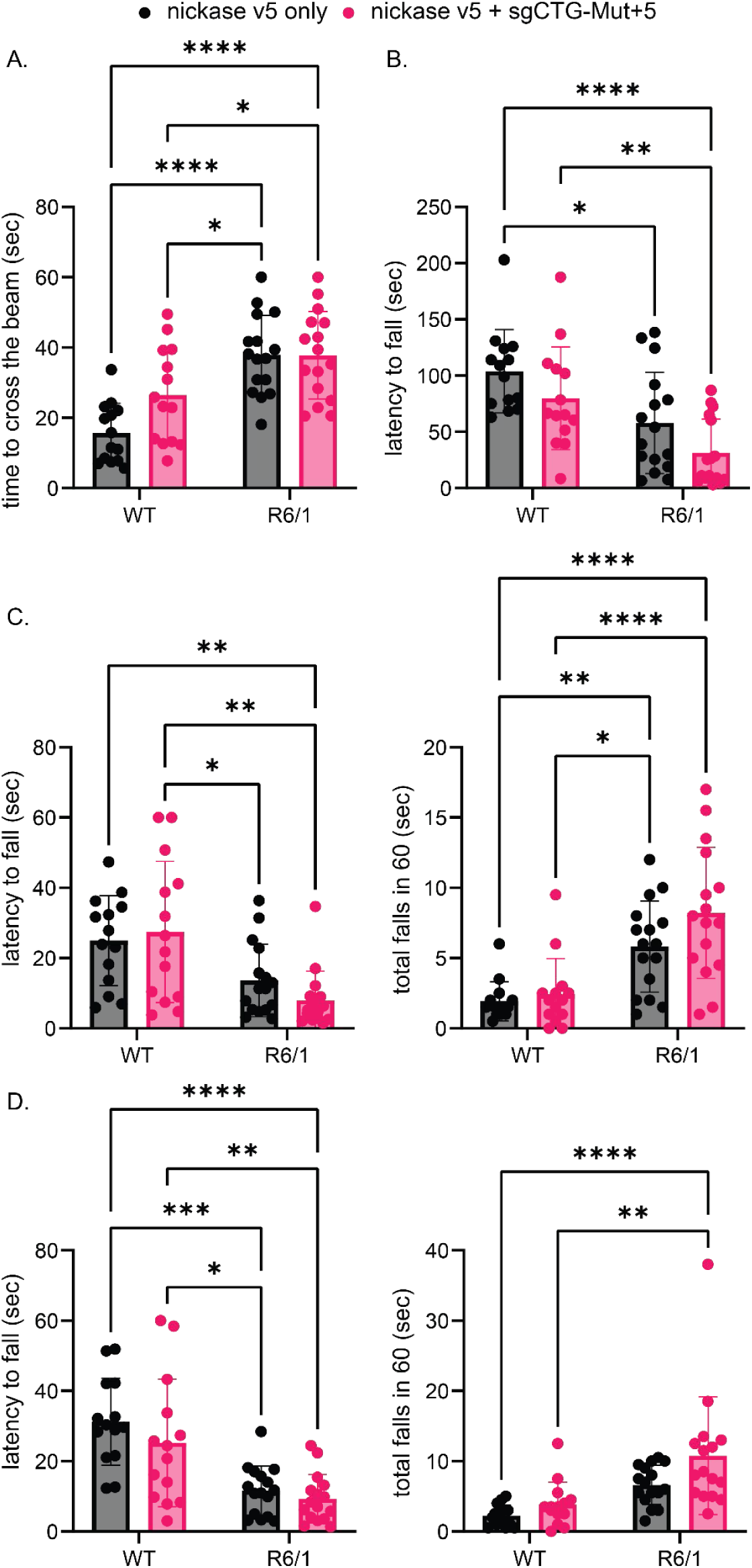
Treated R6/1 animals do not improve in motor coordination tests. A) Average of the timing needed for the animals to cross the beam was measured and no differences induced by the treatment were found. B) We measured the first fall timing in the accelerated rotarod and no improvement after the treatment was found in R6/1. C) First fall timing (left) and total number of falls (right) from a fixed rotarod set at 12 rpm for 3-month-old mice of the indicated genotypes that received bilateral intrastriatal injections of Cas9D10A only and the Cas9D10A and the sgCTG-Mut+5 at 2 months of age. D) Same as C) with the rotarod set at 24rpm. All data are an average of two trials. N of animals=14-16 per group. Statistical analyses are in Supplementary Table 8.

**Supplementary Figure 19.**
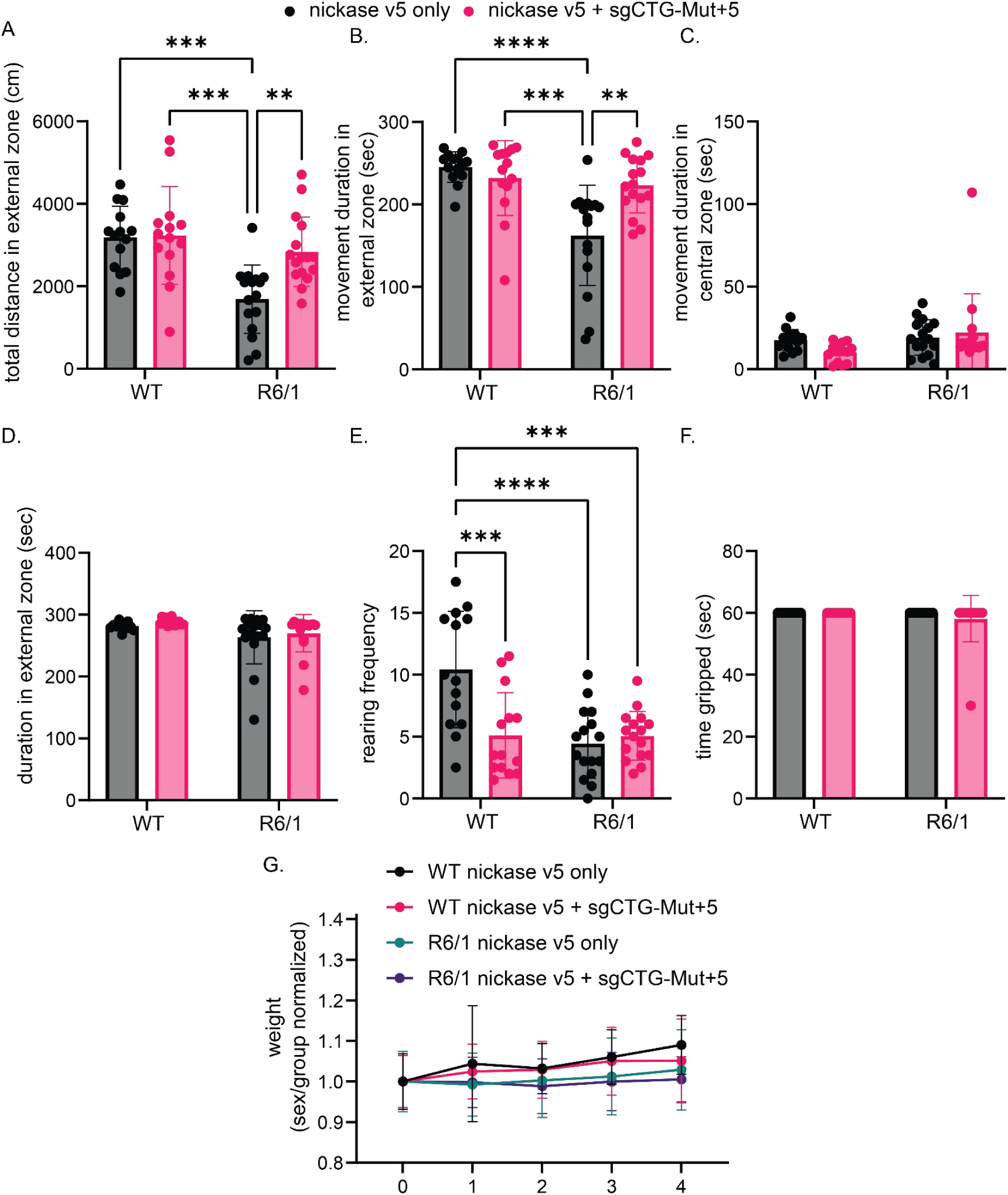
Injection with the CRISPR system does not alter or rescue exploratory and anxiety parameters. A) Average of total distance traveled in the external part of the arena. B) Average time in movement on the external part of the arena. C) Average of the total movement time in the central part of the arena. D) Total movement and stop time in the external part of the arena. E) Exploratory behavioural measured by the rearing frequency shows no differences between R6/1 animal groups but it is reduced in wild type (WT) animals. F) Animal muscular strengths are not affected by the injections. G) Weight of the animals were measured before the injection and every week after. Normalization per gender and groups shows no differences caused by the treatments. N of animals=14-16 per group. Statistical analyses are in Supplementary Table 9.

**Supplementary Figure 20.**
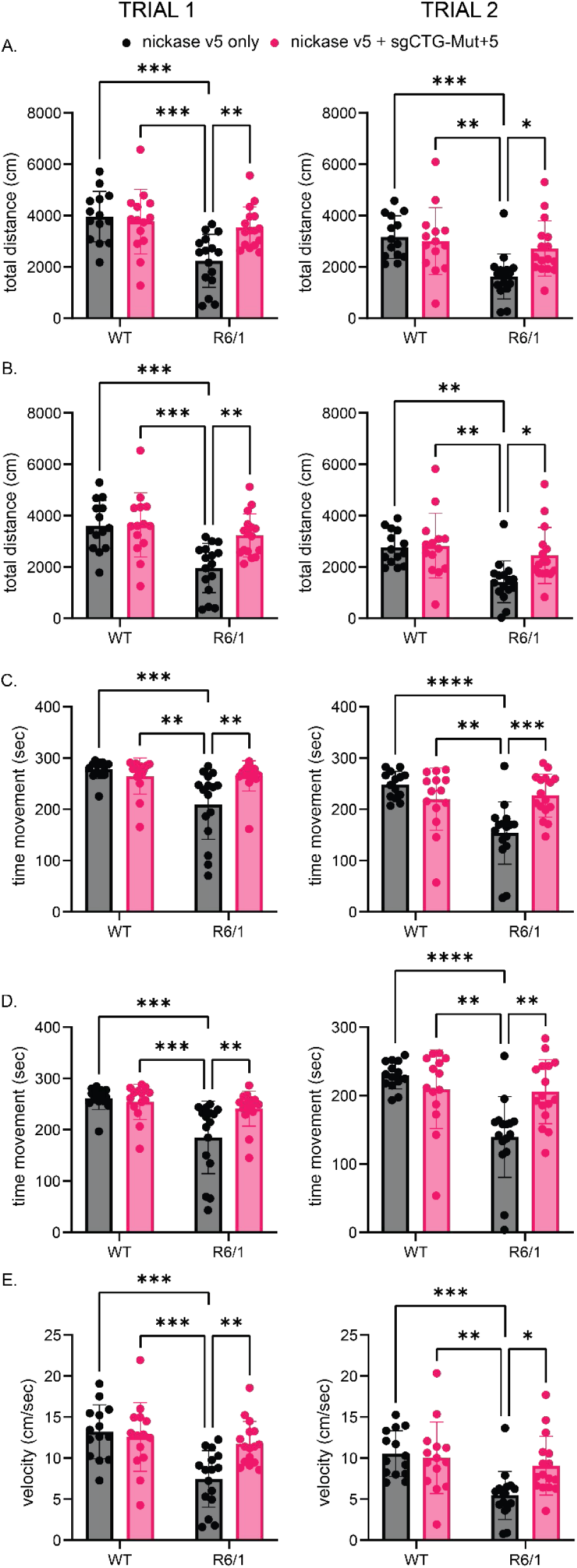
Locomotor activity is stable between Open Field trials. Same data as in Fig. 5c-e and Supplementary Fig. 19ab showing differences between 2 performed open field trials; first trial left column and second trial right column. A) Total distance moved in the 5 min test. B) Same as A but total distance in the external part of the arena. C) Same as A but with the total movement time. D) Same as C but total movement time in the external part of the arena. E) Mean velocity in the total arena. N of animals=14-16 per group. Statistical analyses are in Supplementary Table 10.

**Supplementary Figure 21:**
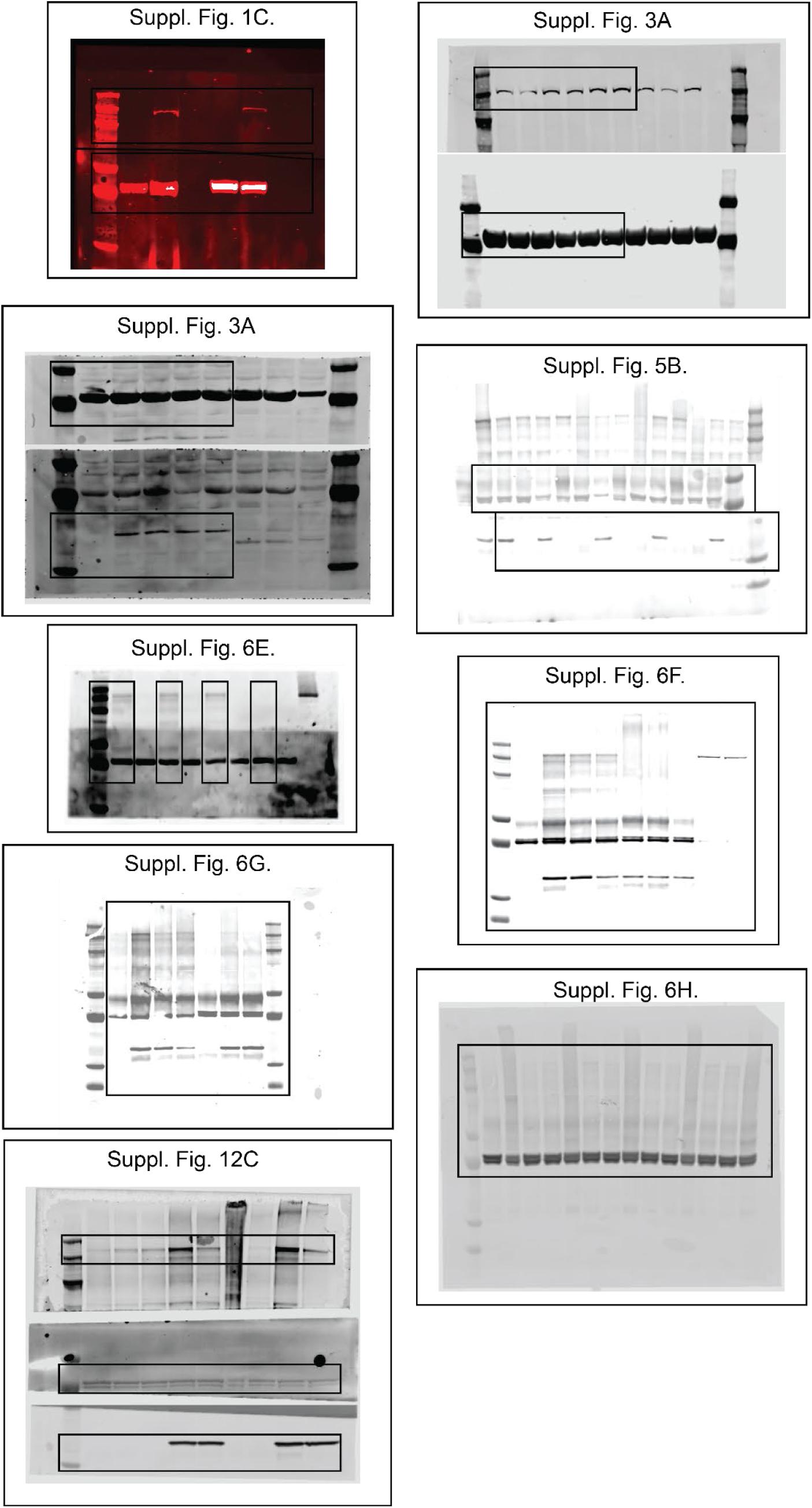
Unaltered full western blots membranes. Black boxes indicate where the blots were cropped.

**Supplementary Figure 22:**
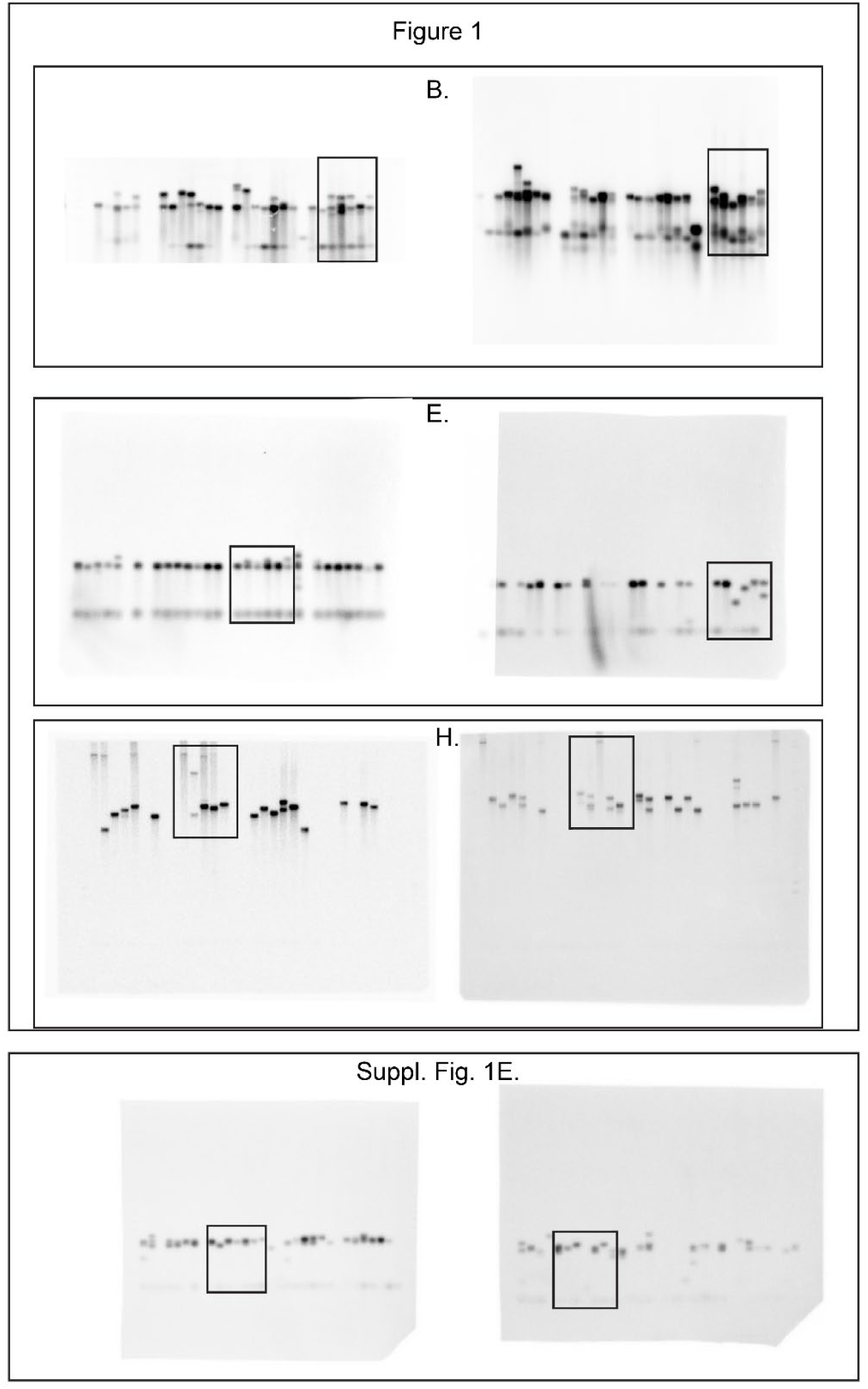
Unaltered full small-pool PCR membranes from Fig. 1beh and Supplementary Fig. 1e. Black boxes indicate where the membranes were cropped.

**Supplementary Table 1:**
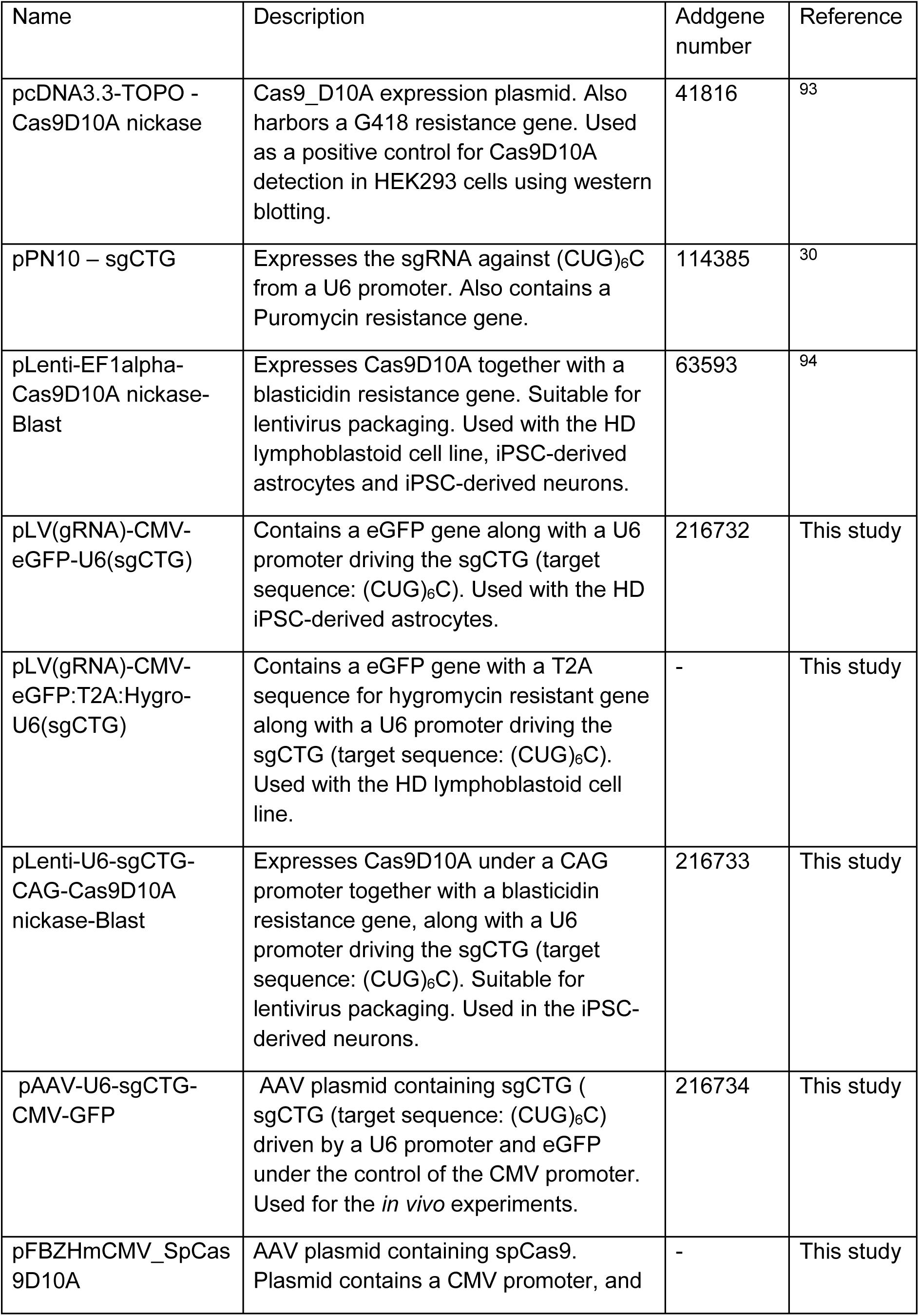
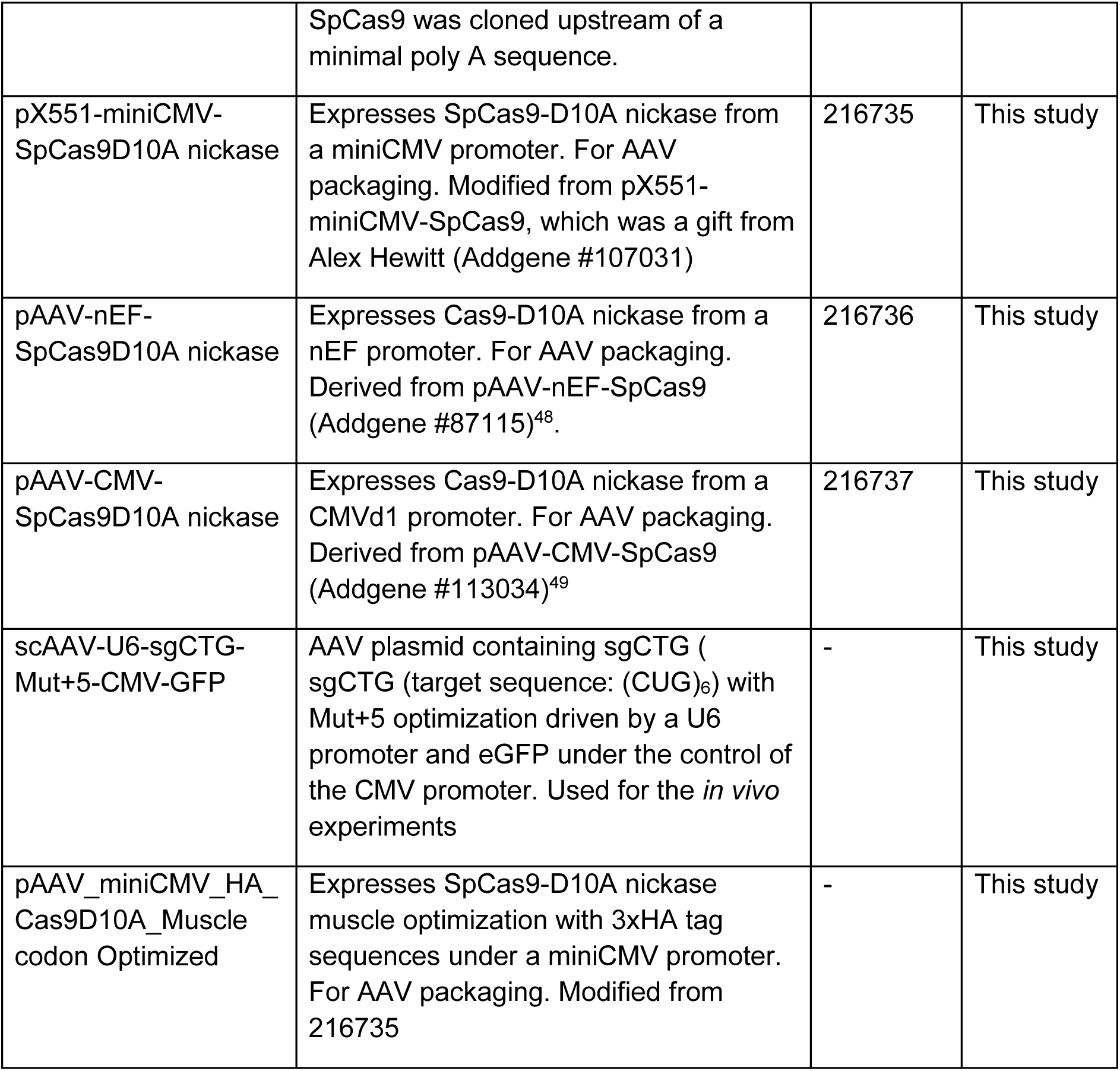
Plasmids used herein.

**Supplementary Table 2:** Whole Genome Sequencing of off-target mutations found in HD iPSc-derived cells and in genes containing a potential off-target site. See Excel sheets attached.

**Supplementary Table 3.** CRISPR system AAV construct versions used in this study.

| Name | Other names | Description | Plasmid in table 3 | Used in figures: | Addgene number | Reference |
| --- | --- | --- | --- | --- | --- | --- |
| Cas9D10A v1 | Nickase v1 | No optimization of spCas indicated | pFBZHMCMV_SpCas9D10A | Supplementary Fig. 6a | - | This study |
| Cas9D10A v2 | Nickase v2 | No optimization of spCas indicated | pX551-miniCMV-SpCas9D10A nickase | Suppl Fig6A-F | 216735 | This study |
| Cas9D10A v3 | Nickase v3 | Optimization of nuclear transport of spCas9. | pAAV-nEF-SpCas9D10A nickase | Suppl Fig6A-E | 216736 | This study |
| Cas9D10A v4 | Nickase v4 | human codon-optimized spCas9 from F Ann Ran et al., 2013 | pAAV-CMV-SpCas9D10A nickase | Fig 2;Suppl Fig5-11 | 216737 | This study |
| Cas9D10A v5 | Nickase v5 | spCas9D10A mouse muscle codon optimization | pAAV_miniCMV_HA_Cas9D10A_Muscle codon Optimized | Fig 3-5;Suppl Fig10-19 | - | This study |
| sgCTG-1 | Guide-1 | Express sgCTG (sgCTG (target sequence: (CUG)<sub>6</sub>C) with an original tracrRNA sequence | pAAV-U6-sgCTG-CMV-GFP | Fig 2;Suppl Fig5-11 | 216734 | This study |
| sgCTG-2 | Guide-2 | sgCTG-1 adapted for higher efficiency following Ying Dang et al., 2015 modification of the "Mut+5". (target sequence: (CUG)<sub>6</sub>) | scAAV-U6-sgCTG-Mut+5-CMV-GFP | Fig 3-5;Suppl Fig10-19 | - | This study |

**Supplementary Table 4:** statistics of HTRF tests Supplementary Figure 14.

|  |  |
| --- | --- |
| **HTRF soluble assay 2B7-tb with MW8-d2** |  |
| **2-way-ANOVA post-hoc Tukey's multiple comparison test** |  |
| Genotype | F (1, 15) = 572.8, P<0.0001 |
| Treatment | F (1, 15) = 3.375, P=0.0861 |
| Genotype X Treatment | F (1, 15) = 2.917, P=0.1083 |
| **HTRF polyglutamine assay 2B7-tb with MW1-d2; 2-way-ANOVA** |  |
| Genotype | F (1, 16) = 522.9, P<0.0001 |
| Treatment | F (1, 16) = 522.9, P<0.0001 |
| Genotype X Treatment | F (1, 16) = 522.9, P<0.0001 |
| **HTRF aggregate assay 4C9-tb with MW8-d2; 2-way-ANOVA** |  |
| Genotype | F (1, 16) = 413.3, P<0.0001 |
| Treatment | F (1, 16) = 1.176, P=0.2942 |
| Genotype X Treatment | F (1, 16) = 1.204, P=0.2888 |

**Supplementary Table 5:** snRNA-seq extended data. Including cell cluster module score, single cell DE output & pathways, pathway files, heatmaps selected genes, cell maker lists, RStudio package versions, and the list of potential off-target genes and differentially expressed genes in those potential off-targets. See Excel sheets attached.

**Supplementary Table 6:** snRNA-seq extended data of pseudo-bulk DE output. Including pseudo-bulk DE output of all cell clusters.

**Supplementary Table 7:**
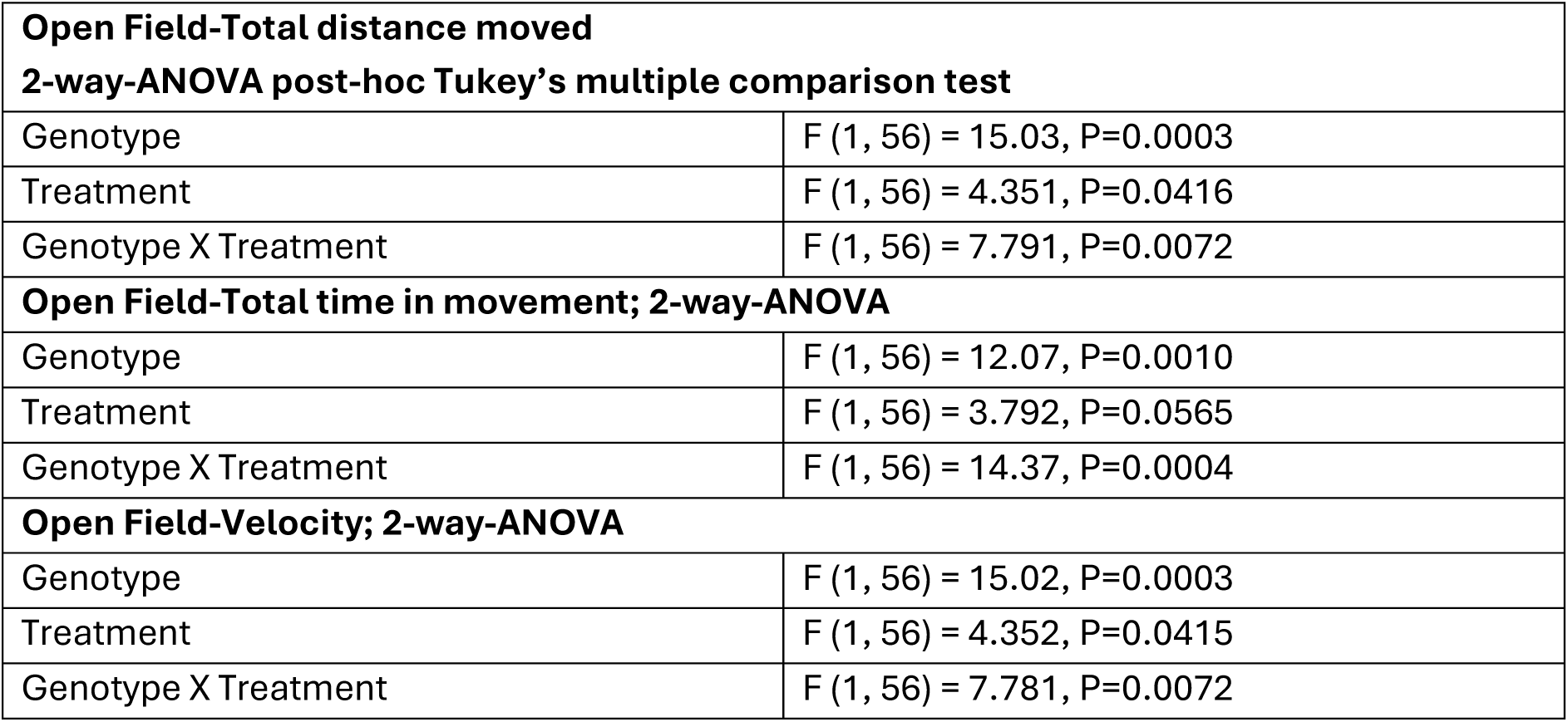
statistics of behavioral tests figure 5.

**Supplementary Table 8:** statistics of behavioral tests Supplementary Fig. 18.

|  |  |
| --- | --- |
| **Balance beam; 2-way-ANOVA post-hoc Tukey's multiple comparison test** |  |
| Genotype | F (1, 56) = 31.34, P<0.0001 |
| Treatment | F (1, 56) = 3.198, P=0.0791 |
| Genotype X Treatment | F (1, 56) = 3.375, P=0.0715 |
| **Accelerated rotarod-latency to fall; 2-way-ANOVA** |  |
| Genotype | F (1, 56) = 21.05, P<0.0001 |
| Treatment | F (1, 56) = 6.043, P=0.0171 |
| Genotype X Treatment | F (1, 56) = 0.01428, P=0.9053 |
| **Fixed rotarod-12rpm latency to fall; 2-way-ANOVA** |  |
| Genotype | F (1, 56) = 19.82, P<0.0001 |
| Treatment | F (1, 56) = 0.2038, P=0.6534 |
| Genotype X Treatment | F (1, 56) = 1.351, P=0.2501 |
| **Fixed rotarod-12rpm total falls in 60 seconds; 2-way-ANOVA** |  |
| Genotype | F (1, 56) = 33.01, P<0.0001 |
| Treatment | F (1, 56) = 2.979, P=0.0899 |
| Genotype X Treatment | F (1, 56) = 1.282, P=0.2624 |
| **Fixed rotarod-24rpm latency to fall; 2-way-ANOVA** |  |
| Genotype | F (1, 56) = 34.25, P<0.0001 |
| Treatment | F (1, 56) = 1.891, P=0.1746 |
| Genotype X Treatment | F (1, 56) = 0.3699, P=0.5455 |
| **Fixed rotarod-24rpm total falls in 60 seconds; 2-way-ANOVA** |  |
| Genotype | F (1, 56) = 20.00, P<0.0001 |
| Treatment | F (1, 56) = 5.205, P=0.0264 |
| Genotype X Treatment | F (1, 56) = 1.018, P=0.3173 |

**Supplementary Table 9:**
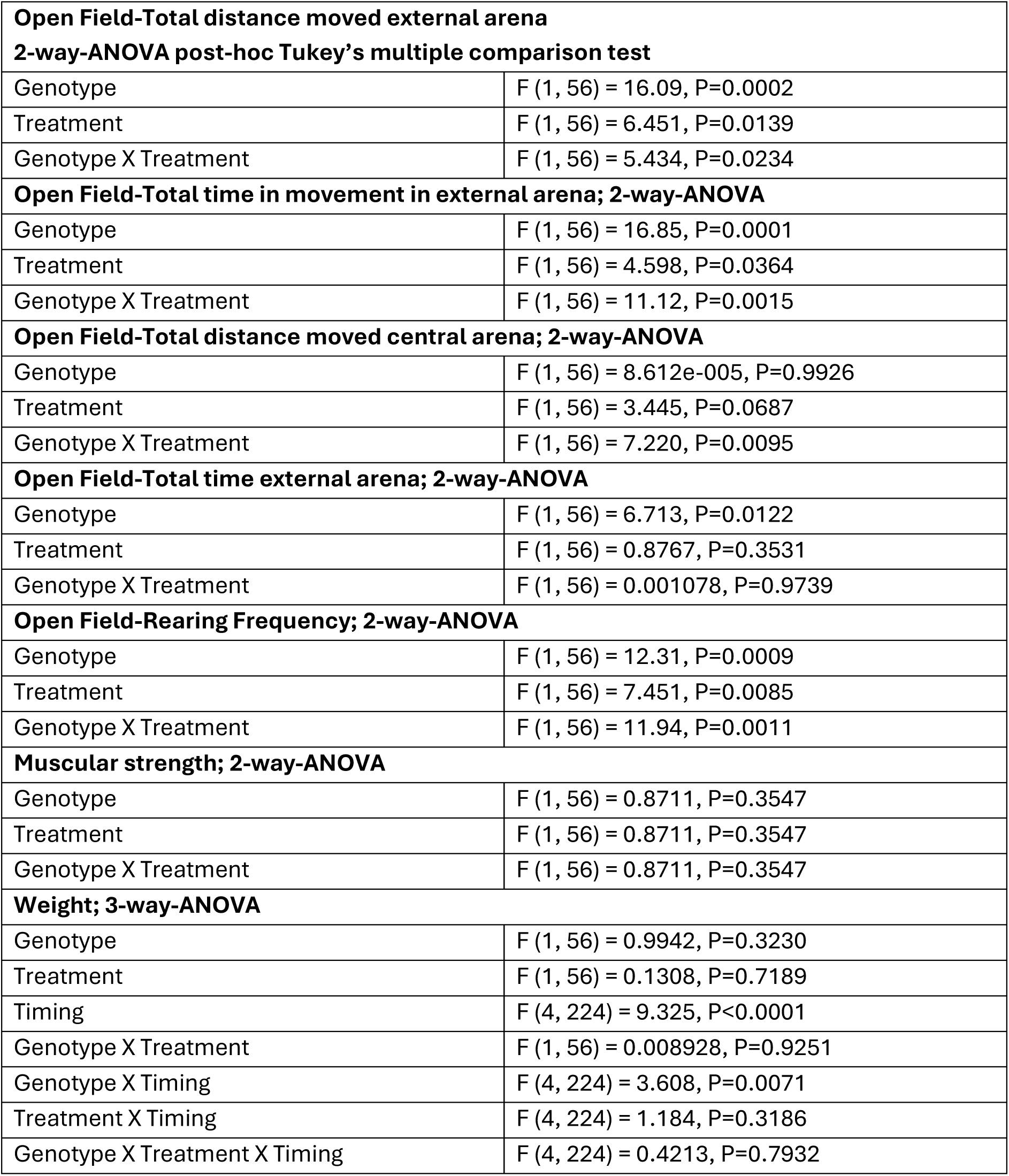
statistics of behavioral tests Supplementary Fig. 19.

**Supplementary Table 10:**
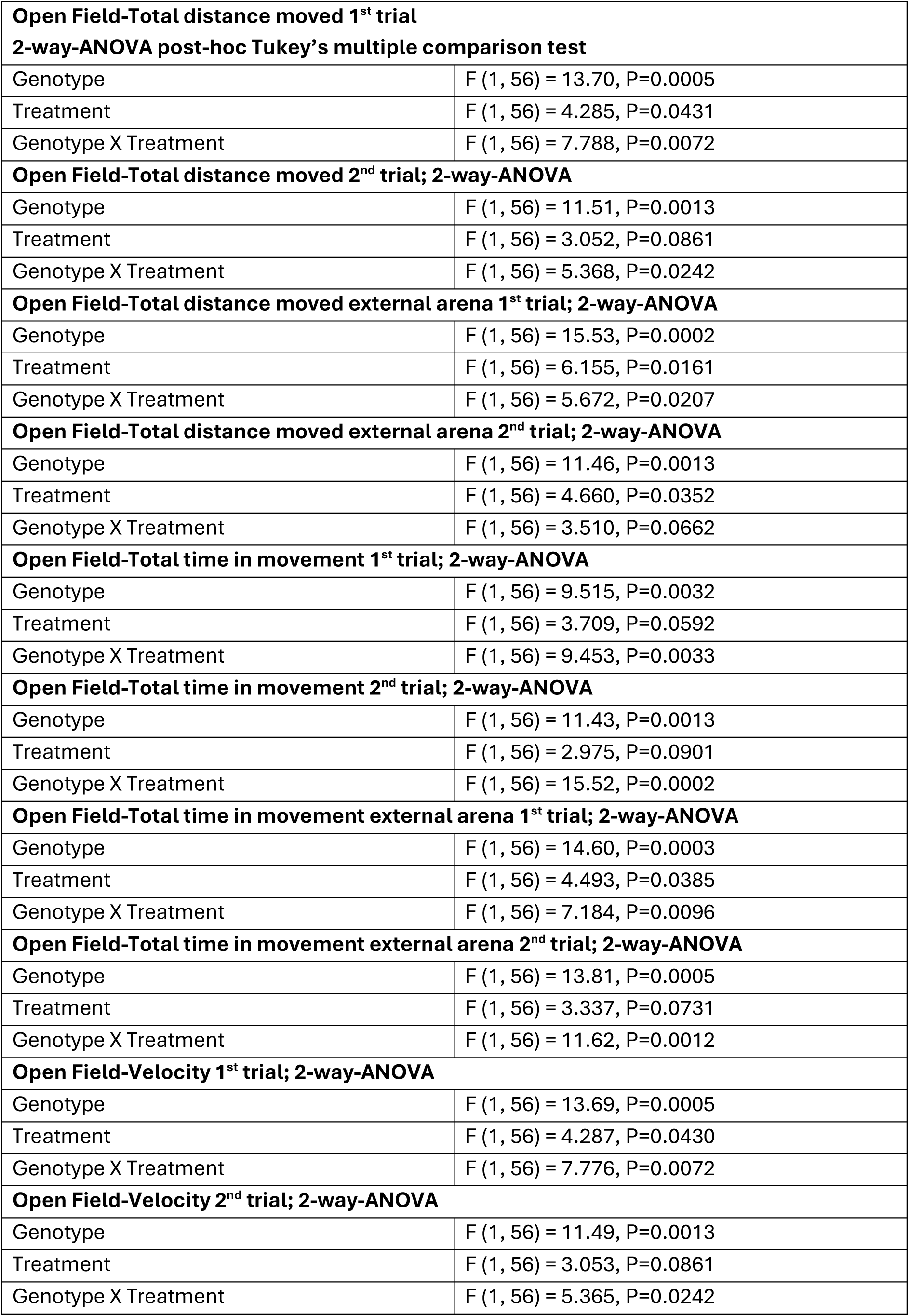
statistics of behavioral tests Supplementary Fig. 20.

**Supplementary Table 11:** Antibodies used in this study.

| Antibody | Reference | Species | Clonality | Method and concentration |
| --- | --- | --- | --- | --- |
| Anti-HA-Tag (C29F4) | Cell Signalling-3724T | Rabbit | Monoclonal | Immunofluorescence (1:500) |
| Anti- Anti-huntingtin Hybridoma MW8 | 11719267<br>11792860 | Mouse | Monoclonal | Immunofluorescence (1:1000) |
| Anti-S100 beta antibody [EP1576Y] - Astrocyte Marker | ab52642 | Rabbit | Monoclonal | Immunofluorescence (1:100) |
| Anti-Beta III Tubulin, Cy3 Conjugate Antibody | AB15708C3 | Rabbit | Polyclonal | Immunofluorescence (1:1000) |
| Anti-CRISPR/Cas9 monoclonal antibody 4G10 | C15200216-100 | Mouse | Monoclonal | Western blotting (1:1000) |
| Anti-β-Actin | A5441 | Mouse | Monoclonal | Western blotting (1:2000) |
| Anti-Green Fluorescent Protein | MAB3580 | Mouse | Monoclonal | Western blotting (1:1000) |
| Anti-Green Fluorescent Protein | ZRB1097 | Rabbit | Monoclonal | Immunofluorescence (1:1000) |
| Anti-NeuN clone A60 | MAB377 | Mouse | Monoclonal | Immunofluorescence (1:500) |
| Anti-iba1 | 09-19741 | Rabbit | Polyclonal | Immunofluorescence (1:2000) |
| Anti-GFAP | MAB360 | Mouse | Monoclonal | Immunofluorescence<br>(1:1000) |
| Anti-Green<br>Fluorescent Protein | AB16901 | Chicken | Polyclonal | Immunofluorescence<br>(1:1000) |
| Anti-DARPP-32 | 2306T | Rabbit | Monoclonal | Immunofluorescence<br>(1:500) |
| Goat anti-Rabbit IgG<br>(H+L) Cross-Adsorbed<br>Secondary Antibody,<br>Alexa Fluor™ 488 | A11008 | Goat | Polyclonal | Immunofluorescence<br>(1:1000) |
| Goat anti-Mouse IgG<br>(H+L) Highly Cross-Adsorbed Secondary<br>Antibody, Alexa<br>Fluor™ 680 | A21058 | Goat | Polyclonal | Western blotting<br>(1:1000) |
| Anti-β-Tubulin III<br>antibody clone<br>SDL.3D10 | T8660 | Mouse | Monoclonal | FACS<br>(1:500) |
| Anti-human CD44<br>Antibody, FITC<br>Conjugate Antibody | 130-113-896 | Mouse | Monoclonal | FACS<br>(1:500) |
| Goat anti-Mouse IgG<br>(H+L) Highly Cross-Adsorbed Secondary<br>Antibody, Alexa<br>Fluor™ 568 | A-11031 | Goat | Polyclonal | FACS<br>(1:1000) |
| Anti-HTT aa 1-17 2B7 | CHDI<br>Foundation | Mouse | Monoclonal | HTRF (1ng well<sup>-1</sup>) |
| Anti-polyQ MW1 | CHDI<br>Foundation | Mouse | Monoclonal | HTRF (40ng well<sup>-1</sup>) |
| Anti-HTT aa 51-71<br>4C9 | CHDI<br>Foundation | Mouse | Monoclonal | HTRF (1ng well<sup>-1</sup>) |
| Anti-HTT end at<br>proline MW8 | CHDI<br>Foundation | Mouse | Monoclonal | HTRF (40ng well<sup>-1</sup>) |

**Supplementary Table 12:**
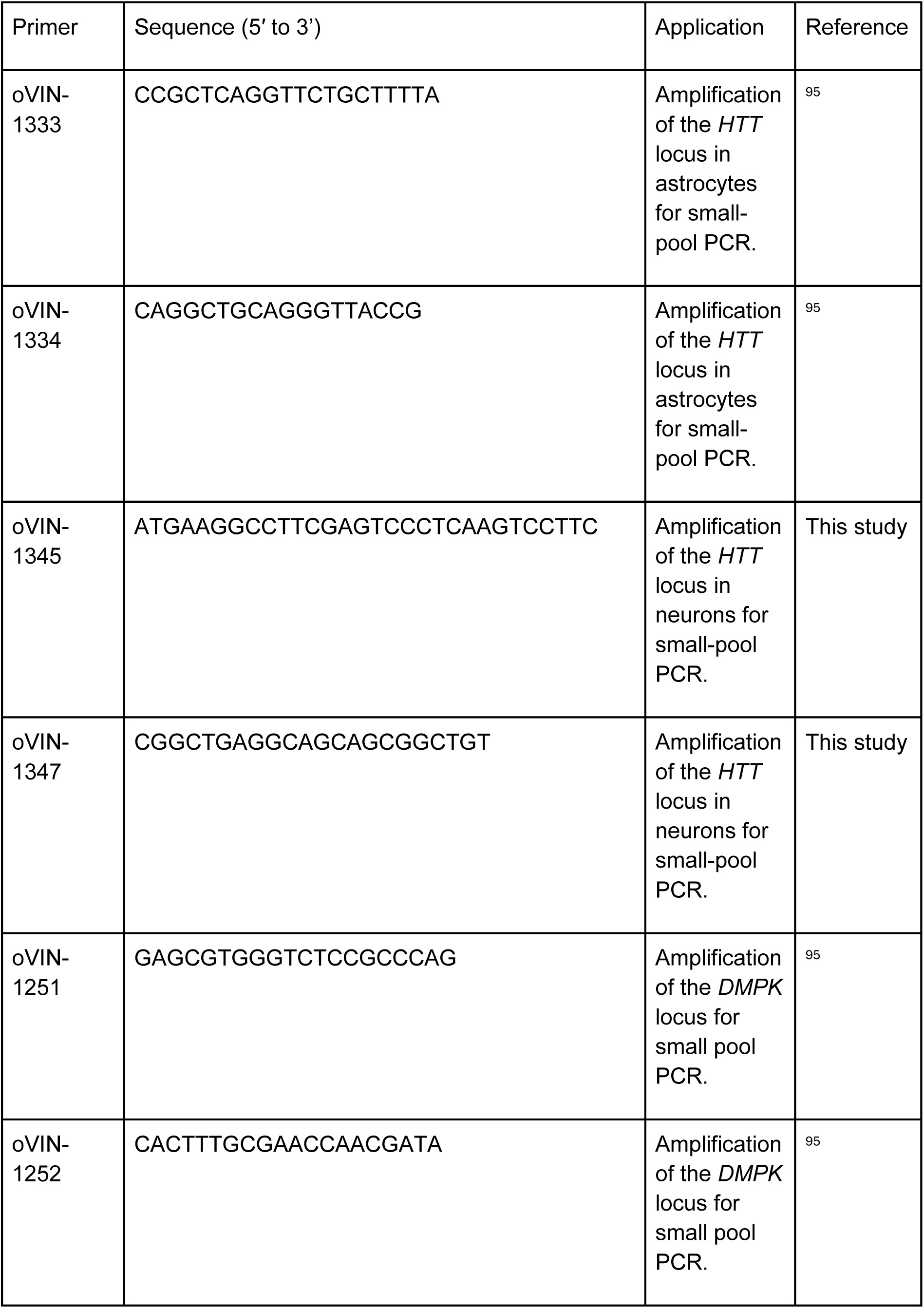
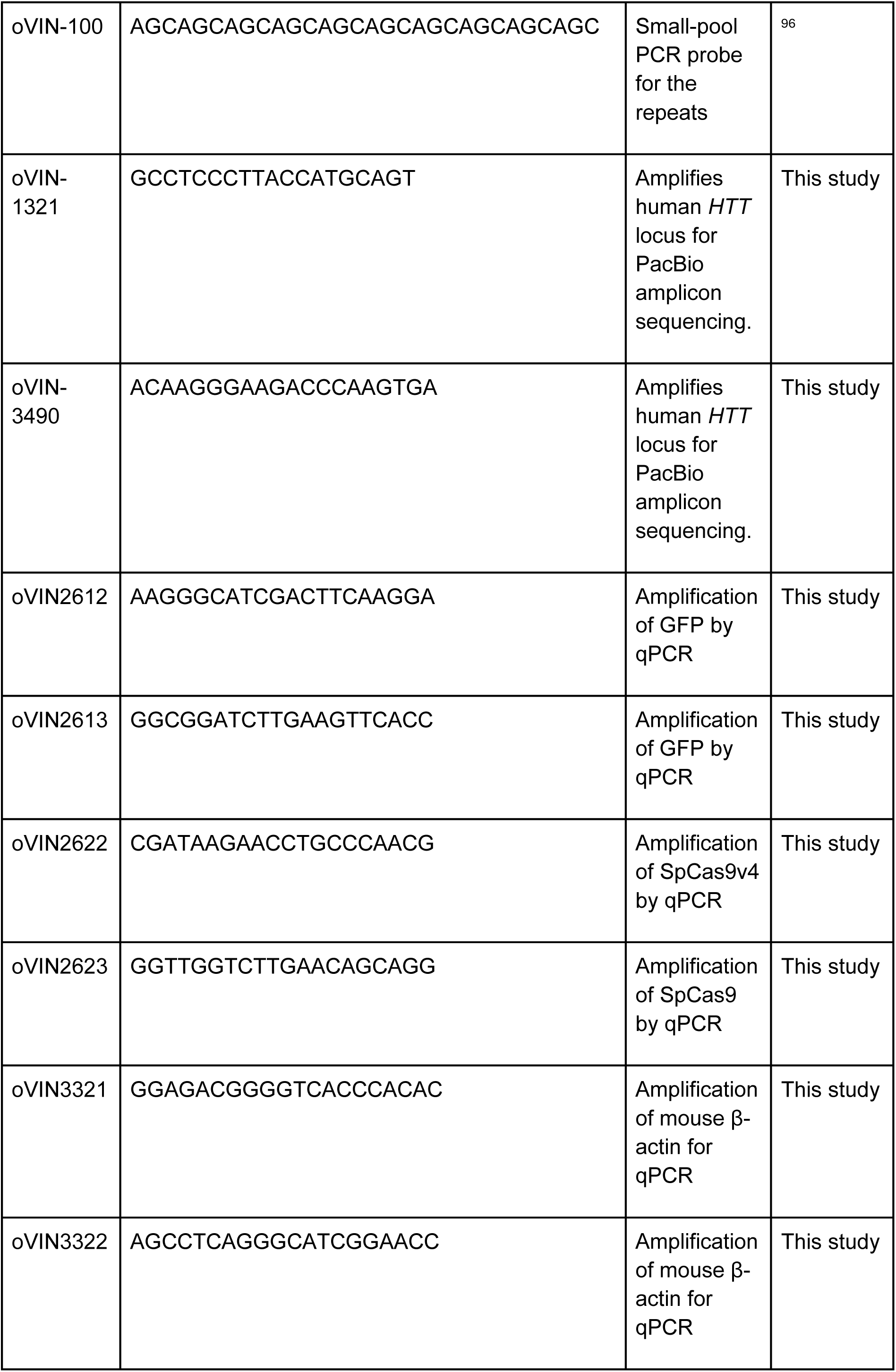
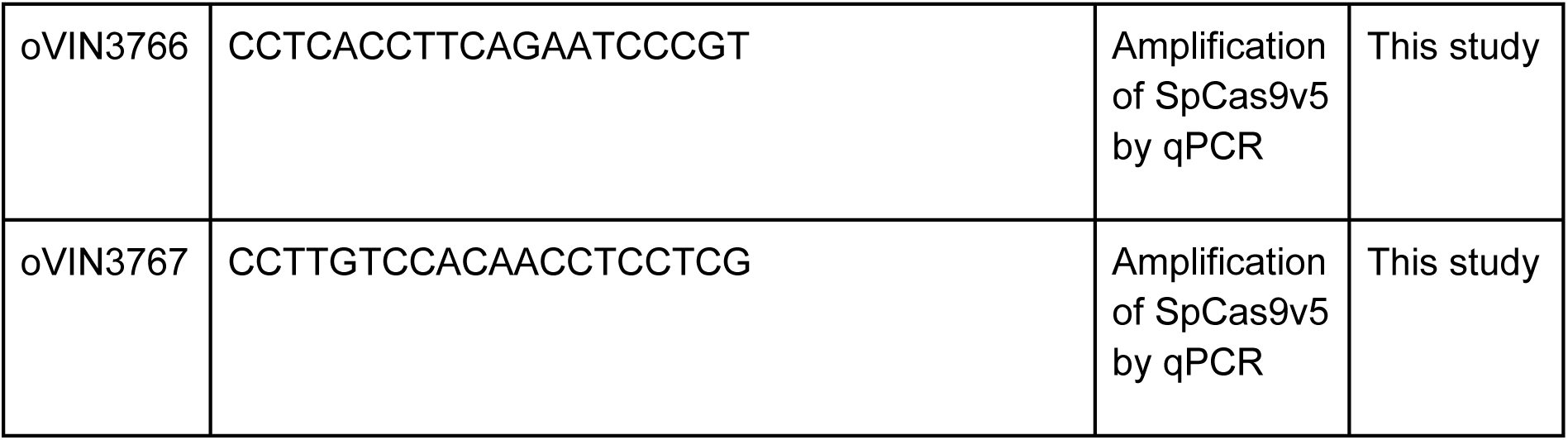
Primers used herein.

